# The taxonomics of the diverse, lithe basal eutyrannosaur genera and species of late Maastrichtian western North America

**DOI:** 10.64898/2025.12.10.693447

**Authors:** Franco Sancarlo, Gregory S. Paul

## Abstract

In 1946 Gilmore observed that the Hell Creek Formation component of the late Maastrichtian TT-zone contained more tyrannosaur taxa than just titanic *Tyrannosaurus rex*, based on a small, ontogenetically mature skull with a higher tooth count than the tyrant lizard. Over four decades later the skull was assigned to its own genus, *Nanotyrannus lancensis*. Also tagged shortly after was the taxon *Stygivenator molnari*. At the last turn of the century it was contended that all TT-zone tyrannosaurs are adult or juvenile *T. rex*, a view that became accepted by many. Because of the unusual morphological alterations that this hypothesis requires, it was consequently proposed that during ontogeny *T. rex* experienced a peculiar fish like metamorphosis. This conservative analysis rejects radical ontogenetic speculations, in favour of *Tyrannosaurus* having shared much the same average amniote/diapsid conventional growth observed in its fellow tyrannosaurids and especially tyrannosaurins. Actual juvenile *Tyrannosaurus s*pecimens show the same tooth count and other osteological features observed in the adults, negating the need for the metamorphosis not observed in any amniotes. This option has become widely accepted. The most extensive phylogenetic analysis to date confirms that the comparative anatomical analysis that most TT-zone mini tyrannosaurs possess a number of features that indicate they are not only not *Tyrannosaurus*, but represent an array of basal eutyrannosaur taxa that are neither *Nanotyrannus* nor *Stygivenato*r, with high tooth counts, and at least in some examples elongated hands, prominent dentary grooves and other features. Two new taxa are correspondingly named at the species and/or genus level, *Gilmoretyrannus lethaeus* and *Larsonvenator elegans*. These are postulated to represent at least in part the migration of basal eutyrannosaurs from former Appalachia into the Laramidia region via the new Laralachia land bridge. The total number of tyrannosaur taxa now known to have inhabited the TT-zone for over 1 MA is seven species in five genera from multiple subfamilies, with a smaller portion of this collection extant at any given stratigraphic level.

## Introduction

The following is a brief overview of publications on the subject focusing on highlights pertinent to this analysis. Found in the latest Maastrichtian Hell Creek Formation, the gigantic avepod theropod *Tyrannosaurus rex*, then represented mainly by two semi-complete skulls and skeletons, was recognized after the prior turn of the century (Osborn 1905, 1906). During World War 2 another, much smaller tyrannosaur skull with a higher number of maxillary teeth was found in the same formation (Fig. 1A, 2H). Using standard comparative anatomy, Gilmore (1946) recognized that CMNH 7541 was a distinct taxon that he assigned to *Gorgosaurus lancensis*, establishing the multiple tyrannosaur taxa hypothesis (MTTH) for TT-zone (sensu Paul *et al*., 2022, Paul 2025a) tyrannosaurs, a view supported by Paul (1988). The same year Bakker *et al*. (1988) described CMNH 7541 as a new genus, *Nanotyrannus lancensis,* Larson (2008) concurred with the validity of the taxon. A detailed redescription of CMNH 7541 was skeptical of it being a *Tyrannosaurus* juvenile (Witmer & Ridgely 2010), in part because is tooth count is too high for it to be in the latter genus. A broadly similar sized partial cranium from the Hell Creek (Fig. 2F; Molnar 1978), LACM 28471, was rendered *Aublysodon molnari* (Paul 1988; supported by Molnar & Carpenter 1989) and then *Stygivenator molnari* (Olshevsky *et al*. 1995) – these were the first works to further increase the apparent diversity of TT-zone lesser sized (femur ∼900 mm or less, ∼ 2 tonnes or less) tyrannosaurs. Currie (2003a) continued to support the MTTH. Larson (2013a) argued that fairly complete, slender proportioned, many toothed BRMP 2002.4.1 (Figs. 1B, 2G, 3D) is a *Nanotyrannus lancensis* also of about the same size. That year Larson (2013b) critically demonstrated that new Hell Creek and Lance Formation specimens, including the virtually complete, many toothed skull and skeleton NCSM 40000 of parallel dimensions to the others (Figs. 1C, 3C, 6C), sport a manus as a long or longer than those of adult *Tyrannosaurus* (Fig. 4D,E), further effectively precluding their being juveniles of the latter. Schmerge & Rothschild (2016a,b) supported the placement of CMNH 7541 and BRMP 2002.4.1 outside *Tyrannosaurus* due to their possession of a prominent lateral dentary groove absent from adult and juveniles of the tyrant lizard (Fig. 2). Jevnikar & Zanno (2021) observed that the growth of BRMP 2006.4.4 does not conform with it being a juvenile *Tyrannosaurus*, and Griffin *et al*. (2025) finding that CMNH 7541 is an adult. Extensive analyses by Longrich & Saitta (2024) and Paul (2025a, also 2024b) detailed reasons that most small-bodied TT-zone tyrannosaurs are not juvenile *Tyrannosaurus*, while a minority are. Those studies further were the first to find that the non*Tyrannosaurus* specimens preserve a taxonomically diverse set of nontyrannosaurid baso-eutyrannosaurs (eutyrannosaurs with or likely to have had long forelimbs, excluding tyrannosaurids [and aliormanians], modified from Paul 2025a), and indicated some were outside *Nanotyrannus*. Paul (2024b, and formally in 2025a) was the first to diagnose NCSM 40000 outside that genus, and as a possible new genus. Paul (2024b, 2025a) further proposed that the exceptional diversity of TT-zone tyrannosaurs was due at least in part to a paleogeographically novel influx of long-handed baso-eutyrannosaurs from eastern North America over the just emerged Laralachia bridge that had reunited the long split continent. Later in 2025 Zanno & Napoli further demonstrated that CMNH 7541, BRMP 2002.4.1, NCMS 40000 and other specimens are basal eutyrannosaurs, concentrated in species of *Nanotyrannus* whose predecessors probably moved in from Appalachia. The results of Raun *et al*. (2025a) imply that the growth of *Tarbosaurus* favors the MTTH. That more than one tyrannosaur taxon can share the same habitat, sometimes both gigantic, other cases very disparate in size, has been established (Russell 1970; Currie 2003a; Kurzanov 1976; Brusatte *et al*. 2009; Paul 2010, 2016, 2024a,b; Lu *et al*. 2014; Mo & Xing 2015; Brusatte & Carr 2016; Carr 2022; Zheng *et al*. 2024). Paul (2016, 2024a,b, 2025a,b), Paul *et al*. (2022) and Longrich & Saitta (2024 tentatively) split mature *Tyrannosaurus rex* itself into multiple intragenera sibling species, adding *T. imperator* and *T. regina* (compare wider variation in *Tyrannosaurus* skulls compared to in other specious tyrannosaurid genera, Figs. 5, 7). Zanno & Napoli (2025) offered that the basic premise may be correct. That gigantic *Tarbosaurus* appears to contain sibling species (Raun *et al*. 2025b) adds support for the MTTH.

**FIGURE 1.**
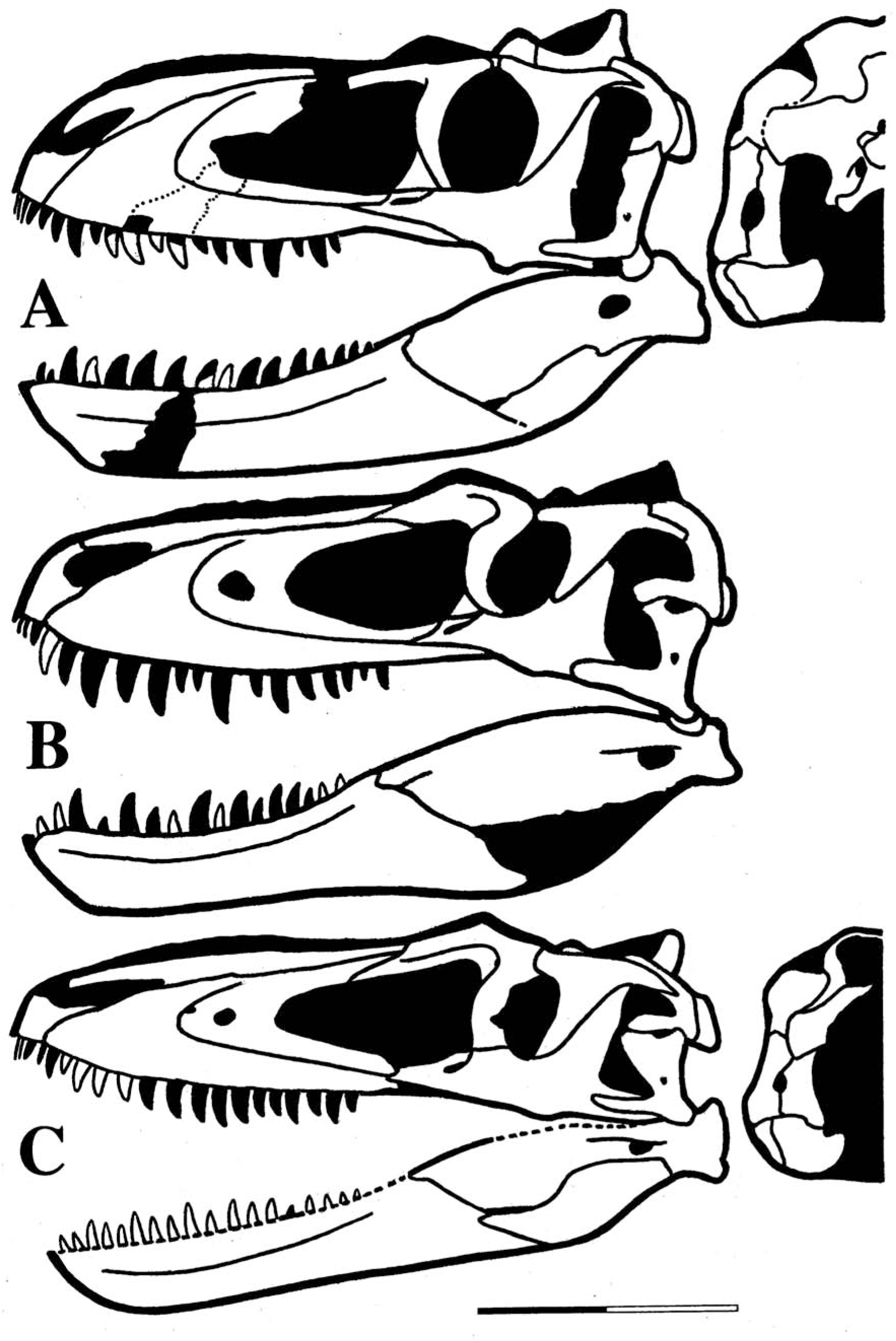
TT-zone known bone baso-eutyrannosaur holotype crania in lateral and posterior views to approximate same scale, bar equals 250 mm. **A** *Nanotyrannus lancensis* CMNH 7541 holotype (adult, 650 mm, lacrimal hornlet and anterior rami of squamosal-quadratojugal not preserved, area of breakage of mid maxillae approximated by dotted lines); **B** *Gilmoretyrannus lethaeus* BMRP 2002.4.1 holotype (subadult, 685); **C** *Larsonvenator elegans* NCSM 40000 holotype (near adult, 660).

**FIGURE 2.**
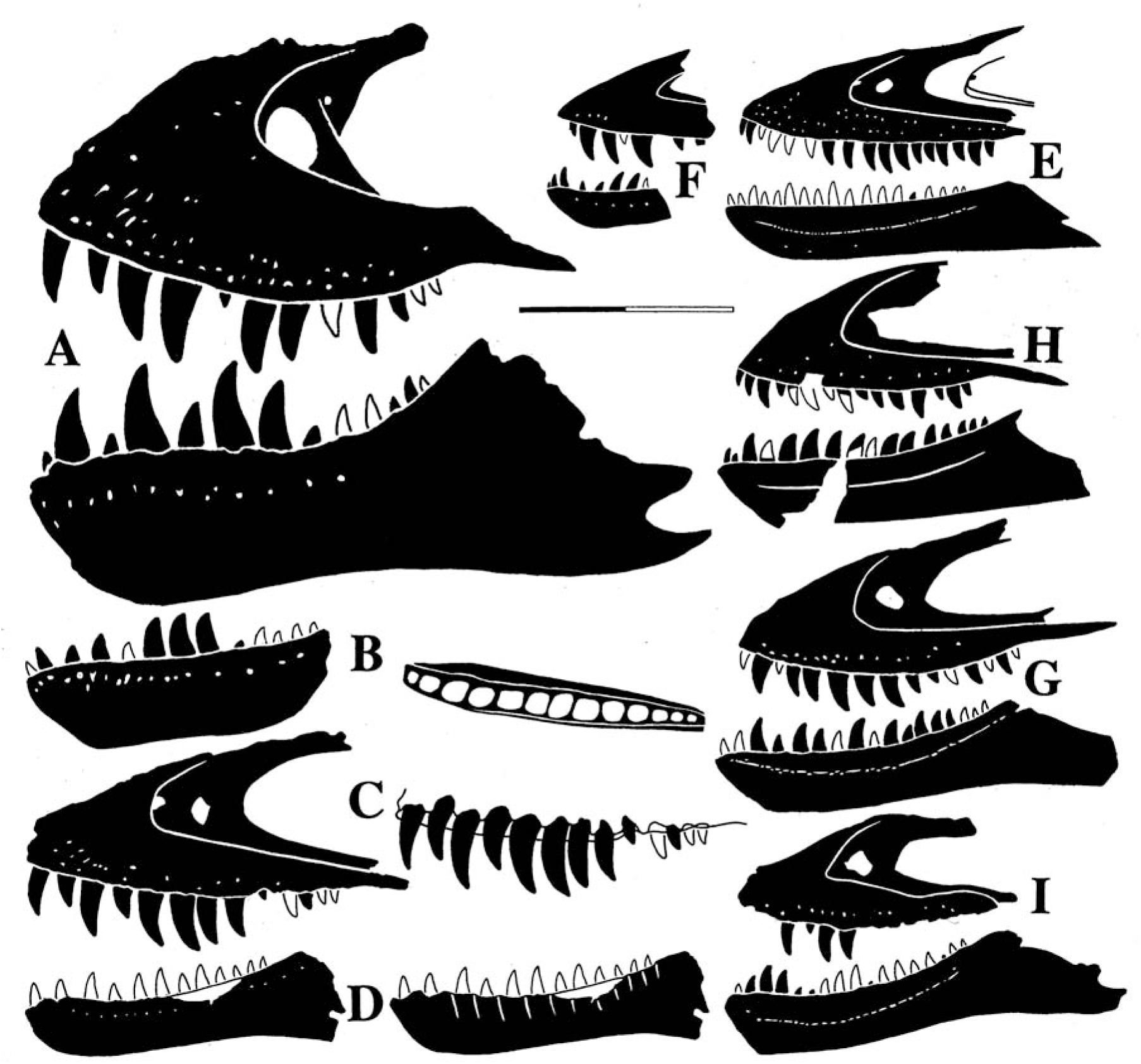
TT-zone eutyrannosaur known-bone maxillae and dentaries to same scale, bar equals 200 mm, dimensions for E are approximate (derived in part from Fig. 8 in Paul 2025a with additions and/or alterations to **B** and **D**). Maxilla and/or dentary in left view (some reversed, some combined from both sides), known tooth positions lacking teeth or not visible by eye or published scans indicated by tooth outlines: **A**, *Tyrannosaurus rex* CMNH 9380 holotype (adult, 6.5 tonnes); **B** *Tyrannosaurus incertae sedis* BHI 6439 (juvenile, ∼1, dorsal view of alveoli); **C** *Tyrannosaurus imperator*? KU 156375 (juvenile, ∼800 kg); **D** *Tyrannosaurus incertae sedis* Baby Bob (juvenile, ∼500, medial view of tooth sulci). Baso-eutyrannosaurs: **E** *Larsonvenator elegans* NCSM 40000 holotype (near adult, ∼600); **F** *Stygivenator molnari* LACM 28471 holotype (subadult? ∼400); **G** *Gilmoretyrannus* BMRP 2002.4.1 holotype (subadult, 570); **H** *Nanotyrannus lancensis* holotype CMNH 7541 (adult, ∼450); **I** *Gilmoretyrannus incertae sedis*, HRS 08 (subadult, ∼450).

**FIGURE 3.**
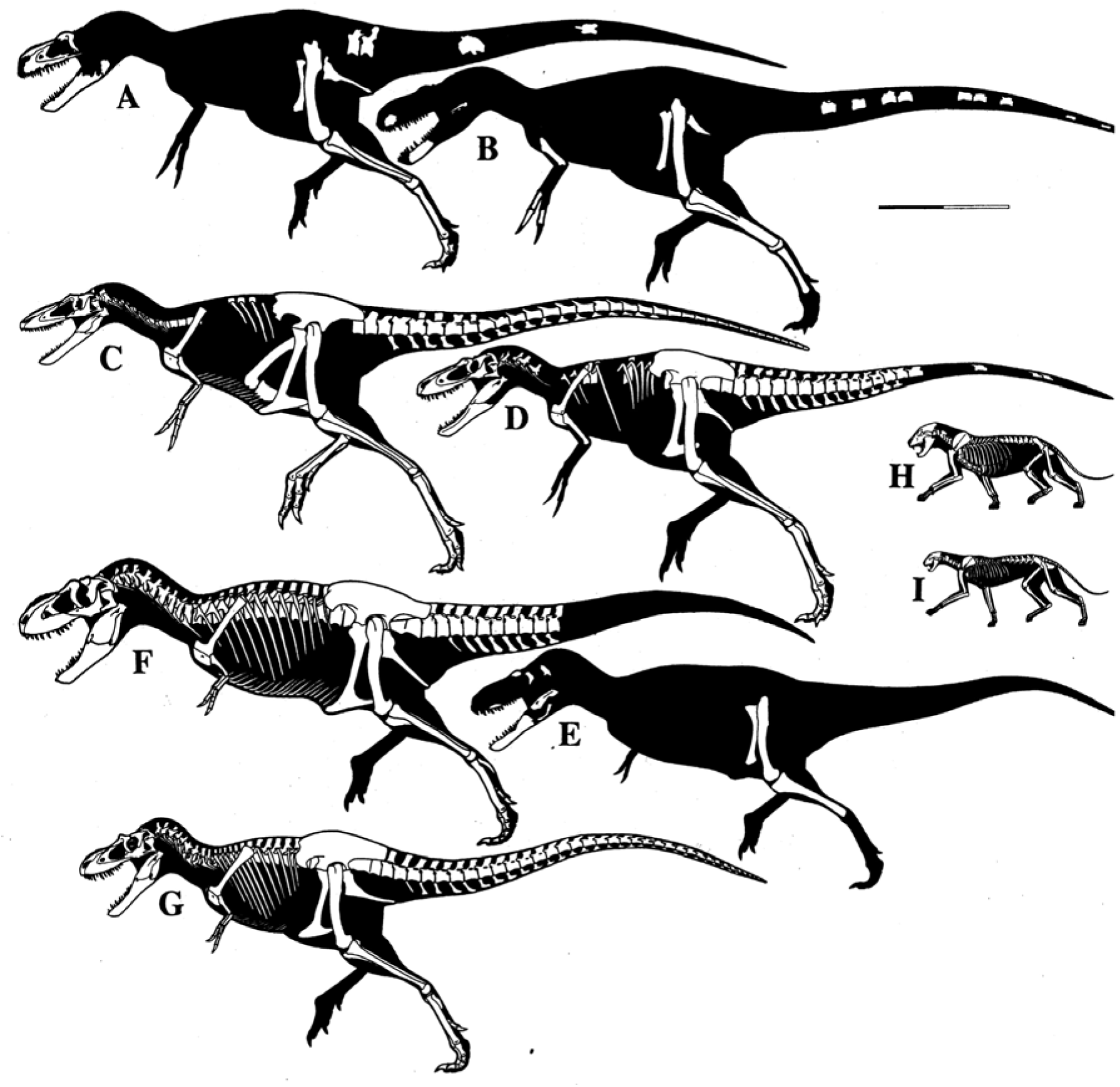
Known/visible-bone profile-skeletals of subjects to same scale, bar equals 1 m. **A** *Appalachiosaurus montgomeriensis* holotype RMM 6670 (femur length 786 mm, ∼700 kg); **B** *Dryptosaurus aquilunguis* holotype ANSP 9995 (780, ∼700); **C** *Larsonvenator elegans* holotype NCSM 40000 (near adult, 717, ∼600); **D** *Gilmoretyrannus lethaeus* holotype BMRP 2002.4.1 (subadult, ∼720, ∼570) with lower forelimb proportions extrapolated from those of other baso-eutyrannosaurs with similarly long humeri; **E** *Tyrannosaurus incertae sedis* Baby Bob (juvenile, 645, ∼670); **F** *Tarbosaurus* MgD-1/3 (juvenile, 700 mm, 850); **G** *Gorgosaurus libratus* TMP 1991.36.5000 (juvenile, 645, 590); **H** pentherine felid *Panthera* (immature, 30); **I** feline felid *Acinonyx* (immature, 20).

**FIGURE 4.**
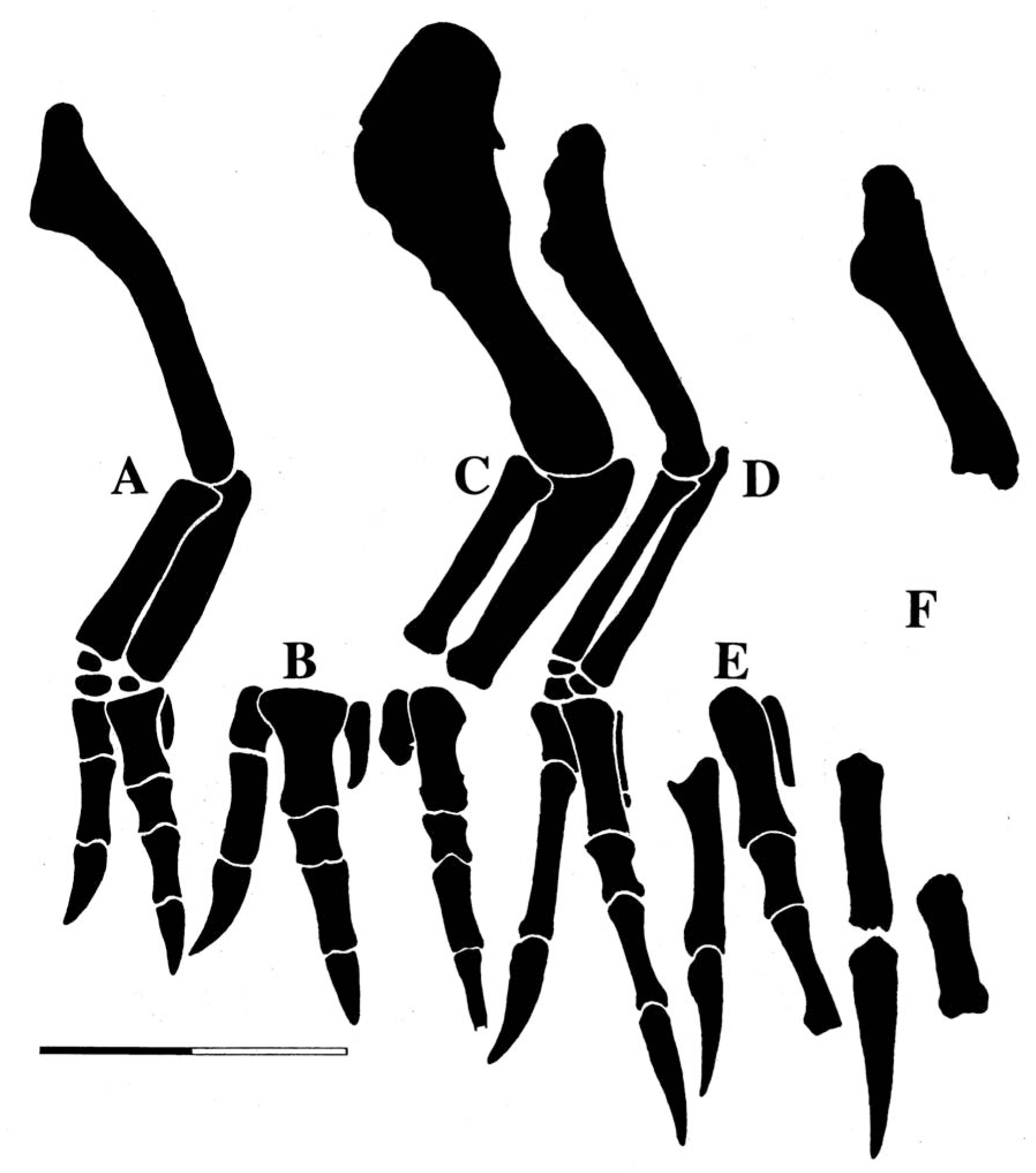
TT-zone known-bone tyrannosaurin and baso-eutyrannosaur forelimbs to same scale, bar equals 250 mm (modified from Fig. 9 in Paul 2025a except for **D**). **A** *Tyrannosaurus incertae sedis*, UCRC V1 (subadult); **B** *Tyrannosaurus regina*, MOR 980 (adult); **C** *Tyrannosaurus imperator* holotype FMNH PR 2081 (adult; placement of distal elements not certain); **D** *Larsonvenator elegans* holotype NCSM 40000 (subadult); **E** *Larsonvenator elegans* (near adult), Jodi (adult?); **F** *Dryptosaurus aquilunguis* holotype ANSP 9995 (placement of one distal element not certain).

**FIGURE 5.**
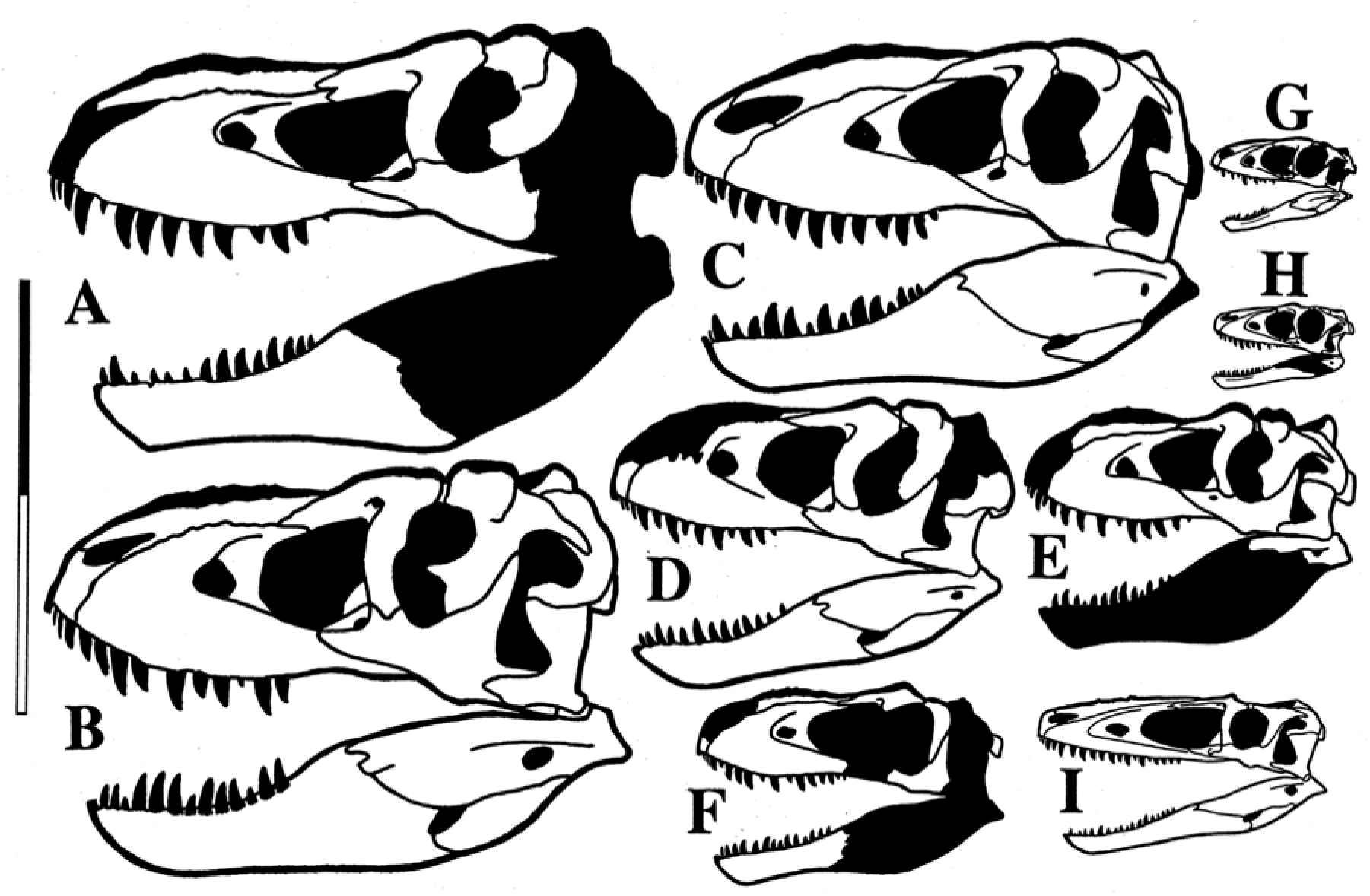
Known-bone Nemegt complex tyrannosaur crania to same scale, bar equals 1 m (from Fig. 2 in Paul 2025a except for **I** which is from Paul, 2024a,b). Some skulls reversed, and/or both sides used to complete restoration. *Tarbosaurus* ontogenetic series: **A** PIN 551-1 (mature, ∼4.5 t); **B** MgD-1/4 (mature, ∼3.5); **C** PIN 551-3 (immature, 2.5); **D** PIN 553-1 (juvenile. ∼1.5); **E** MgD-1/3 (juvenile, 850 kg); **F** PIN 552-2 (juvenile, 600); **G** LHP V18 (*Raptorex kriegsteini* holotype) (juvenile, 75); **H** MPC-D107/7 (juvenile, ∼75); **I** *Alioramus* MPC-D 100/1844 holotype (subadult, ∼250).

**FIGURE 6.**
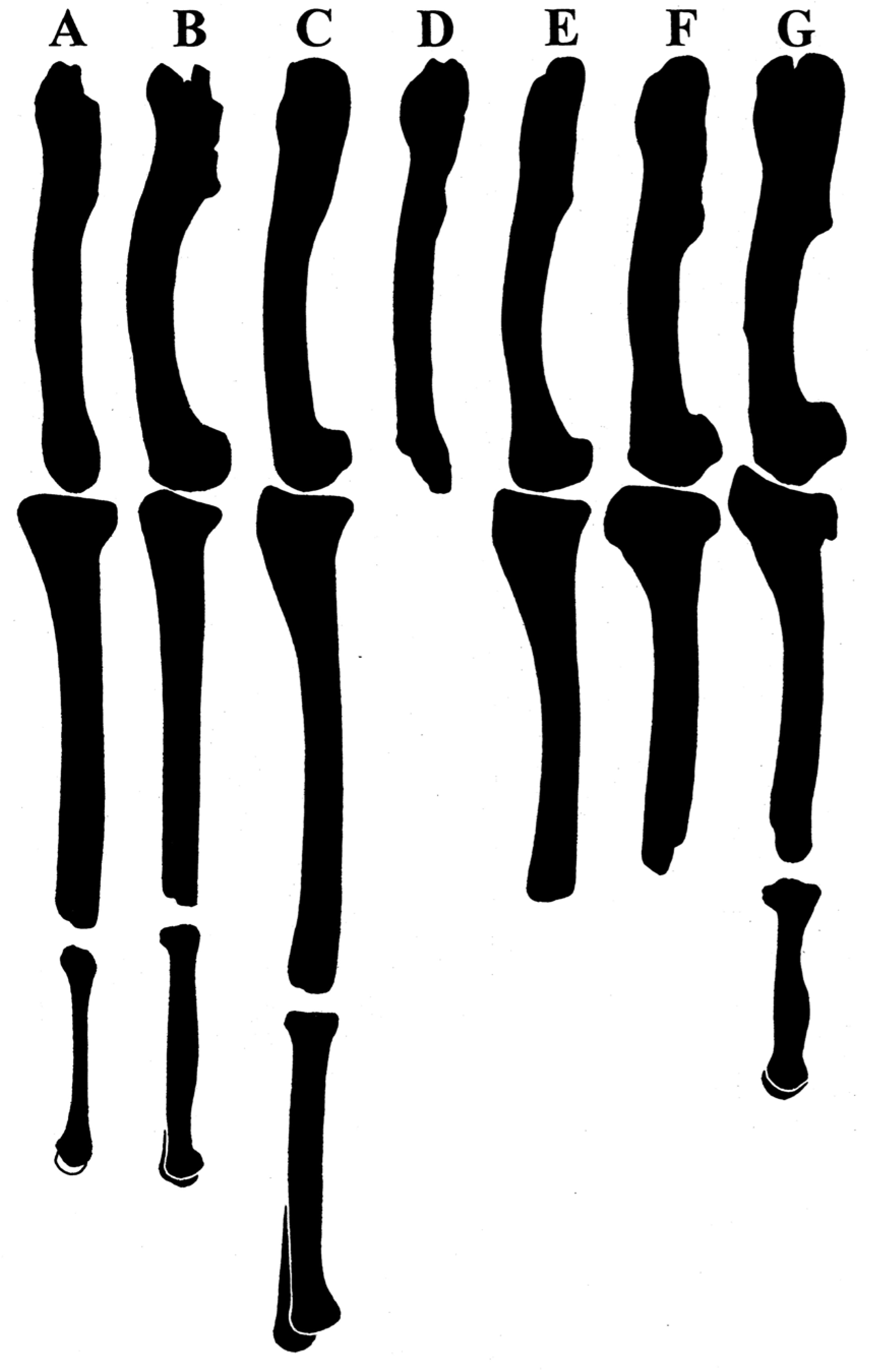
Known-bone profiles of late Campanian to late Maastrichtian Appalachia and TT-zone eutyrannosaur hindlimbs reproduced to same femur length to facilitate visual comparisons of femoral shaft curvatures and element ratios. **A** *Dryptosaurus aquilunguis* holotype ANSP 9995 (femur 780 mm long, tibia 760 mm, metatarsus ∼410); **B** *Appalachiosaurus montgomeriensis* holotype RMM 6670 (786, 762, ∼480); **C** *Larsonvenator elegans* holotype NCSM 40000 (near adult. 717, 820, 560); **D** *Gilmoretyrannus lethaeus*? BMRP 2006.4.4 (774); **E** *Tyrannosaurus incertae sedis* Baby Bob (juvenile, 645. 605); **F** *Tyrannosaurus incertae sedis* USNM 6183 (subadult, 990, 885); **G** *Tyrannosaurus imperator* holotype FMNH PR208 (adult, 1321, 1143, 671).

The fossil growth series of *Tarbosaurus* fossils (Fig. 5A-H; also see Paul 2024a,b skeletal illustrated entries on the taxon) inspired Rozhdestvensky (1965) to casually suggest that CMNH 7541 was a juvenile *Tyrannosaurus* in a one sentence part of a footnote that lacks any analysis. Had the inconsistent tooth counts between the mini tyrannosaur and the tyrant king been contrasted with the steadier values in the terror lizard been noted and used to dismiss *N. lancensis* as a juvenile, then the ETRH may have never arisen. Carpenter (1992) further speculated that *Nanotyrannus* is a juvenile *T. rex*, again without regard to the former’s high number of teeth. In 1999 Carr established the modern everything is *T. rex* hypothesis (ETRH). It was based in part on an incorrect claim of tooth reduction in growing *G. libratus*. Despite that critical error being quickly corrected by Currie 2003a,b), studies focused directly on the ontogenetic segment of the ETRH that support the thesis include Carr & Williamson (2004), Brusatte & Carr (2016), Brusatte *et al*. (2016), Woodward *et al*. (2020), Voris *et al*. (2025), culminating in an extensive study by Carr (2020). Faced with the problems posed by the exceptionally radical ontogenetic changes in *T. rex* required if all the highly anatomically diverse TT-zone lesser tyrannosaur fossils are included in the one taxon, Carr (2020) engaged of the major speculation that the species experienced a sudden ontogenetic metamorphosis unique among amniotes. The recently devised version of the ETRH was often presented as the widely accepted modern paradigm that therefore requires refutation to lose its privileged status, the above citations show that the number of researchers who presented studies that concentrate on the TT-zone data regarding the ETRH versus the MTTH was actually broadly similar. In any case, in the combined data and analysis wake of Longrich and Saitta (2024), Paul (2025a) and then Zanno & Napoli (2025) and Giffin *et al*. (2026), it appears that the full blown ETRH no longer has publicly confirmed absolutist adherents, varying forms of the MTTH now being operative. In reaction, Woodward et al. (2026) takes a relatively neutral stance on the issue. The inherently rigid ETRH had been effectively discouraged research into TT-zone tyrannosaur diversity – which is why the first paper to take a large data set based look at the question of mature *Tyrannosaurus* species was Paul *et al*. (2022) followed by Paul (2025a) -- the MTTH does the opposite. So the ascendency of the latter hypothesis has opened to field to innovative, cutting edge research on the issues with a vigor not see before. It is not just *T. rex* anymore.

Paul (2025a) provisionally placed BMRP 2002.4.1, NCSM 40000 and other TT-zone specimens in previously named genera. So have Zanno and Napoli. (2025), with them considering only *Nanotyrannus* to be viable and sinking *Stygivenator* which was not analyzed separately, while including in the prior the closely related new intragenus sibling species *N. lethaeus*. Initiated by the senior author to test the question, this paper focuses on determining whether or not the mini tyrannosaurs of the TT-zone are all juvenile *T. rex*, if not how many taxa the non*Tyrannosaurus* specimens represent, examine if they are the same taxa as late Cretaceous tyrannosaurs from Appalachia, and if any can be diagnosed on phylogenetic and gradistic grounds sufficient to confirm or require generic and/or specific titles. By doing so, the diversity of the resident fauna can be better determined. Related items involving the origins and diversity of the TT-zone dinosaur predators are considered. This is achieved by presenting and then parsimoniously following the critical aspects of actual, ordinary, observed amniote-diapsid-tyrannosaur-tyrannosaurin ontogeny, rather than venturing into extraordinary speculative hypotheses. This sets the needed conventional standards for addressing the problem moving forward. The largest character matrix phylogenetic analysis to date is conducted (AppendFig. 1). Some aspects of speciation in mature *Tyrannosaurus* are touched upon, but this is not a major subject herein and persons are referred to Paul (2025a) for an extensive examination of that subject.

AMNH, American Museum of Natural History, New York; ANSP, The Academy of Natural Sciences, Philadelphia; BDM, Badlands Dinosaur Museum, Dickinson; BHI, Black Hills Institute, Hill City; BRMP, Burpee Museum of Natural History, Rockford; CMN, Canadian Museum of Nature, Ottawa; CMNH, Cleveland Museum of Natural History, Cleveland; DINO, Dinosaur National Monument, Vernal; DDM, Dinosaur Discovery Museum, Kenosha; DMNS, Denver Museum of Nature and Science, Denver; FMNH, Field Museum of Natural History, Chicago; HRS, Hanson Research Station, Newcastle; KU, Kansas University, Lawrence; LACM, Los Angeles County Museum, Los Angeles; MOR, Museum of the Rockies, Bozeman; MPC, Mongolian Palaeontological Center, Ulaanbaatar; NCSM, North Carolina Museum of Natural Sciences, Raleigh; NHMAD, Natural History Museum Abu Dhabi, Dubai; PIN, Palaeontological Institute, Moscow; RMM, Red Mountain Museum, Birmingham; RSM, Royal Saskatchewan Museum, Regina; TMP, Royal Tyrrell Museum of Palaeontology, Drumheller; UCMP, University of California Museum of Palaeontology, Berkeley; UCRC, University of Chicago Research Collection, Chicago; USNM, National Museum of Natural History, Smithsonian, Washington DC

## Material and methods

### Issues and discourse addressed versus referenced

This analysis is not a comprehensive review of all intricate aspects of the subjects covered. It is intended to address specific issues. As such detailed discussions of many issues that have been addressed in-depth elsewhere of late (as per Paul *et al*. 2022; Longrich and Saitta 2024: Paul 2025a; Zanno & Napoli 2025) are not repeated, the prior analyses being cited when they pertain.

### Phylogenetics

A total of 869 characters and 42 taxa were used in this study, with *Allosaurus* being the outgroup (Appendix 1). Of these characters, 394 are derived from Voris *et al*. (2020, 1–394), with modifications applied to characters 31, 261, 263, and 369; 248 from Dalman *et al*. (2024, 395–642), with a modification applied to character 513; 201 from Zanno & Napoli (2025, 643–843), with modifications applied to characters 763 and 781; and 15 from Larson (2013b, 844–858), while an additional 11 new derived characters are proposed below (859–869, some of which are from Paul *et al*. 2022 and Paul 2025a, including characters assigned to three TT-zone species – some character states are partly overlapping as per other species such of those of *Triceratops* Paul 2025a). A number of the characters are used to help diagnose the systematics of the groups and species. In order to not prejudice their taxonomic placements, CMNH 7541, BM BMRP 2002.4.1 HRS 08, LACM 28471 and NCMS 40000 are scored separately. The analyses were conducted using TNT version 1.6 (Goloboff & Morales 2023) using the New Technology search and Traditional Search (AppendFig. 1).

Due to substantial data gaps in the TT-zone baso-eutyrannosaur fossils the results are somewhat tenuous in terms of their intrarelationships, and relative to the Appalachian baso-eutyrannosaurs. The phylogenetic results need to be supplemented by other aspects of their form and function.

### Critical diagnoses factors

With taxonomy being a combination of clade plus grade, the latter plays a key role in taxonomic determinations when diagnosing genera and species (Mihlbacher 2008; Carpenter 2010; Mader 2010; Knutsen 2012; Carr *et al*. 2017; Chure & Loewen 2020; Johnson *et al*. 2020; Larramendi *et al*. 2020; Paul *et al*. 2022; Danison *et al*. 2024; Sanchez-Fenollosa *et al*. 2024; Paul 2025a,b). Grade differences are the result of the functional differences that help assess higher level taxonomic distances between species. If two species consistently cladistically score and phylogentically position as sibling taxa, that does not automatically mean they are in the same genus. Gradistic criteria expressed as diagnoses also apply, as does the probability of ghost taxa between them due to the gaps in the fossil record. For example, within equids *Plesippus* and *Equus* are close generic relatives to one another (Barrón-Ortiz *et al*. 2019), but are distinguishable from each other at the genus level on gradistic grounds. Within each genus are sibling species that are diagnosable despite being very close relations that in some cases inhabit the same habitats. Similar situations apply in other cases, such as the rhinocerotids *Diceros bicornis* and *Ceratotherium simum* (Liu *et al*. 2021). It is becoming increasingly apparent that intragenera species are more common than had been realized: *Loxodonta* now includes two species (Grubb *et al*. 2000), *Giraffa* four (IUCN 2025), small *Canis* are increasingly specious (Koepfli *et al*. 2025). The probability that this cryptic species factor applies to fossil taxa, tyrannosaurids included, is being increasingly recognized of late (Paul *et al*. 2022; Napoli *et al*. 2023; Longrich & Saitta 2024; Stock *et al*. 2024; Griffin *et al*. 2025; Paul 2025a, Zanno & Napoli 2025; Raun *et al*. 2025b). Within eutyrannosaurs being sibling taxa in mini-clades does not necessarily demand they be a united genus if gradistic differentiation is deemed sufficient, as per *Alioramus* and *Qianzhousaurus*, *Bistahieversor* and *Jinbeisaurus* and, *Albertosaurus* and *Gorgosaurus*, and *Terathoponeus* and *Lythronax* (Append Fig. 1; Brusatte & Carr 2016; Dalman *et al*. 2024; Longrich & Saitta 2024; Voris *et al*. 2025; Zanno & Napoli 2025)

Assessing generic and species status is not just a matter of scoring characters. It is also one of holistic visual comparative anatomy (Paul 2025a). For instance, Voris *et al*. (2025) cited the presence of one attribute on one TT-zone *Tyrannosaurus* dentary, MOR 008, as negating the placement of “*T.” mcraeensis* outside the latter taxon. Aside from the MOR 088 dentary being too damaged to determine the bone’s shape (Paul 2025b), the “*T.” mcraeensis* element is so distinctive in overall form from those of TT-zone fossils (Fig. 4C in Dalman *et al*. 2024) that it is clearly a distinct, probably more basal, taxon. More generally, in this analysis the degree of difference in overall shape and morphological profiles and proportions of individual superficial bones in the adult crania of accepted avetheropod genera, including those that are multispecific (Fig. 7), are compared to the total anatomical configurations of TT-zone non*Tyrannosaurus* mature skulls (Fig. 1). This follows the practical procedure whereby visual anatomical comparisons are used to help distinguish taxa, as per bird watching (Peterson 1934). Likewise, local native animal identification is usually in good accord with Linnean determinations (Paul 2025a and references therein). In standard taxonomic principle gradistic variation within each of a set of genera, especially when closely related, needs to be about the same. *Tyrannosaurus* contains at least as much osteological variability as multispecific *Daspletosaurus* (Fig. 7C-E versus F-J) because the former is at least as specious. Ergo, if differentiation in the TT-zone baso-eutyrannosaur skulls exceeds that observed within the other avetheropod genera, then the former represent multiple genera. If not, then they are not different genera.

**FIGURE 7.**
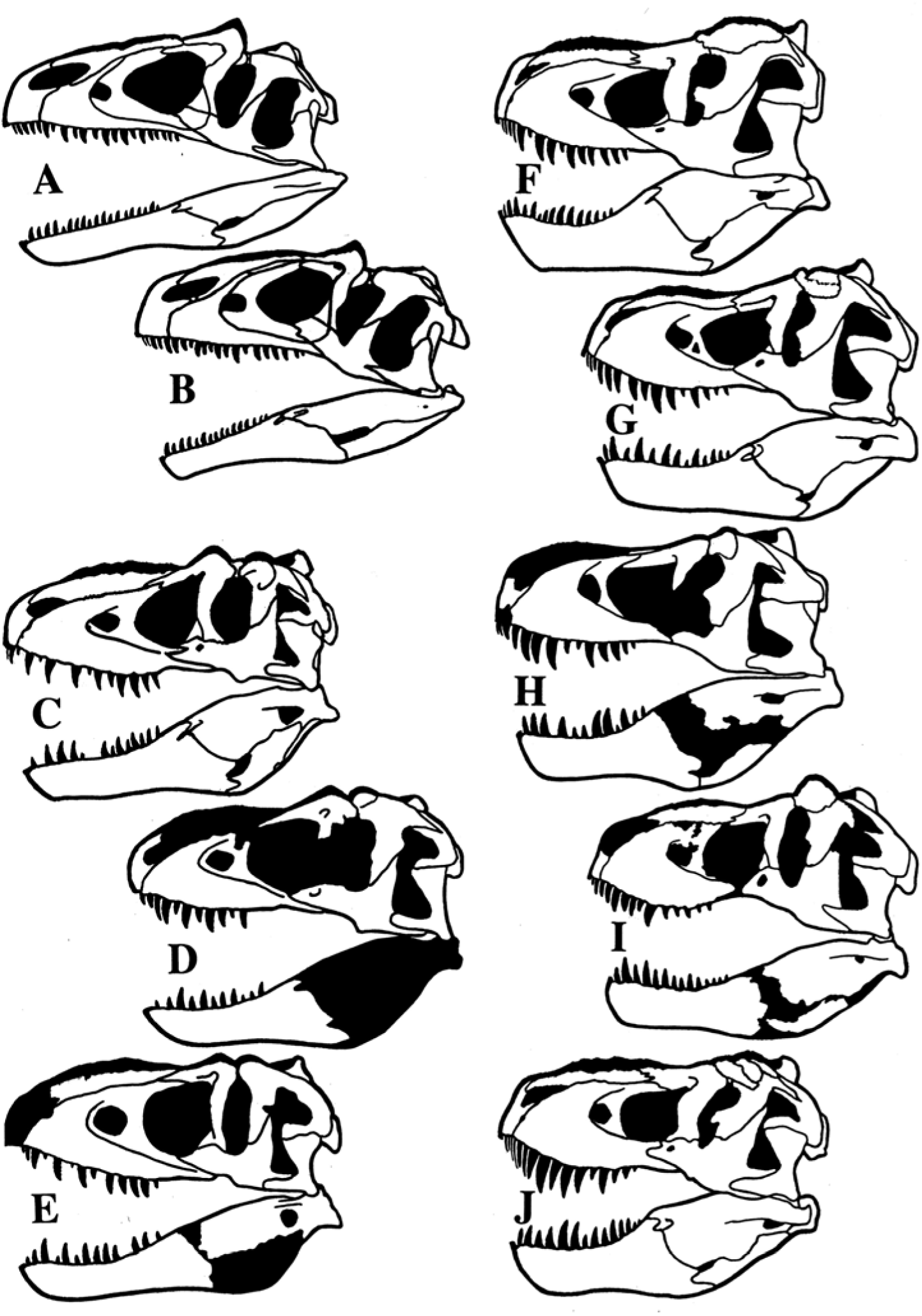
Known-bone avetheropod avepod adult or close to it crania reproduced to same length to facilitate intra and inter genera visual comparisons from Paul (2024b, 2025a, see Fig. 2B,**D**,**E**,**G**,**I** in latter reference for same scale reproductions of **F-J**, as well as additional *Tyrannosaurus* skulls). **A** *Allosaurus jimmadseni* holotype DINO 11541; **B** *Allosaurus fragilis* DINO 2560; **C** *Daspletosaurus torosus* holotype CMN 8506; **D** *Daspletosaurus wilsoni* (or *horneri*) holotype BDM 107; **E** *Daspletosaurus horneri* holotype MOR 590; **F** *Tyrannosaurus imperator* holotype FMNH PR208; **G** *Tyrannosaurus regina* NHMAD 2020.00001; **H** *Tyrannosaurus regina* LACM 150167; **I** *Tyrannosaurus rex* RSM2523.8; **J** *Tyrannosaurus incertae sedis* AMNH 5027.

### The time and niche factors

Evidence has grown that dinosaur species were prone to turning over rapidly, with species typically not lasting more than a few hundreds of thousands of years (Paul 2010, 2016, 2024a,b; 2025a,b; Scannella *et al*. 2014; Fowler 2017; Mallon 2017; Chure and Loewen 2020; Paul *et al*. 2022; Danison *et al*. 2024). The same works also often found that large predators can exhibit considerable diversity at the same level in a particular formation, with the miscellany often increasing with decreasing size. A factor in the fast and diverse evolution of dinosaur taxa may have a genetic basis due to high chromosome numbers that are still present in the species prolific birds (O’Connor *et al*. 2018). The rapid, r-strategy reproduction of giant avepods depositing large numbers of rather small eggs may have also favoured swift evolution and diversification. It therefore presumed herein than if fossils are separated by more than a few hundred thousand years that they are probably different species, unless strong comparative anatomical evidence indicates otherwise. The latter includes near identical morphology of overlapping elements. Nor can it be presumed that incomplete specimens found at the same stratigraphic level of a formation from a given family are those of one species—unless found in very close association without duplications -- rather than being from more than one taxon. Any conclusion one way or another has to be based on the cumulative preponderance of the data. Consisting of multiple species, genera can last over long stratigraphic periods. In the Asian tyrannosaur faunas combining a large or giant taxon with lesser taxa, the latter often appear to constitute one taxon (Brusatte *et al*. 2009; 2016; Brusatte & Carr 2016; Paul, 2010, 2016, 2024a,b). But cryptic small taxa may be present, and two small forms along with a giant appear to be the case in the Nanxiong Formation (Lu *et al*. 2014; Mo & Xu 2015; Zheng *et al*. 2024). It is consequently not assumed that there is only one mini tyrannosaur taxon present in the TT-zone, either in its entirety, or at any particular stratigraphic level. There being multiple small taxa is at least as likely. In western North America the probable presence of a small tyrannosaur taxon was demonstrated some time ago (Lehman & Wick 2012). The possibility that a substantial number of cryptic tyrannosaur species existed in their formations, and are preserved among the fossils collected from the sediments, is very substantial (Paul *et al*. 2022, Longrich and Saitta, 2024; Stock *et al*. 2024; Griffin *et al*. 2025; Paul 2025a; Zanno & Napoli 2025; Raun *et al*. 2025b). The stratigraphic levels of the specimens under consideration when known and not are from Paul (2025a) and Zanno & Napoli (2025).

### Critical emphasis on actual amniote ontogenetic biology

The comparative anatomy is combined with basic ontogenetic norms and principles to produce a parsimonious null hypothesis for the work on TT-zone tyrannosaurs. The degree to which standard realistic ontogenetics is detailed and employed in this study is unprecedented (building upon Longrich & Saitta 2024, Paul 2025a; Zanno & Napoli 2025). This is in contrast to Carr (2020); in order to try to fit all TT-zone specimens of all sizes into *T. rex*, Carr contends that *Tyrannosaurus* underwent a rapid, dramatic metamorphosis that among items included a significant reduction in numbers of teeth via reduction. Among vertebrates, metamorphosis is limited to fish and amphibians, not having been observed among amniotes, tyrannosaurids included (Currie 2003a,b; Longrich & Saitta 2024; Paul 2025a; Yun *et al*. 2025; Zanno & Napoli 2025). Juveniles within a species, and between closely related species, tend to be very similar in form, especially in early ontogeny (Paris *et al*. 2008; Laudet 2011; Napoli 2024). As growth occurs, variation at any life stage may remain minimal, as within nondomestic *Equus* species, and within *Dromaius novaehollandiae.* That appears to be the norm among nonavian dinosaurs including tyrannosaurins (Mallon 2017; Mallon & Hone 2024). Or variability may increase dramatically, as in species of *Homo* and *Otis.* Major adult differentiation is generally associated with sexual dimorphism. To put it another way, variation within a given species when present occurs with maturity, juveniles do not start out diverse and lose the variability as they grow up. In Reptilia/Sauropsida tooth counts nearly always remain stable with growth, or less often increase (Witmer & Ridgely 2010; Larson 2013b; Brown *et al*. 2015; Napoli 2024). Tooth loss with maturity is rare, being limited to captorhinid basal sauropsids (Haridy *et al*. 2018), fully marine ichthyosaurs that reduce teeth to a nonfunctional level (Dick & Maxwell 2015; Paul 2025a), and some centrosaurian basal avepod theropods that underwent a radical transformation from predaceous toothed juveniles to herbivorous beaked adults while they retained the alveoli (Wang *et al*, 2017; Paul 2024a, b; Zanno & Napoli 2025). Ontogenetic tooth loss does not occur in predaceous terrestrial reptiles, including *Gorgosaurus* and *Tarbosaurus* (Rozhdestvensky 1965; Currie 2003a; Longrich & Saitta 2024; Paul 2025a), and claims of such in *Alioramus* are unsubstantiated (Longrich & Saitta 2024; Paul 2025a). Tooth loss with ontogeny in synapsids is not particularly pertinent to this analysis, that remote group relative to diapsids undergoing a very different evolutionary path regarding dental adaptations (Benton 2014; Bertin *et al*. 2018; LeBlanc *et al*. 2018).

*Tarbosaurus* ontogeny is of critical importance in setting the parsimonious null hypothesis standard for assessing the ontogeny of its close relative. It was a similarly robust, gigantic fellow tyrannosaurin, the closest in form and function to the tyrant lizard yet known. The terrifying and tyrant lizards are so alike that they have sometimes been considered congeneric (Maleev 1955; Rozhdestvensky 1965; Paul 1988, 2008, 2010, 2016, 2024a; Carpenter 1992; Carr 1999, 2020, 2022, 2025). Its osteological ontogeny is well documented (Fig. 5A-H; Rozhdestvensky 1965; Carpenter 1992; Longrich & Saitta 2024; Paul 2020, 2016, 2024a,b, 2025a; Yun *et al*. 2025). Regarding the latter, Yun *et al*.’s (2025) extensive study on the cranial ontogeny of *Tarbosaurus* casually suggests that the growth patterns they observed generally parallel those of *Gorgosaurus* and *Tyrannosaurus*. This was not verified by applying the same methodologies to the American tyrannosaurids. Nor was the dramatic, sudden mid-growth metamorphosis proposed by Carr (2020) for *T. rex* observed in *T. bataar*. Also not considered by Yun *et al*. (2025) was the data and analysis presented by proponents of the MTTH, including the major new works by Longrich and Saitta (2024) and Paul (2025a). As a result, issues concerning tooth counts and form, lateral dentary grooves and other critical items were not addressed. It is concluded that the similarities in growth between the two tyrannosaurins Yun *et al*. (2025) commented on were only of the broadest nature, often related to increasing robustness with size, not verification that the atypical ETRH version of *Tyrannosaurus* growth closely matches that actually present in *Tarbosaurus*. To the contrary, viewed parsimoniously that two taxa that are so similar in their genetics and form and function experienced dramatically different ontogenies is sufficiently low that it must be a-priori assumed that the growth of the sister taxa was very similar. That means that showing otherwise has the major burden of evidence upon it. During the entire observed ontogeny from under 100 kg to over 4 tonnes of *Tarbosaurus* there is nothing out of the amniote/diapsid routine (Rozhdestvensky 1965; Carpenter 1992; Paul 2024a,b; Raun *et al*. 2025a; Yun *et al*. 2025). It shows a persistently consistent morphology that starts with basic features retained in the adults. These include a stable tooth count, absence of a long prominent dentary groove and diminutive forelimbs and manus that are about two thirds femur length at any growth stage, and a smooth allometric growth arc with no sudden metamorphic proportional and morphological alterations – *Tarbosaurus* always looked like *Tarbosaurus* as it matured. The degree of ontogenetic niche partitioning that occurred in *Taarbosaurus* was correspondingly modest, consisting largely of an allometric increase in robustness and moderate decrease in relative limb length, striking alterations to body form related to dramatic changes in predation are not observed. And immature *Tarbosaurus* are readily distinguishable from non-*Tarbosaurus* lesser sized taxa that lived in the same habitat (Fig. 5E-H versus 5I). That the maturation of the Nemegt tyrannosaurin was entirely normal bolsters it as a model to apply to its very close relation *Tyrannosaurus*. It follows that to convincingly establish that growth in the so similar *Tyrannosaurus* was dramatically outside the ordinary tarbosaur pattern is inherently nonparsimonious, and demands extraordinary positive evidence that such was the case. This conventional standard should have always been the norm in this field of research, and that it has not among so many has been a failing of dinosaurology. For the first time the conservative, parsimonious, null hypothesis analytical technique that needs to applied to the ontogenetic-taxonomic status of TT-zone lithe tyrannosaurs is comprehensively listed in Table 1.

**TABLE 1.** Growth restoration biological parameters.

It follows that if substantial osteological and dental differences are present between lesser sized individual TT-zone tyrannosaurs at given size stages, then they represent multiple taxa. In that case, those specimens that most closely match the features of adult *Tyrannosaurus* are the juveniles of that genus and its species. If growth slowed at smaller sizes according to bone ring spacing in preteen and teen aged individuals, then the possibility of being a juvenile of the fast-growing tyrant lizard giant is low at best. If anatomical variation within the fossils that do not qualify as immature *Tyrannosaurus* is itself extensive, then they very probably represent multiple taxa at at least the species level, and possibly higher depending on the degree of variation -- if some of those specimens possess nontyrannosaurid features such as long hands then they lie outside that family. The prior methods are entirely conservative and normal in nature. Doing anything outside those conventional procedures constitute at best radical propositions and hypotheses that require abundant positive evidence to support them. Building on prior work most recently by Longrich and Saitta (2024), Paul (2025a) and then Zanno & Napoli (2025), this study goes further to describe and apply these reasonable principles than any to date. These routine methods need to be adhered to as much as possible in all future work on the subject.

### Conventional, parsimonious criteria for examining and restoring growth in fossil amniotes, including TT-zone eutyrannosaurs **--**

Close correspondence in growth patterns with those of close relations, which in this case is albertosaurs and especially tarbosaurs, which include the below standard amniote/diapsid features.

Juveniles of a single species will be highly similar in core characteristics and are visual predecessors of their adult for -- this requires consideration of all TT-zone tyrannosaur fossils below two tonnes estimated live mass in order to assay all the variation they contain.

No sudden fish style metamorphosis including limb atrophy.

Reptilian tooth counts are always in a narrow range, and do not decrease with increasing size down to the adult counts.

Growth as recorded by bone rings is very likely to follow a fairly smooth arc peaking in mass accumulation in the teens.

In the case of exceptionally powerfully built *Tyrannosaurus,* its juveniles should be at least about as robustly built in dentition, skull and skeleton than are the young of the nearly as hefty *Tarbosaurus*, if not more so.

### Normal *Tyrannosaurus* growth recorded by actual juveniles of the genus --

There is a close correspondence with the normal amniote-diapsid-tyrannosaurid growth pattern of its fellow tyrannosaurin *Tarbosaurus*.

As far as is known the juveniles are both highly similar to one another, and are fairly robust, clear cut antecedents of their adult form.

There is no sudden fish style metamorphosis, including atrophy of the manus.

Dental counts are always in a narrow range, and do not experience a highly atypical decrease with increasing size down to the adult counts.

There is not a shift from a prominent lateral dentary groove to little or none. A diversion from a standard growth S-curve is not involved.

There is nothing radical or abnormal from ontogenetic norms.

### Aberrant improbable or impossible growth associated with and required by the ETRH **--**

A highly abnormal and divergent from the normal ontogeny of amniotes, diapsids, and tyrannosaurids that is drastically different from that of fellow tyrannosaurin *Tarbosaurus*, that is otherwise so similar it potentially qualifies as congeneric with *Tyrannosaurus* according to Carr.

Lack of a compelling explanation for the above when it does not occur elsewhere, other tyrannosaurids and tyrannosaurins included.

A unique or close to it divergence in dental and cranial attributes among juveniles of a given growth stage, covered over by neglecting the fossils that qualify as juvenile *Tyrannosaurus*.

A sudden metamorphosis of the type seen among some fish, but not amniotes.

A very rare contraction and decrease of reptilian tooth counts not seen in beakless predaceous theropods, tyrannosaurids and tyrannosaurins included.

Growth irregular, involving peculiar slow-downs among pre/teens.

### Notes on *Tyrannosaurus* species

Although the absolute measurement values by Carr *et al*. (2022) and Carr (2025) differ from those of Larson (2008), Paul *et al*. (2022) and Paul (2025a), the similar relative sizes of the left dentary 2^nd^ and 3^rd^ tooth positions (as measured by alveoli and/or tooth base diameter depending on what is available) of the *T. rex* CM 973 holotype (Fig. 2A) is affirmed, as is the smaller dimensions of the 1^st^ position. The result confirms the derived condition of only one small tooth at the anterior end of a CMNH 9380 dentary that had been rare in the lower TT-zone, and then become dominant in the upper. Carr *et al*. (2022) and Carr (2025) show that the situation regarding the right dentary differs, with the 2^nd^ position shorter than the 3^rd^. However, the overall cross-sectional area of the 2^nd^ position appears to be substantially greater than that of the 1^st^ in Figure 25B in Carr (2025). That suggests that the 2^nd^ tooth, which is not preserved, could have been markedly larger than is the preserved, slender incisiform at the tip of the element. That possibility favours at least a partial presence of the derived *Tyrannosaurus* condition in the holotype of the type species. The tooth position length ratio used by Larson (2008), Paul *et al*. (2022) and Paul (2025a) may have limits. A comparison of the cross sectional areas of dentary positions 1-3 in the *Tyrannosaurus* fossil sample could provide superior comparative results, but would be very difficult to produce due to practical logistical issues. In any case, citing one or a few exceptions to a larger pattern of character distribution as refuting its entire diagnostic utility in taxonomic work is not automatically definitive. Because intragenera species are so similar to one another, the diagnostic characters of sibling taxa often show overlap (Paul *et al*. 2022; Paul 2025a), as per the species of *Triceratops* (Scannella *et al*. 2014; contra the contention otherwise by Carr *et al*. 2022). Because bioevolution is irregular, not precise, there being divergence in the proportions of the right and left tooth positions in CM 9380 is not sufficient to negate the strong general trend of reduction of two small anterior dentary teeth down to one over time in *Tyrannosaurus*. All the more so because the change is in broad-spectrum tune with other, time related changes in the osteology of the genus that indicate it was speciating over time. The opinion that the pro-ETRH Carr (2020) constituted a prebuttal of MTTH Paul *et al*. (2022 has been damaged by the widely acknowledged rebuttal of the ETRH it advocated. Reinforcing intragenera sibling species is the readily visible greater variation in cranial anatomy, especially the very different postorbital bosses and interfenestral pillar, of *Tyrannosaurus* (Figs. 7F-J, 9B,C,E) compared to other avepods including specimens of *Tarbosaurus* (Fig. 5A-H; Paul 2025a), and species of *Daspletosaurus* (Fig. 7C-E) and *Allosaurus* (Fig. 7A,B). It is worth noting that the proportional means being used to distinguish intra *Tarbosaurus* sibling species (Raun *et al*. 2025b) are broadly similar in nature to those utilized for the same purpose for the tyrant lizard, and may be less extensive and stratigraphically correlated. If in the future earlier *T. imperator* is accepted as distinct from *T. rex* while contemporary *T. regina* is not (as implied by Longrich & Saitta 2024), that continues to treat *T. rex* as a special species that suddenly at the very end of the Mesozoic evolved unprecedented proportional and display diversity than seen prior in other tyrannosaur and dinosaur species (Paul *et al*. 2022; Paul 2025a). The presence of a robust and a gracile tyrannosaurid taxa in the same level of a formation being comparatively normal leaves it the parsimonious probability. Note that two giant tyrannosaurid, probably tyrannosaurins, taxa appear to be present in the same quarry from the Hongtuya Formation (Hone *et al*. 2011).

Voris *et al*. (2025) propose that the flat ventral margin of the “*T.” mcraeensis* holotype dentary is also observed in *Tyrannosaurus* specimen MOR 008, contradicting a taxonomic separation at least at the species level. But the dentary of the latter appears to be damaged, in which case it does not negate a well-founded taxonomic distinction presented by Dalman *et al*. (2024 that may be at the genus level (Paul 2025a). If Voris *et al*. (2025) are correct that “*T.” mcraeensis* is more contemporary with TT-zone *Tyrannosaurus* than thought by Dalman *et al*. (2024, then the giant tyrannosaur/s of the southwest appear to have retained into the later Maastrichtian the geographic differentiation common to western North America in the late Cretaceous.

### Specimens and work thereon

Specimens of modest dimensions under consideration are USNM 6183 (Hatcher), UCRC V1, LACM 23845 (*Dinotyrannus megagracilis* holotype), BHI 6439 (Nicklas), KU 156375 (Laurel), UCMP 84133, MOR 1189. Baby Bob, DMNS Teen Rex which can be at least provisionally assigned to *Tyrannosaurus*, and BMRP 2002.4.1 (Jane, *Gilmoretyrannus lethaeus* holotype) & 2006.4.4, (Petey), CMNH 7541 (*Nanotyrannus lancensis* holotype), DDM 344.1 (Tom), FNMH PR2411, HRS 08 (Zuri), 15001, Jodi, KU 155809 (Maddy), LACM 28471 (*Stygivenator molnari* holotype), MOR 6625 (Chomper), NCSM 40000 (Bloody Mary, ex BHI 6437, NCMNSBM in Paul, 2025a), RSM P2347.1, TMP 80.16.425 that lie outside the genus (Figs 1, 2B-I, 3C-E, 6C-E, 8B, 9G,H; Gilmore 1946; Molnar 1978; 1980; Bakker *et al*. 1988; Paul 1988, 1998 Fig. 3C, 2025; Olshevsky *et al*. 1995; Larson 2008, 2013a, b; Schmerge & Rothschild 2016a,b; Burnham *et al*. 2018; Paul *et al*. 2022; Longrich & Saitta 2024; Zanno & Napoli 2025; Woodward *et al*. 2026). That many of these specimens have not been described is not critical to taxonomic work, such being common in paleontological systematic work (as per the treatment of BMRP 2002.4.1 by Zanno & Napoli 2025). Tooth socket counts not in the literature are based on direct examination and/or high quality photographs. Preservation of these specimens ranges from essentially complete to very fragmentary, with all of the *Tyrannosaurus* being towards the latter end of the preservation spectrum. None of these specimens indicates individuals below a few hundred kilograms, unlike *Tarbosaurus* juveniles of less than 100 kg (Fig. 5G,H; Paul 2024a, b). The putative non*Tyrannosaurus* remains are broadly similar in size (Figs. 1, 2E-I, 3C,D), precluding major ontogenetically driven shape reformation as being responsible for the extensive variation that is present. Also precluded is strong sexual dimorphism in such immature individuals. With the TT-zone having been deposited over 1+ MA (Paul *et al*. 2022 and refs. therein; Zanno & Napoli 2025) this is not an issue regarding subfamilies and genera, but can be for species. Appalachian baso-eutyrannosaurs date from the late Campanian (*Appalachiosaurus*, *Carr et al.* 2005) to the late Maastrichtian (*Dryptosaurus,* Carpenter *et al*. 1997; Gallagher 2023).

Cranium only *N. lancensis* (Gilmore 1946) and rostrum only *S. molnari* (Paul 1988) were named on incomplete materials when the TT-zone tyrannosaur fossil record was sparse. It has since become extensive, allowing new holotypes to be based on specimens that are largely complete regarding both the cranium and postcranium. It is strongly recommended that high skull-skeletal fossil completeness be a strict requirement for any future new tyrannosaur taxa from the TT-zone complex of formations. This is in accord with Paul *et al*. (2022) and Paul (2025a) in which the new *Tyrannosaurus* species were likewise founded only on specimens possessing sufficient diagnostic skull-skeletal features including orbital display in association with skull and skeletal proportions including the diagnostically critical complete femur, and basic stratigraphic levels were known – classic AMNH 5027 for instance is an unsuitable holotype on some of these grounds.

Use of profile (rather than shaded or photographic) images has the advantage of maximizing visual comparisons of critical shape and proportional differences. They also have the advantage of being able to present more direct plan views of the anatomy of skulls who actual depth/length proportions may be obscured in photographs of complex topography crania. Those in Figures 1,2,4,5,7,9 are derived in part from Paul (2025a) as noted in captions, with modification and additions. Illustrations and restorations (Figs. 1-9) were produced following standard procedures for generating technical paleoillustrations that are as meticulous as possible, and are based on personal examination, scans, photographs and illustrations, including those in Gilmore (1946), Witmer and Ridgely (2010), Larson (2013a), Longrich & Saitta (2024) and Zanno & Napoli (2025). Bones have been carefully traced from source images -- any who disagree with their accuracy need to demonstrate each of the errors with measurements and/or comparative images – the first alone can be confusing and misleading. Elements including teeth from one aide are used to fill in those missing from the other. What bones are or are not known is not always clear, many postcranial elements of the complete NCSM 40000 are concealed for the long term. None of the baso-eutyrannosaur skulls restored (Fig. 1) are complete, fully articulated and undistorted, so all require considerable restoration. Occasional subtle differences between skull elements in different figures are due to elements from different specimens and/or sides being used. The left sides that are restored are often compromises to varying degrees that naturally result from combing right with left elements to obtain an average. Due to these factors all intraskull restored dimensional ratios are approximations to varying degrees – total skull length (Fig. 1 caption) is premaxilla tip to tip of paroccipital wing; temporal box length is latter to anterior edge of lacrimal preorbital bar; rostrum length is latter to premaxilla tip; rostrum height is measured at the anterior end of the antorbital fossa; general skull height is measured at the postorbital bar vertical sans any ventral jugal cornual rugosity; dentary length is anterior tip to posterior edge at intermandibular fenestra; dentary depth is at mid length; posterior mandible height is maximum depth at about mid length. Profile-skeletals that are based on fragmentary specimens involve extensive extrapolation and cross-scaling from related taxon. For example, the shape and size of the cranium of the Appalachia *Dryptosaurus* dimensionally stems from the depth of its dentary applied to that of its baso-eutyrannosaur relation *Appalachiosaurus* (Fig. 3A,B). The body form of the modest sized USNM 555000 is downsized to accommodate the remains of the large juvenile LACM 23845 (Fig. 9D,G).

BMRP 2002.4.1 (Figs. 1B, 2G, 3D) has yet to be described -- Zanno & Napoli (2025) proceeded to diagnose it as the holotype of a new species before that work has been presented (by Carr). Parts of the specimen have been extensively imaged in the technical literature (Larson 2013a; Longrich & Saitta 2024; Zanno & Napoli 2025) and elsewhere, many measurements have not been published. The skull elements appear little distorted, but are disarticulated, the absence of the frontals and braincase precludes an accurate dorsal view restoration. The lateral view presents fairly typical eutyrannosaur proportions. General height/total length is a moderately low ∼0.24. Rostrum height/total length also is a moderately low ∼0.21. Rostrum/temporal box length is a long ∼1.65. Dentary height/length is ∼0.16, but that value is on the low side because the mid dentary is vertically pinched relative to the ends. The fairly complete HRS 08 is missing too much of the temporal region and posterior mandible to be reliably restored for these purposes, what is known appears to have a similar general shape to BMRP 2002.4.1 (including the profiles of the dentary and maxilla, although the antorbital fossa of the latter is distinctive (Fig. 2G,I).

NCSM 40000 (Fig. 1C) has been technically extensively documented, measured and diagnosed in a major paper as within *N. lancensis* by Zanno & Napoli (2025), who then used it to partly characterize the taxon. Paul (2025a had provisionally diagnosed it as a probable new taxon, possibly a species within *Stygivenator*). NCSM 40000 is further available as images in various locations. The accessioned and published specimen therefore meets all modern standards for further taxonomic analysis, rendering it even more available for holotype designation than was BMRP 2002.4.1. Damaged by amateur excavation before professionals arrived, the skeleton is undergoing long term preparation, and extensive soft tissues obscure much of the presacral series and thorax, so the profile-skeletal (Fig. 3C) is the most that can be generated for the foreseeable future. Images utilized are Figures 2, 5 and Extended Data Figures 1, 2 from Zanno & Napoli (2025) supplemented by online photographs, Measurements are from their Table SI.1.5. The cross scaling of the detached rostral and temporal sections of the cranium in Zanno and Napoli’s (2025) Extended Data Figure 3 is in good general accord with those present in online photographs showing the skull in its sediment block. In Extended Data Figure 3c,e in Zanno and Napoli’s (2025 the two images are not exactly the same scale, c being reproduced ∼4 larger than e). Breakage of the lacrimals in the region of the dorsal hornlets tilted the roof elements anterior to them, giving the illusion of a flexed skull at mid-length. The false curve has been removed, leaving the skull straight in lateral profile. The correction also shows the hornlets have the normal single peak. Because both sides of the NCSM 40000 cranial temporal box are somewhat distorted and damaged (EDFig. 3 in Zanno & Napoli 2025), the compromises resulting from using right and left elements leaves the exact shape and dimensions of the orbit uncertain, but it was rather small. That said, the generally good similarity of the shape and proportions of elements from the different sides do not appear to record serious dorso-ventral cranial crushing. The temporal region is skewed somewhat to the right dorsally (EDFIg. 3b,d in Zanno & Napoli 2025), but not to the degree of seriously impacting the similar depths and configurations of the lateral surfaces. The length/height ratio of the fossil’s temporal box, is replicated in the restoration. The maxillae are very similar in their low height and acute triangular shape, any difference not being sufficient to significantly alter the profile of the shallow, sharp rostrum, which is a close match on both sides as seen in Extended Data Figure 3c,e in Zanno & Napoli (2025), especially when the differing scales of the two images are adjusted as noted above. The rim of the right anterior antorbital fenestra is a little shallower and more sharp angled than the left (Fig. 2E). The former condition is seen in the articulated skull image in Zanno & Napoli (EDFig. 3a). The discrepancy may be the result of right/left asymmetry and/or minor dorso-ventral crushing, the left version is used in the final restorations. The elongated alioramin-like (Paul 2010, 2016, 2024a,b; Brusatte *et al*. 2012) cranial configuration is similar to that present in Extended Data Fig. 3a in Zanno & Napoli (2025) as well as online photographs of the skull in the sediment block. The fore versus aft sections ratios are inherently approximate because of the detachment of the fore and aft sections precludes precision. The slightly ventrally disarticulated premaxillae are moved dorsally a little to bring their ventral margins correctly in the same line as those of the maxillae. Although the dentaries are tightly articulated with the rostrum, a short portion of the right dentary’s dorsal edge exposed with a tooth in place indicates their depth. The maximum total length of the NCSM 40000 restoration is a little shorter than in EDFigure 3a and Table SI.1.5 in Zanno & Napoli (2025) entirely because of the proper close application of the paroccipital wing to the posterior surface of the squamosal. General height/total length is a shallow ∼0.22. Total skull length/rostrum height is a very shallow 0.19 to 0.2, the detachment of the front and aft halves precludes a tighter value. Rostrum/temporal box length is a long high 1.65 to 1.85, again the detachment of the front and aft halves precludes a tighter value. Dentary height/length appears to be consistently shallow with a minimum of ∼0.16. Temporal box/posterior mandible height is low at ∼1.35. Posterior mandible height/total skull length is a shallow ∼0.17. How reliably the cranium can be restored in dorsal view is not clear.

The largely albeit not entirely complete and significantly damaged CMNH 7541 (Fig. 1A) can be restored in lateral view, although doing so faces numerous issues due to the missing bone elements and perplexing distortions. Judging from Gilmore’s (1946), the specimen is a single block including the tightly articulated mandibles, in which case the basic original length is preserved. The mid maxillae are fragmented and much of the skull roof is absent (Fig. 4 in Witmer and Ridgely 2010), but this has little effect on the overall length of the cranium. Witmer and Ridgley (2010) praised the original restoration work. The general shapes of the maxillae and antorbital fossae are preserved, particularly their antero-posteriorly short, vertically tall and correspondingly blunt proportions. Comparing different photographs and scans (as per those in Witmer & Ridgley 2010) differences in proportions of the overall cranium and elements are unusually substantial. This is because the temporal box is strongly splayed out ventrally relative to the roof (Figs. 2B.C3A,B, 4A,D-F in Witmer & Ridgely 2010). As a result the aft half of the skull appears shallower than it actually is when the skull roof is posed perpendicular to the viewer (as in Pl. 1 in Gilmore 1946; Fig. 8.8 in Larson 2008; Figs 1, 3B,C in Witmer & Ridgely 2010; Fig. 3A,B in Longrich & Saitta 2024). Illustrations of the skull have been prone to be correspondingly too shallow, especially aft (Fig. 3 in Bakker *et al*. 1988; Fig. 6 in Carr 1999; Fig. 3 in Longrich & Saitta 2025). The excessive beam of the temporal box is a major factor in the additional variations between photographs, because small differences in camera angles, distances and lenses are exaggerated in the resulting images. Restoring the lateral view most accurately was achieved by rotating an online 3-D scan (sketchfab.com/3d-models/nanotyrannus-lancensis-young-t-rex-7b0967fa27674d959647868686b6717b) until the left side was maximally deep and correspondingly as flat on as possible to present a direct lateral plan view. This reveals that the skull was distinctly deep both fore and aft, more so than the norm for eutyrannosaurs of this size. The frontoparietal, sagittal and occipital crests are exceptionally vertically prominent The quadrates and quadratojugal have been pushed forward and somewhat medially, shortening the lateral temporal fenestra. This distortion was corrected to allow room for the anterior rami of the quadratojugal and squamosal, which lengthens the skull ∼4%. Even with the correction, the lateral temporal fenestra appears antero-posteriorly shorter than the eutyrannosaurs norm, in association with the usually tall postorbital and post temporal fenestra bars. The posterior view (Fig. 1A) is de-distorted to show the post temporal bar properly subvertical. The mandible is primarily after Figure 15C in Witmer and Ridgely (2010); because the lower jaw is severely jammed up into the skull (far more than possible in life, p. 34 in Paul 2024b, and too the degree that the teeth have been pushed postero-ventrally) its massive depth has not been previously realized. That is in part because most side view images of the preserved right dentary on not flat on to the element, as it is in the Witmer and Ridgely and online scans. The position of the detached anteriormost dentary relative to the rest of the lower jaws in not certain. General height/total length is a deep ∼0.28. Rostrum height/total length also is a deep ∼0.25. Rostrum/temporal box length is a short ∼1.45. Dentary height/dentary length is a deep ∼0.18. Temporal box/posterior mandible height is a high ∼1.4. Posterior mandible height/total skull length is a deep ∼0.22. Because CMNH 7541 has been asymmetrically distorted in a complex manner, and extensive dorsal cranial elements are missing (Fig. 8B), it proved impractical to reliably restore its configuration in dorsal view (Paul 2025a; Zanno & Napoli 2025). The very narrow CMNH 7541 rostrum appears to be laterally compressed an uncertain amount (but Longrich & Saitta 2024 disagree). Carr’s (1999) restoration of the rostrum as broader than preserved (Fig. 8C) is as plausible if not probable, as it is nonconfirmable. Carr’s viable version is more similar in proportions to that of *Daspletosaurus* than *Tyrannosaurus* (Fig. 8A,D), so the reasonable restored dorsal dimensions do not particularly favor the specimen being within the tyrant lizard ((Paul 2025a and then Zanno & Napoli 2025). Because any effort to restore CMNH 7541 in dorsal aspect produces an arbitrary rather than scientifically scoreable and diagnosable result, the item cannot be used for phylogenetic or gradistic assessments as explained by Paul (2025a) and then Zanno & Napoli (2025). The absence of a prefrontonasal processes of the frontals observed by Voris *et al*. (2025) may be due to the large break of this skull at this position (Fig. 6B in Carr 1999; Fig. 3B in Witmer & Ridgley 2010) leading to the loss of the small and delicate processes (see Fig. 5C in Carr 1999), and cannot be reliably scored as a character either. Longrich & Saitta (2024) note that Carr (Fig. 6 in 1999 in which the lateral restorations do not entirely match each other, Fig. 12 in 2020) appears to have restored the skull in a manner that makes it look more like *Tyrannosaurus* than it actually is, including premaxillary teeth that are too vertical, a snout tip that is too broad U-shaped in dorso-ventral view, and a maxillary fenestra that is placed too far anteriorly, especially in one version.

**FIGURE 8.**
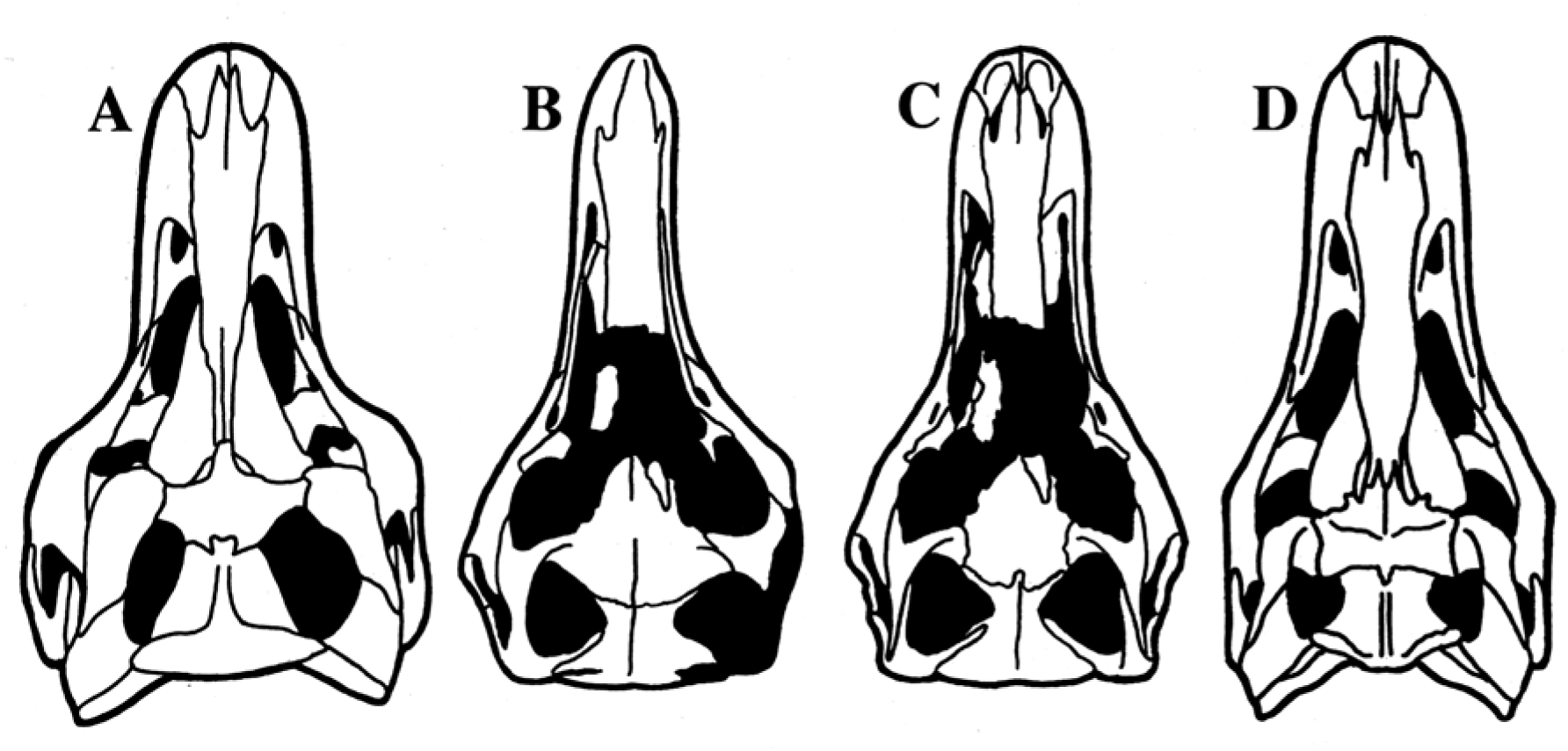
Known-bone eutyrannosaur crania in dorsal view reproduced to same midline length to facilitate intergenera visual comparisons. **A** *Tyrannosaurus* AMNH 5027; **B** *Nanotyrannus* holotype CMNH 7541 as preserved from 3-D scan, extent of preserved elements aided by Figures 3A, 4D in Witmer and Ridgely (2010); **C** CMNH 7541 as restored by Carr (1999); **D** *Daspletosaurus* holotype CMN 8506.

Conspicuous differences in the vertical versus the longitudinal dimensions of the elements of the skulls that are presented in Figure 1 support the variations in their restored overall length/depth ratios. The CMNH 7541 quadratojugal is tall, the vertical height approaching the horizontal distance between the anterior tip of the maxilla and its antorbital fossa. The shorter BMRP 2002.4.1 quadratojugal is just ∼80% the length of the anterior maxilla, in line with a shallower cranium. The even shorter quadratojugals of NCSM 40000 are about three quarters the anterior maxilla length, in accord with its exceptionally low cranial profile vis-à-vis the other specimens. The lower height to length of the maxilla also contributes to its very shallow skull. To stretch the *Nanotyrannus* cranium to match the lower profiles of the others would require large arbitrary space gaps between the fragments of the maxillae that are not preserved in the fossil.

The *Appalachiosaurus montgomeriensis* holotype RMM 6670 skull restoration (Fig. 3A) is little different from that of Carr *et al*. (2025). The incomplete elements preclude ratio calculations, but the general boxy proportions appear more like that of CMNH 7541 than the elongated NCSM 40000.

Produced via standard consistent comparative techniques (Paul 1988, 2024a,b, Larramendi *et al*. 2021), volumetric mass estimates for specimens as preserved, or projected for adults, are derived in part from mass and growth stage data in Paul (2024b, 2025a), Zanno & Napoli (2025) and Griffin *et al*. (2025), with modifications herein. As per the diapsid-dinosaur norm (Paul 1988, 2024a,b), articulated tyrannosaur skeletons preserve the trunk ribs swept postero-ventrally (as per Zanno & Napoli 2025), which significantly reduces the volume of the trunk compared to restorations showing the chest ribs vertical or swept ventro-anteriorly (contra the mounted skeleton scans in Hutchinson *et al*. 2011). The latter incorrectly inflates the volume/mass of the restoration. So do restorations showing the gastralia basket bowed convex ventrally; such represents the subject distended by gorging on a carcass, or post-mortem bloating as is commonly preserved in the fossils. Profiles are restored in the healthy lean-hungry condition suitable for flesh eaters on the hunt, with empty hollow bellies, and the minimal fat deposits that maximize speed performance and agility in pursuit predators. Use of bone circumferences for mass estimation has higher +/- errors; mass/circumference ratios can differ considerably within species due to sexual, individual and other variations, even more between species, yet more between genera. Nor can circumference based estimates take into account dimensional differentiations present in actual fossil skeletons. Using circumferences led Zanno & Napoli (2025) to estimate that BMRP 2002.4.1 was ∼18% heavier than NCSM 40000, this result is not sufficiently dependable for comparative size work because of the inherent +/- error. The pelvis and tail of the latter appear larger than that of the former (Fig. 3C,D) – the two specimens have similar sized limbs – and the volumetric method favours 40000 being at least as weighty as 2002.4.1. Because the latter was not as far along in its growth (Paul 2025a; Zanno & Napoli 2025) the adult mass of its species was probably substantially higher than that of NCSM 40000, as per their systematic diagnoses.

Systematic diagnoses are based on Paul (2025a which was based in part on Longrich & Saitta 2024), Zanno & Napoli (2025) and work herein.

### Systematic analysis and diagnoses

Systematic diagnoses for assessing North American Campanian/Maastrichtian baso-eutyrannosaur and TT-zone tyrannosaurid taxonomics are based largely on Paul (2025a) which are in part based on previous efforts referenced therein, with additions and revisions following the characters employed in the phylogenetic matrix (see above and Appendix 1). The diagnoses are differential cumulative character compilations for each taxon that when combined with some distinctive characters/apomorphies (indicated with an*) are intended to detect and describe gradistic separation of genera and species. Some characters known from only some members of a group are listed as general attributes of that group, with exceptions noted in the genus or species diagnoses when such is known to be the case. Because the number of *Tyrannosaurus* intragenera species is not a focus of this study, that taxon is diagnosed only at the genus level, with an emphasis on characters known to be present in juvenile remains. Diagnostic characters are sometimes overlapping non-bimodal between taxa at a given level, and not entirely consistent within taxon as per Maisch (2008), Maxwell (2012), Scannella *et al*. (2014), MacDonald & Currie (2018), Harvati & Ackermann (2022), Paul *et al*. (2022), Carr *et al*. (2022), Longrich & Saitta (2024). A few specimens have been reassigned or their status modified relative to Paul (2025a). Because the cumulative evidence presented in this analysis finds that some small TT-zone tyrannosaur fossils are not juvenile *Tyrannosaurus*, and as near adults they possessed adult attributes, their anatomical characteristics can be used to taxonomically define and diagnose them differentially from adult tyrannosaurins (Bakker *et al*. 1988; Larson 2008, 2013a, b; Schmerge & Rothschild 2016a,b; Longrich & Saitta 2024; Paul 2025a; Zanno & Napoli 2025).

### Systematic palaeontology North America baso-eutyrannosaurs general

#### Character attributes

1-2 tonnes, at least fairly gracile; anterior margin of premaxilla sloped dorsoposteriorily, ventral margin anteriorly upturned, form narrow U or V in dorso/ventral view, subnarial process faces anterolaterally, subnarial foramen absent, nasals narrow except broad where contact frontals, transversely fairly flat, contact with maxilla fairly smooth, maxilla long and low, ventral margin gently convex, antorbital fossa shallow, accessory antorbital fenestra small and does not contract rim of antorbital fossa, interfenestral pillar always broad, antorbital fossa rim broad along lower edge of antorbital fenestra, promaxillary recess small, posterodorsal process long and robust, posterodorsal process posteriorly elongated, deep medial recess above antorbital fenestra absent, medial sinus above antorbital fenestra small and shallow, medial antorbital fossa weakly developed, palatal shelves not elevated, lacrimal more T shaped with long posterodorsal projection, subtriangular hornlet present, antorbital fossa contribution deep, frontals participate in orbit rim, long and narrow in dorsal view although broad between lacrimals, contribute to short portion of sagittal crest, supratemporal fossa short, parietal nuchal crest broad, postorbital main body shallow, boss is subtle subcircular, knob-like discs limited to the frontal process that do not project above the dorsal rim of the skull, jugal process narrow and lacks orbital flange, jugal contact fairly straight, jugal anterior wing shallow, orbital margin long and gently curved, antorbital fossa contribution shallow, pneumatic recess shallow, quadratojugal process short, quadratojugal jugal process shallow: squamosal not large, ventral fossa lacks pneumatic recess, quadratojugal lateral foramen present, tip of quadratojugal process not squared off, abducens nerve nearly intersects pituitary fossa so sinus cavernosus, ascending diverticulum of anterior rostral tympanic recess present, condylotuberal crests strong, subsellar sinus extends into cultriform process, vomer anteriorly narrow and deep, palatine anterior processes long and slender, pneumatic fossa small, body weakly inflated, ectopterygoid pneumatic opening is a thin slot, dentary shallow, fairly straight in dorsal view, symphysis weak, long and prominent lateral groove usually present, interdental plates weakly developed, surangular contact slopes strongly anterodorsally, surangular shallow, lateral shelf short and not prominent, premaxillary teeth procumbent, lack serrations, atypically blunted, 15–16 maxillary and 16–17 dentary teeth that are bladed, anteriormost maxillary tooth incisiform; proatlantal arches strap-like; scapula blade neck not especially narrow, glenoid fossa extends onto lateral surface of scapula, humerus head not massive, forelimb often if not always elongated distally, especially manus, so forelimb is about same length as femur or greater, and manus absolutely larger than those of adult tyrannosaurins; ilium shallow, dorsal rim fairly straight, anteroventral prong hook shaped, pubic boot gracile, femur fourth trochanter weakly developed, tibia cnemial crest gracile, rounded.

**Genus Appalachiosaurus Carr et al., 2005**

**Type species. Appalachiosaurus montgomeriensis Carr et al., 2005**

**Holotype.** RMM 6670.

**Diagnosis.** 700 kg; Rostrum deep, subrectangular because of maxilla and anterior fossa boundaries forming an angle over 30 degrees, promaxillary fenestra not visible in lateral view; dentary tip not upcurved; 15 maxillary teeth, 7 teeth anterior to anterior end of antorbital fossa; femoral shaft strongly bowed; femur and tibia similar in length, metatarsus not highly elongated.

**Locality and horizon.** Late Campanian, lower Demopolis, Alabama.

**Genus *Dryptosaurus* Marsh, 1887**

**Type species. *Dryptosaurus aquilunguis* Marsh, 1887**

**Holotype.** ANSP 9995.

**Diagnosis.** 700 kg; manal digit 1 claw very large, humerus/femur ratio 0.375, manal phalanx 1-2/femur ratio 0.22 and forelimb probably longer than femur; femoral shaft nearly straight*, femur and tibia similar in length, metatarsus not highly elongated. **Locality and horizon.** Late/st Maastrichtian, upper New Egypt, New Jersey.

**Genus *Nanotyrannus* Bakker *et al*., 1988**

**Type species. *Nanotyrannus lancensis* Gilmore, 1946**

**Holotype.** CMNH 7541.

**Diagnosis.** ∼500 kg; Mature skull fairly deep, subrectangular because of deep, blunt rostrum with maxilla and anterior fossa boundaries forming an angle over 30 degrees, 8 teeth anterior to anterior end of antorbital fossa, promaxillary fenestra not visible in lateral view, orbit large, jugal large and tall*, quadratojugal tall, lateral temporal fenestra tall, jugal ventral cornual boss prominent, postorbital dorsal prong absent, frontoparietal peak prominent, quadratojugal-quadrate foramen large, paroccipital processes subhorizontal and distal ventral edge pendant, anterior palatine recess forms discrete pneumatic sinus extending anteriorly into maxillary process of palatine, dentary tip not upcurved and has prominent chin, dentary deep, posterior mandible deep, two small anterior dentary incisorforms, 15 maxillary and 16 dentary teeth, maxillary teeth neither large nor strongly recurved.

**Locality and horizon.** Late Maastrichtian, lower? Hell Creek, Montana.

Genus Gilmoretyrannus gen. nov.

**Etymology.** In honor of Charles W. Gilmore for recognizing the existence of multiple tyrannosaur taxa in the TT-zone

**Type species. *Gilmoretyrannus lethaeus* Zanno & Napoli, 2025**

*= Nanotyrannus lancensis* in Larson, 2013

*=Nanotyrannus lethaeus* (Zanno & Napoli, 2025) Bakker *et al*. 1988

**Holotype.** BMRP 2002.4.1

**Potential referred specimens.** BMRP 2006.4.4? HRS 08?, HRS 15001?

**Diagnosis.** ∼1 tonne; skull fairly shallow, subrectangular, 6 teeth anterior to anterior end of antorbital fossa, promaxillary fenestra not visible in lateral view, maxillary fenestra large, maxillary flange weak, orbit not large, jugal, quadratojugal and lateral temporal fenestra not tall, dorsal prong present on squamosal process of postorbital, anterior palatine recess forms open fossa, anterior buttress on jugal process of palatine present, dentary tip not upcurved and has prominent chin, dentary moderately deep, posterior mandible moderately deep, probably 1 small anterior dentary incisiform, 15 maxillary and 17 dentary teeth, maxillary teeth fairly large and not strongly recurved; axial neural spine narrow and sharp tipped in lateral view, distinctly crenulated along entire anterior rim, single pleurocoel on axial centrum, caudal vertebrae apneumatic; entepicondyle of humerus subvertical in distal view, humerus/femur ratio ∼0.38; strongly dorsally convex margin of posterior portion of pubic boot, pubic shaft bowed posteriorily, tibia markedly longer than femur, metatarsus very elongated.

**Locality and horizon.** Hell Creek, Lance, upper? TT-zone; Montana, Wyoming.

**Tribe Larsonvenatorini tribus nov.**

**Type genus.** Larsonvenator

**Diagnosis.** Skull subtriangular because of sharp rostrum with maxilla and anterior fossa boundaries acute at an angle of ∼30 degrees*, dentary chinless and upcurved*.

**Genus *Stygivenator* Olshevsky, Ford & Yamamoto, 1995**

**Type species. Stygivenator molnari Paul, 1988**

**Holotype.** LACM 28471

**Diagnosis.** 5 teeth anterior to anterior end of antorbital fossa, promaxillary fenestra not visible in lateral view, maxillary teeth large and strongly recurved except for one anterior small incisoriform, lateral dentary groove absent; height of the largest anterior maxillary tooth equals or even surpasses the depth of the dentary at its location*

**Locality and horizon.** Late Maastrichtian, upper? Hell Creek; Montana.

Genus *Larsonvenator* gen. nov.

**Etymology.** In honor of Peter Larson who excavated the type specimen, and for his extensive research on the TT-zone tyrannosaur taxa great and small

Type species. *Larsonvenator elegans* sp. nov.

*=Nanotyrannus lancensis* (Gilmore, 1946) Bakker *et al*. 1988 in Zanno & Napoli 2025

*=Stygivenator molnari* (Paul, 1988) Olshevsky *et al*. 1995 in Longrich & Saitta 2024

*=Stygivenator sp.* (Paul, 2025a) Olshevsky *et al*. 1995

**Etymology.** *Elegans* meaning elegant in latin.

**Holotype.** NCSM 40000 (ex BHI 6437).

**Provisional referred specimen**. Jodi.

**Diagnosis.** 700 kg; skull very shallow, 8 teeth anterior to anterior end of antorbital fossa, promaxillary fenestra visible in lateral view, maxillary fenestra small, orbit small*, jugal not tall, quadratojugal short, lateral temporal fenestra short, postorbital dorsal prong absent, frontoparietal peak low, quadratojugal-quadrate foramen small, paroccipital processes slope lateroventrally and distal ventral edge straight, anterior palatine recess forms discrete pneumatic sinus extending anteriorly into maxillary process of palatine, dentary shallow, posterior mandible shallow, 16-17* maxillary teeth are neither large nor strongly recurved, 2 small anterior incisiforms, 17-18 dentary teeth; height of the largest anterior maxillary tooth do not surpasses the depth of the dentary at its location; axial neural spine broad in lateral view, anterior rim poorly crenulated, two or more pleurocoels set in fossa on axial centrum*, infraprezygapophyseal pneumatic recesses on anterior proximal caudal vertebrae, caudal count 35; bulbous supraradiocondylar ridge and pronounced craniomedially projecting entepicondyle on humerus, pubic shaft straight, boot has straight anterodorsal, posterodorsal and ventral margins, humerus/femur ratio 0.39, manal phalanx I-1/femur ratio 0.22 and forelimb longer than femur, very elongated manal phalange I-1 markedly longer than those of adult *Tyrannosaurus,* as are manal claws, remnant phalange on manal digit 3; femoral shaft strongly bowed; tibia markedly longer than femur, metatarsus very elongated.

**Locality and horizon.** Late Maastrichtian, lower Hell Creek; Montana.

**Subfamily Tyrannosaurinae Olshevsky, Ford & Yamamoto 1995**

**Diagnosis**. 4+ tonnes, robust; rostrum deep, anterior margin of premaxilla vertical, ventral margin flat, form broad U in dorso/ventral view, subnarial process faces anteriorly, subnarial foramen present, nasals broad except narrow where contact frontals, transversely arced, contact with maxilla irregular, maxilla deep, ventral margin strongly convex, antorbital fossa deep, premaxillary fenestra not visible in lateral view, accessory antorbital fenestra fairly large and contracts rim of antorbital fossa, interfenestral pillar breadth variable, antorbital fossa rim narrow along lower edge of antorbital fenestra, promaxillary recess large, posterodorsal process short and slender, posterodorsal process posteriorly abbreviated, deep medial recess above antorbital fenestra present, medial sinus above antorbital fenestra large and deep, medial antorbital fossa well developed, palatal shelves elevated, lacrimal more L shaped with short posterodorsal projection, hornlet very shallow or absent, antorbital fossa contribution shallow, frontals do not participate in orbit rim, short and broad in dorsal view although narrow between lacrimals, contribute to long portion of sagittal crest, supratemporal fossa long, frontoparietal peak prominent, parietal nuchal crest is a ridge, postorbital main body deep, bosses highly variable in size and shape, jugal process broad and has often prominent orbital flange, jugal contact convex ventrally, jugal anterior wing deep, orbital margin short and strongly in cut, antorbital fossa contribution deep, pneumatic recess deep, jugal ventral cornual boss prominent, quadratojugal process fairly long, quadratojugal jugal process fairly deep, quadratojugal lateral foramen absent, intra quadratojugal-squamosal fenestra absent, squamosal large, ventral fossa has pneumatic recess, tip of quadratojugal process squared off, paroccipital processes subhorizontal and distal ventral edge pendant, abducens nerve does not nearly intersects pituitary fossa so sinus not cavernosus, ascending diverticulum of the anterior tympanic recess absent, condylotuberal crests strong, subsellar sinus extends into cultriform process, vomer anteriorly broad, palatine anterior processes short and robust, pneumatic fossa large, body inflated, ectopterygoid pneumatic opening is large; dentary tip not upcurved and has prominent chin, dentary deep, laterally bowed in dorsal view, symphysis well developed, prominent lateral groove absent, interdental plates well developed, surangular contact subvertical, posterior mandible deep, lateral shelf long and prominent, premaxillary teeth vertical, serrated, crowns not blunted, 11–13 maxillary and 12–15 dentary teeth at all growth stages, anteriormost maxillary tooth nonincisiform; proatlantal arches triradiate, caudal count 40+, mid-posterior centra not elongated, transverse processes expanded posteriorily, proximal haemal arches vertically elongate, mid heamal arches deeply curved; scapula blade neck slender, glenoid fossa does not extend onto lateral surface of scapula, forelimb very reduced*, manus especially, to about two thirds femur length, humerus/femur ratio -0.27-.034, manal phalanx 1-2/femur ratio 0.053-0.96, mature manus absolutely smaller than those of at least some adult baso-eutyrannosaurs, first manal phalanx of digit 1 not elongated, no phalanges on manal digit 3; ilium deep, dorsal rim dorsally arced, anteroventral prong weakly developed, pubic boot large; pubic shaft fairly straight, femoral shaft strongly bowed, femur fourth trochanter well developed, tibia cnemial crest large, squared off, tibia not elongated relative to femur even in juveniles, metatarsus not strongly elongated.

**Genus *Tyrannosaurus* Osborn, 1905**

= *Dinotyrannus* Olshevsky, Ford & Yamamoto, 1995)

**Diagnosis.** 7.5 tonnes*; 4-5 teeth anterior to anterior end of antorbital fossa, presence of anterodorsal process on anterior ramus of lacrimal that projects into nasal, dorsal lacrimals nearly meet at midline*, sublunate in dorsal shape partly because lateral swelling on supraorbital process is absent*, quadratojugal foramen absent, quadratojugal-quadrate foramen small, vomer sometimes has anterior spear point, contact with premaxilla not as extensive, anterior prong shallow, deep ventral flange absent, dentary lacks significant lateral groove, 11–12 maxillary and 12–14 dentary teeth at all growth stages, two (*T. imperator*) or one (*T. rex, T. regina*) small anterior dentary incisiforms, sometimes very large teeth robust; axial neural spine broad in lateral view, caudal count 42-43; humerus/femur ratio ∼0.3; phalanx 1-2/femur ratio ∼0.6; pubic boot massive.

**Type species *Tyrannosaurus rex* Osborn, 1905**

**Holotype.** CM 9380.

**Referred specimens.** As per Paul (2025a)

**Diagnosis.** As per Paul (2025a).

**Locality and horizon.** Latest Maastrichtian, upper and possibly middle Hell Creek and Lance, Ferris, Denver, Frenchman, Willow Creek, lower Scollard; Montana, Colorado, Dakotas, Wyoming, Alberta, Saskatchewan.

*Tyrannosaurus imperator* Paul *et al*., 2022

**Holotype.** FMNH PR2081.

**Referred specimens.** As per Paul (2025a).

**Diagnosis.** As per Paul (2025a).

**Locality and horizon.** Late Maastrichtian, lower, lower middle and possibly middle Hell Creek and Lance, Laramie, Arapahoe; Montana, Dakotas, Wyoming, Colorado.

*Tyrannosaurus regina* Paul *et al*., 2022

**Holotype.** USNM 555000 (MOR 555).

**Referred specimens.** As per Paul (2025a).

**Diagnosis.** As per Paul (2025a).

**Locality and horizon.** Latest Maastrichtian, upper and possibly middle Hell Creek and Lance, Ferris, Denver, Frenchman, Willow Creek, lower Scollard, lower North Horn?; Montana, Colorado, Dakotas, Wyoming, Alberta, Saskatchewan.

## Results and Discussion

### Phylogenetics

The phylogenetic results (AppendFig. 1) are broadly similar to other recent efforts in general aspects, with the same taxa placing as either noneutyrannosaur baso-tyrannosauroids or as eutyrannosaurs, and among the latter as baso-eutyrannosaurs versus tyrannosaurids (Lu *et al*. 2014; Brusatte & Carr 2016; Carr *et al*. 2017; Dalman *et al*. 2024; Longrich & Saitta 2024; Voris *et al*. 2025; Zanno & Napoli 2025). Within each of these major groups relative placements of specific genera are often somewhat different than in the past efforts, these items are not a primary focus of this study.

Focusing in on the results that are most pertinent herein, the placement of the Appalachian *Appalachiosaurus* and long armed *Dryptosaurus* holotypes at the base of eutyrannosaurs as per the usual pattern. The new results agree with the positions of CMNH 7541, BMRP 2002.4.1, HRS 08, LACM 28471 and NCSM 40000 outside short armed tyrannosaurids as per Longrich & Saitta (2024) and Zanno & Napoli (2025), contrary to the ETRH. This pattern obviously works against any one of these specimens being juvenile *Tyrannosauru*s. With the novel characters including those related to the limbs of the baso-eutyrannosaurs incorporated, and LACM 28471 processed on its own, it scores as outside *Nanotyrannus* (contra Zanno & Napoli 2025). The latter as the holotype skull alone is basal relative to BMRP 2002.4.1, HRS 08, and NCSM 40000, with the last allied with similarly sharp snouted LACM 28471 (contra Zanno & Napoli 2025). *Nanotyrannus* does not share 5 synapomorphies with the clade containing BMRP 2002.4.1, LACM 28471 and NCMS 40000, and additionally does not share 4 synapomorphies with the Larsonventorini. The siting of CMNH 7541 in a clade with BMRP 2002.4.1, HRS 08, LACM 28471 and NCSM 40000 may be an artifact of the absence of postcrania with the *Nanotyrannus* holotype as further discussed below, it may be a short legged dryptosaur. BMRP 2002.4.1 and HRS 08 appear to form a clade.

Forming their own clade are sharp snouted *Alioramus* and *Qianzhousaurus* (Lu *et al*. 2014; Brusatte & Carr 2016; Carr *et al*. 2017; Dalman *et al*. 2024; Longrich & Saitta 2024; Zanno & Napoli 2025). Our results have them at the base of the clade that includes tyrannosaurids, similar to Carr *et al*. (2017), Dalman *et al*. (2024 and Longrich & Saitta (2024), the other studies place them well up in the family. If the new results are correct then the clade may be a sister family to tyrannosaurids. These results and alternative possibilities cannot be currently further tested via the nature of the Asian graciles forelimbs because of those appendages being absent, but their not being especially close relatives of the North American baso-eutyrannosaurs is indicated by features of their skulls and dentition. The results of the New Technology search with standard setting search equivalent to the strict consensus tree position *T.*? *mcraeensis* directly basal to TT-zone *Tyrannosaurus*, which is agreeable with its being at the base of the latter genus, or outside it on gradistic grounds (as noted above). In the traditional search the most parsimonious results have *T*.? *macraeenis* basal to the rest of the tyrannosaurin including the Asian genera, in which case it is not a *Tyrannosaurus* according to standard definitions of the species (but see comments below). Highly robust, spindle bossed *T. imperator* comes out as basal to later *T. rex* and *T. regina* with their derived single slender anterior dentary teeth, the exceptional gracility and lack of prominent orbital boss indicates *T. regina* is not within contemporary *T. rex* (Paul *et al*. 2022; Paul 2025a). Because the *Tyrannosaurus* species do not fall into a straightforward stratigraphic series their evolution via simple anagenesis is ruled out, the details of how *T. regina* and *T. rex* are interrelated is not clear.

### The actual juvenile *Tyrannosaurus* fossils

The study of *Tyrannosaurus* ontogeny has been seriously limited -- and the survival of the ETRH seemingly enhanced - by the absence of a fairly complete skull and skeleton below a subadult femur length of substantially below 1250 mm and 6 tonnes (Fig. 9A-F) if the CMNH, BMRP, HRS, LACM and NCSM fossils are excluded from the genus; unlike the substantial growth series for *Gorgosaurus* (Fig. 11 in Paul, 2025a) and *Tarbosaurus* (taxon entries in Paul 2024,a,b). But that does not mean there are not juvenile tyrant lizard fossils. As a collective USNM 6183, UCRC V1, LACM 23845, BHI 6439, KU 156375, UCMP 84133, MOR 1189, Baby Bob and DMNS Teen Rex – the last being the most complete among the collection including the skull -- possess the youthful attributes anticipatory of final adult form expected in juveniles of giant adult *Tyrannosaurus*. Among these are low maxillary and dentary tooth counts of 12–13 fairly robust teeth, anterior maxilla and antorbital fossa fairly deep, frontal not elongated, dentary robust, prominent lateral dentary groove lacking, femur about the same length as the tibia which is normal in juvenile albertosaurs and tarbosaurs of similar dimensions (Fig. 2B-D; Schmerge & Rothschild 2016a,b; Longrich & Saitta 2024, Paul 2025a; Raun *et al*. 2025a; Zanno & Napoli 2025). These remarkable remains show little variation between them. Zanno & Napoli (2025) placed KU 156375 in *N. lancensis*, but it has a *Tyrannosaurus* grade low tooth count (Fig. 2C; Burnham *et al*. 2018). The true pre/teen tyrant lizard fossils further show that basic attributes of the mighty tyrant lizard were already in place early in ontogeny, the juveniles being miniature proto *Tyrannosaurus*, stoutish in their few teeth and their bones, well on their way to becoming yet more so when mature. Dramatic changes in form with growth to adapt to ontogenetic shifts in niche partitioning did not happen beyond that seen in growing albertosaurs and, even more telling, in the fellow tyrannosaurin tarbosaurs in which the young are similar in core attributes to their adult morphology (Figs. 3F, 5A-H; Longrich & Saitta 2024; p. 163 in Paul, 2024b; Figs. 11 in Paul, 2025a, Yun *et al*. 2025; Raun *et al*. 2025a; Zanno & Napoli 2025). To put it another way, the amniote and tyrannosaurin growth pattern, and what is known of *Tyrannosaurus* juvenile fossils, mean that if some of those specimens were close to complete, the resulting series of same scale profile-skeletal growth series should be much like that of its close tyrannosaurin tarbosaur kin (as restored for Baby Bob in Figs. 3E, 9H). Not required, therefore, is the drastic divergences from regular amniote/diapsid growth courses forced by the ETRH (Table 1). Those divergences consisting of an exceptional degree of intraspecific anatomical variation within the juvenile population, followed by a similarly remarkable reduction in anatomical variation with growth that involves an extraordinary, nonamniote metamorphosis associated with a sudden transformation in morphobiology, a highly atypical diapsid tooth count contraction in variation via reduction – that in giants whose adult jaws are exceptionally long with plenty of space for lots of teeth -- nor irregular growth curves. *Tyrannosaurus* always looked like *Tyrannosaurus* as it matured, its growth was, as expected, fully within the amniote routine as per its sister taxon *Tarbosaurus* (Figs. 3F, 5A-H). There is not actual, positive evidence otherwise. The conservative, parsimonious, null hypothesis cannot be other than these youthful *Tyrannosaurus f*ossils completely remove and entirely contradict the need or desire of the ETRH to force all lesser sized TT-zone tyrannosaur fossils into *Tyrannosaurus*. Also rendered moot is the Voris *et al*. (2025) observation of possible asymmetrical loss of one tooth in an adult *Tyrannosaurus*, since it would not have had many more teeth when young in any case (see Paul 2025a and Zanno & Napoli 2025; nor has the situation in other adult tyrannosaurids been documented to see if it is different from *Tyrannosaurus*).

**FIGURE 9.**
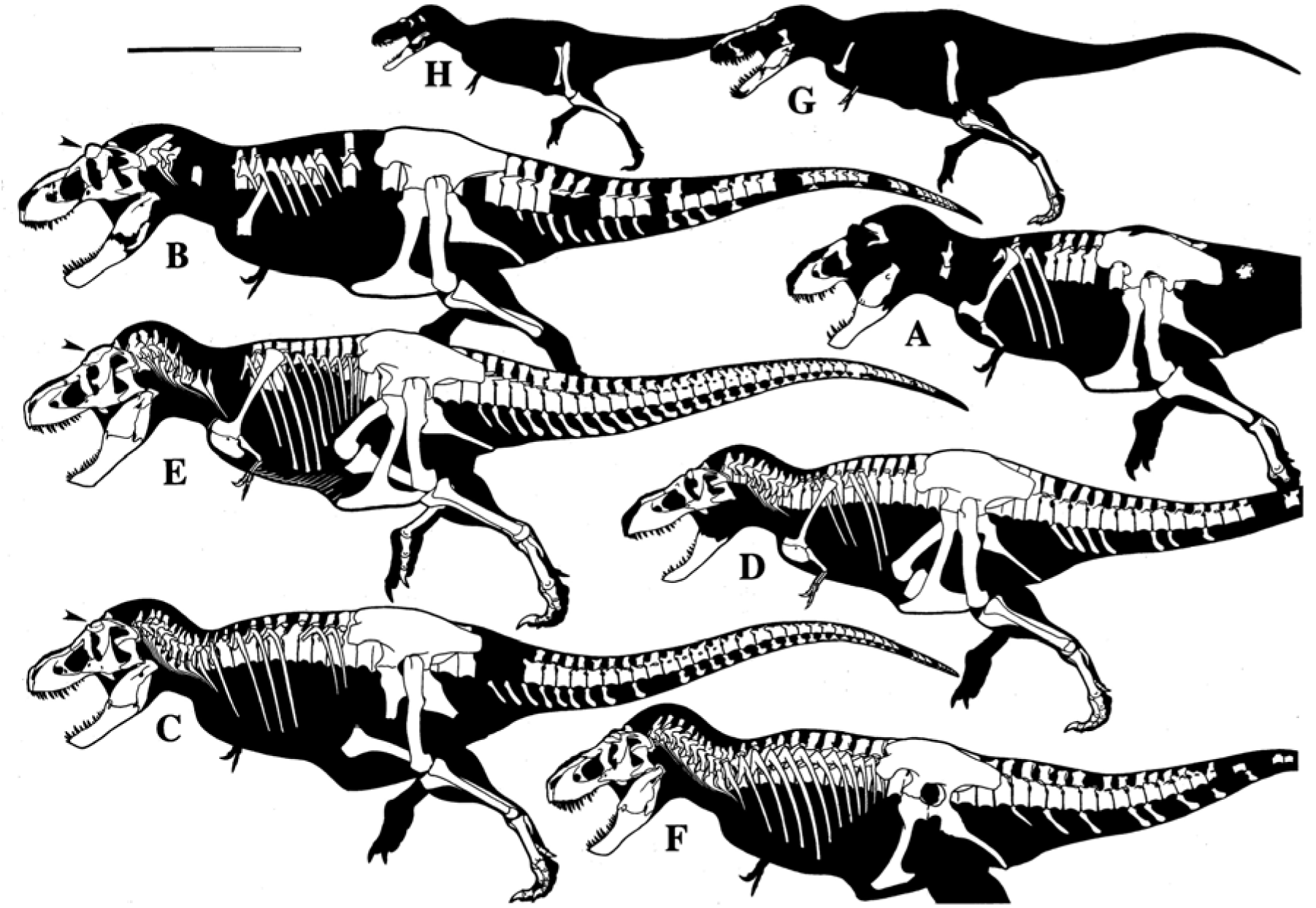
TT-zone adult and juvenile *Tyrannosaurus* known-bone profile-skeletals and skulls to same scale, bar equals 2 m, arrows point to postorbital bosses of large presumed mature males (modified from Fig. 1 in Paul, 2025a). **A** Upper TT-zone *T. rex* holotype CM 9380 (ex AMNH 973) (6.5 tonnes); **B** Upper TT-zone *T. rex* RSM 2523.8 (mature, male? 7.8); **C** Upper TT-zone *Tyrannosaurus regina* NHMAD 2020.00001 (ex BHI 3033, NHMAD S in Paul, 2025a) (mature, male?, 7.5); **D** Upper TT-zone *Tyrannosaurus regina* holotype USNM 555000 (immature? male? 6.1); **E** Lower TT-zone *Tyrannosaurus imperator* holotype FMNH PR2081 (mature, male? 7.8); **F** TT-zone level unknown *Tyrannosaurus incertae sedis* AMNH 5027, preservation of ribs uncertain; **G** upper TT-zone *Tyrannosaurus rex* or *T. regina* LACM 23845 (juvenile, 2); **H** level uncertain *Tyrannosaurus incertae sedis* Baby Bob (juvenile 650 kg).

The fragmentary condition of all these true juveniles, and limited stratigraphic information, precludes placing a number of the specimens in a particular tyrant king species (Paul 2025a). One of the exceptions is KU 156375. From the middle Hell Creek (Burnham *pers. comm.*), that, and its very broad interfenestral pillar, favours assignment of the only known TT-zone juvenile *Tyrannosaurus* maxilla (Fig. 2C) to a late appearing *T*. *imperator* on a tentative basis. From higher in the zone, Baby Bob (Burnham p*ers. comm.*), LACM 23845 and RSM P2347 are candidates for being juvenile *T*. *rex* or *T*. *regina*, further assignments not being practical (Paul 2025a) -- while LACM 23845 had potential as a taxon type if what is known of the fossil had proven highly distinctive from other TT-zone tyrannosaurins (as had been thought by Paul, 1988), it is too fragmentary and immature to be utilized as a type regarding the subtler differences between adult sibling intragenera species of *Tyrannosaurus* (Paul *et al*. 2022, Paul 2025a).

With a number of undoubted juvenile *Tyrannosaurus* fossils on hand, the next question is whether all similar sized TT-zone tyrannosaur remains are also juveniles of the genus, or are so divergent from tyrant lizard juveniles and adults that they must be a different taxon or taxa?

### The subadult and adult non*Tyrannosaurus* TT-zone fossils

As a collective BMRP 2002.4.1 & 2006.4.4 1, CMNH 7541, DDM 344.1, FNMH PR2411, HRS 08, 15001, Jodi, KU 155809, LACM 28471, MOR 6625, NCSM 40000, RSM P2347.1 and TMP 80.16.425 possess attributes that dramatically differentiate and preclude them from being juvenile *Tyrannosaurus* and even tyrannosaurids (Append Fig. 1). Among them are long forelimb elements that in absolute measure are distally about as large or larger than those of adult *Tyrannosaurus*, maxillary and dentary tooth counts exceeding 12 and 14 respectively, teeth more bladed, sharper, more subtriangular anterior maxillae and antorbital fossae, frontal elongated, lack of subnarial foramen, intra quadratojugal-squamosal fenestra, quadratojugal foramen and prominent long lateral dentary groove present, caudal count markedly lower than tyrannosaurins, femur markedly shorter than tibia, metatarsus almost as long as adult *Tyrannosaurus* (Figs. 1, 2E-I, 3C,D, 6C; Table 1; Gilmore 1946; Bakker *et al*. 1988; Currie 2003; Larson 2008, 2013a,b; Schmerge & Rothschild 2016a,b; Longrich & Saitta 2024; Paul 2025a; Zanno & Napoli 2025). The delicate, skull and skeleton of shallow headed, long armed and gracile legged BMRP 2002.4.1, and the even more extremely shallow alioramin style headed NCSM 40000, could hardly be more different from the markedly more massive juvenile of similar overall dimensions of the tyrannosaurin closet to form and function of *Tyrannosaurus* (Fig. 3C,D,F). For the slender forms to have grown into a *Tyrannosaurus* would have required it to hyper morph with growth in a manner common in nonamniotes, but not yet observed in amniotes. With their extra gracile form included exceptionally elongated distal hindlimbs, the likes of NCMA 40000 and BMRP 2002.4.1 are outside the growth series of even the relatively lightly constructed albertosaurs, much less the heftier tyrannosaurins (Fig. 3C,D,G). It is apparent that very lithe American fossils are not the latter or the juveniles of any giant morphotype, or even tyrannosaurids. With their not even being in the same family, it is less anatomically logical to assert that the lithe TT-zone eutyrannosaur nontyrannosaurids are juvenile *Tyrannosaurus* that should have appeared as suggested in Figure 3F, than would be proposing that felid fossils uncovered in the same deposits of ultra-gracile *Acinonyx* fossils were young *Panthera* (Fig. 3H,I). Or using alioramin remains to help restore the growth of tarbosaurs (Fig. 5).

The ETRH proposes that a single taxon was a superpredator that well fulfilled all tyrannosaur predatory niches in its habitat from delicate hatchlings of few kilograms through exceptionally gracile teens up to 7 plus tonne adults. Because the Latter were such massive forms the taxon was effectively precluded from this all-encompassing role. Being on a growth trajectory to mature robusticity the juveniles had to always be on the heftier side of growing juvenile proportions to get to the stout adult condition as they remained in accord with the biological limitations of amniote/reptile growth. Because juvenile *Tyrannosaurus* could not be optimized for tyrannosaur style predation at any stage of immaturity, they were open to competition from mini eutyrannosaurs that were better adapted to be hunters at modest dimensions.

The ETRH scenario being inherently unrealistic, the above specimens being so contrary in their attributes to being juvenile giant tyrannosaurins, and there being little if any actual anatomical or histological positive evidence that they are juvenile *Tyrannosaurus*, arguments otherwise have been unavoidably radical, convoluted ad-hoc attempts to explain away the non*Tyrannosaurus* features the long hands and too many teeth among them, as being in accord with the ETRH because such might not be entirely biologically impossible, even if it is so far out of accord with other amniotes and/or reptiles and tyrannosaurids that it is extremely improbable at best. The few efforts to produce positive ETRH evidence from the specimens have failed. The seemingly *Tyrannosaurus* like anterior frontals of CMNH 7541 (Voris *et al*. 2025) are attributable to distortion and breakage of the fossil (Paul 2025a; Zanno & Napoli 2025). Their being so dramatically different from the authentic juvenile *Tyrannosaurus* fossils of about the same size means the cited specimens cannot be fast-growing tyrant lizards as confirmed by their bone rings. Not only do juveniles not show so much morphological variation within a species or even within a genus, forcing all lesser TT-zone tyrannosaurs into one such taxon requires a sharp reduction in osteological variation with maturity, the opposite of the ontogenetic pattern in organisms. This includes the tooth reduction from as high as 17 to as low as 11 that is not seen in nonmarine, nonbeaked diapsids. The effort by Carr (2020) to selectively use the observations of Brown *et al*. (2015) to defend tooth loss in maturing *Tyrannosaurus* was contrary to the actual results of the latter paper (Longrich & Saitta 2024; Napoli 2024; Paul 2025a; Zanno & Napoli 2025). Supporters of the ETRH have to date evaded addressing these issues by ignoring the existence of the actual *Tyrannosaurus* juveniles (as per Carr 2020; 2025). Nor is sexual dimorphism an explanation for the variation among juveniles. Placing all these variants in *T. rex* would be broadly similar to squeezing all Nemegt tyrannosaurs great and small including *Alioramus* (Fig. 5I) into *T. bataar* when only some of the diverse lessers (Fig. 5D-H) are juveniles of that giant tyrannosaurin (Fig. 5A-C), these being easy to identify because they are visibly younger versions of their parents with which they share common characteristics.

If all the mini TT-zone tyrannosaur specimens were conventionally consistent in morphological form, then the ETRH would be the parsimonious null hypothesis, but that is not the case. Instead, the long forelimbs, manus especially (Figs. 3C, 4D-F) exclude the collection from tyrannosaurids, phylogenetically placing them as basal eutyrannosaurs (Longrich & Saitta 2024; Paul 2025a; Zanno & Napoli 2025). To press all lesser TT-zone tyrannosaurs into *Tyrannosaurus* other than its true juveniles has been an outlier premise that strongly violates the principle null parsimony, and must be rejected. There never has been a paleozoological need for the modern ETRH, and there has never been actual positive evidence for it, the premise resting on perplexing efforts to explain away fatal defects of the hypothesis with increasingly questionably elaborate and thus extraordinary speculations not backed up by the necessary extraordinary evidence, and in some cases are errant (see Longrich & Saitta 2024 and Paul 2025a for further discussion). Instead, the remarkable degree of variation within the TT-zone non*Tyrannosaurus* is itself so high that multiple taxa at at least the species level and probably higher are indicated.

Carr has engaged in an ironic paradox. Although not always direct about it, he continues to be favour *Tarbosaurus* as a junior synonym of *Tyrannosaurus* (Carr 1999, 2020, 2022, 2025). A genus requires limited gradistic variance throughout ontogeny.

Because that is the situation, the congeneric hypothesis is plausible (if the taxa shared a recent, presumably Asian common ancestor which may or may not be correct; Dalman *et al*. 2024; Paul 2025a; also a possible *Tyrannosaurus* in this context is *Zhuchengtyrannus*, particularly if it is intermediate to *Tarbosaurus* and *Tyrannosaurus* as implied by Dalman *et al*. 2024). But if *Tyrannosaurus* featured a growth pattern as far different from the conventional course observed in *Tarbosaurus* that has been proffered by Carr, then they a priori cannot be the same genus. Carr (2025) casually assumes that CM 7541, BMRP 2002.4.1, FMNH PR2411 and LACM 28471 are juvenile *T. rex* without citing the extensive new work of Longrich & Saitta (2024) and Paul (2025a) indicating otherwise, risking contaminating his comparative anatomy of the taxon.

### Growing up not to be *Tyrannosaurus*

Among the errant ETRH speculations was the hypothetical smooth *Tyrannosaurus* growth curve in Figs. 12, 22-25 in Carr (2020) that is predicted by the hypothesis. Although including CMNH 7541, and BMRP 2002.4.1 and 2006.4.4, the plot was not based on the actual, smoothly continuous growth ring sequences from the specimens, Instead the fossils true bone histologies do not conform to the fictional curve. The MTTH in contrast predicts that modest sized baso-eutyrannosaur growth markedly diverged from that of juvenile *Tyrannosaurus* of the same size, as the former slowed down and terminated growth entering maturity, while the latter accelerated mass gains at the middle of their growth cycle. The results of Cullen *et al*. (2020), Woodward *et al*. (2020, 2026), Jevnikar & Zanno (2021), Longrich & Saitta (2024), Paul (2025a) and then Zanno & Napoli (2025) have shown that the BRMP duo experienced a decrease in the pace of growth going into their teens that parsimoniously is most and biologically entirely compatible with, if not requires, their being subadults that would not exceed a tonne. And CMNH 7541 was adult at about half a tonne according to Griffin *et al*. (2025; confirming Gilmore 1946; Bakker *et al*. (1988)), while 600 kg NCMS 40000 was nearly grown up (Zanno & Napoli 2025). Because these results are entirely compatible with and strongly supportive of the MTTH, that the growth data correspondingly affirms the latter means is it is the conventional, parsimonious null conclusion.

Having analyzed the largest yet sample of *Tyrannosaurus* specimens to restore the taxon’s growth, Woodward *et al*. (2026) proceed to speculate that those TT-zone tyrannosaur fossils that fall out of the normal curve of the genus may – if it happens they are in the taxon -- experienced atypical disruptions, or alternative courses, as somehow erratic juveniles of the tyrant lizard. This is done by combining all the remains in a *Tyrannosaurus rex* species complex. This would be like combining the growth data of all Nemegt tyrannosaurs including *Alioramu*s into a *Tarbosaurus bataar* species complex. With the latter venture unlikely to be conducted the question arises if why it was done it was with the TT-zone remains, and its utility. If not for the ETRH having been widely adopted, it is doubtful the *T. rex* species complex would have been proposed, it now coming across as obsolete and improbable thesis in search of a theoretical home. It is a nonparsimonious and therefore weak ad hoc hypothesis that meets no particular current need. There not being any, much less compelling, morphological evidence that any of the slow growth specimens were *Tyrannosaurus* in the first place. Only if the latter were true would it be necessary to consider and perhaps accept their early slowing growth as within *Tyrannosaurus* bounds, and try to produce explanations for the diversion from the taxon’s norm. Ascribing the similar growth of the BMRP pair to similar conditions within their common habitat (Woodward *et al*. 2026) does not explain NCMS 40000 and CMNH 7541 sharing the same attributes, they being from different environments. That the preserved growth of little DDM 35 is within the curve of small *Tyrannosaurus* is not informative because it died before the growth curve divergence of the maturing baso-eutyrannosaurs. Combining the course of maturation from different species to restore growth of genus level is inherently problematic (Paul 2025a), but sibling species of the same adult size have a good chance if not probability of being similar. The fitting of all the actual *Tyrannosaurus* sample into a single coherent curve (Woodward *et al*. 2026) is therefore compatible with the specious *Tyrannosaurus* hypothesis, that neither predicting nor requiring a differentiation between the similar sized sibling species in this factor. Although irregular growth in *Tyrannosaurus* may be biologically plausible, that there is no actual evidence of such within fossils anatomically assignable to the genus or any of its species, leaves the Woodward *et al*. (2020, 2026) hypothesis a paleoosteological stretch that is as entirely unnecessary, as it is inferior to the conservative MTTH null alternative.

## The many lesser-sized TT-zone baso-eutyrannosaur taxa

With it not being possible to fit the majority of mini TT-zone tyrannosaurs into *Tyrannosaurus* or even Tyrannosauridae a number of taxonomic questions ensue. Are they all *N. lancensis*, or do they record a larger number of taxa? If the latter is operative, how many genera and/or species do they represent? Are any of the remains requiring assignment to previously named taxa, or new names? The last question is dependent in part on TT-zone fossils not always being synonymous with the tyrannosaurs of Appalachia. What sub/family or sub/families do the fossils belong to, and do any necessitate new names?

A core feature of Tyrannosauridae is the very small size of the forelimbs, including the manus (Figs. 3F, 9). It follows that any tyrannosaurs that possess larger fore appendages are very probably not tyrannosaurids, instead being more basal eutyrannosaurs (AppendFig. 1). That is assumed to be the case herein for the TT-zone non*Tyrannosaurus* specimens, at least some or all of which were or probably were large handed (Larson 2013; Paul *et al*. 2022; Longrich & Saitta 2024; Paul 2025a). In any case, the focus of this study is on comparative genus and species status over exact phylogenetic position which appears to be undeterminable at this time due to limited data.

With it clear that most non*Tyrannosaurus* lithe specimens are also too different from juvenile and adult *Tyrannosaurus* to be such, it has sometimes been presumed that they themselves all reside on one taxon, presumably *Nanotyrannus* at least at the genus level, if not *N. lancensis* specifically (Larson 2008 2013a,b; Zanno & Napoli 2025). This is uncomfortably similar to placing all giant TT-zone *Tyrannosaurus* in *T. rex*. If the non*Tyrannosaurus* fossils all show a tight osteological similarity, then the every small taxon is *Nanotyrannus lancensis* hypothesis (ESTNLH) is correct. If otherwise, then the multiple small taxa hypothesis (MSTH) is operative, it being a viable possibility in view of the presence of more than one mini tyrannosaur in Asian faunas at least on occasion. That the degree of anatomical diversity being a good deal higher among 1 tonne or less TT-zone tyrannosaurs than within the species of *Tyrannosaurus* which helps show that most of the former are not juveniles of the latter, further favours substantial taxonomic diversity among the lessers. If anything, the greater osteological variation in the smaller tyrannosaurs is likely to reflect even more species and even generic separation that observed in the one genus *Tyrannosaurus*, within which the amount of inconsistency is compatible only with intragenera sibling species. The MSTH is correspondingly highly viable. The next question is whether there is substantial confirmatory evidence in its favor.

Visual examination of the profile cranials (Figs. 1, 2E-I) and skeletals (Fig. 3C,D) of the TT-zone baso-eutyrannosaurs readily reveals the manifest morphological divergences between them that is detailed in the systematic diagnoses. There is more differentiation between the skulls than seen in all three adult *Tyrannosaurus* species skulls combined (Fig. 7F-J). Nor do multispecific *Allosaurus* (Fig. 7A,B), or *Daspletosaurus* (Fig. 7C-E), contain such variation. Among the complete TT-zone baso-eutyrannosaur crania, that of NCSM 40000 is particularly and strikingly distinctive in its low profile, long, shallow, pointed rostrum including a pointed anterior end of the maxillary fossa and fenestra, shallow dentary with an upcurved, chinless tip, reduced posterior mandible complex, and lots of teeth. Also distinctive are the peculiar proportions of its small skulled skeleton (Fig. 3C). There is more divergence in these fossils than is seen among the likes of *Corythosaurus* and *Lambeosaurus* (SupplFig. 4 in Paul 2025a, also 2024a), *Centrosaurus* and *Styracosaurus* (Paul 2024a), within *Canis, Panthera, Metarhinus* (SupplFigs. 2, 3 in Paul 2025a), and is broadly similar to or exceeds that dividing sister genera such as *Ceratotherium* and *Diceros*, or *Plesippus* and *Equus*. Placement of NCSM 40000 in the same genus as either CMNH 7541 or BRMP 2002.4.1 is not viable on basic gradistic grounds.

Within the context of the baso-eutyrannosaur gestalt, the elegantly gracile and delicate NCSM 40000 versus tall, stout and blockish CMNH 7541 could hardly be more divergent in overall configuration and numerous details (Figs. 1A,C, 2E,H). Those include the much more prominent dorsal crests of the 7451 temporal box, a conspicuous ventral jugal boss, the promaxillary fenestra not visible from the side, larger quadratojugal-quadrate fenestra, more horizontal paroccipital process, larger orbit, dentary chin, less teeth and other items cited in the diagnoses. The differences are so dramatic that they indicate major functional differences in predatory actions, and visual display at the aft skull roof. Being well distinct in phylogenetic scoring and positioning (AppendFig. 1), it is questionable whether such can be contained in a subfamily, it is well beyond the species or genus norm. What NCSM 40000 osteologically is easily most similar to the perhaps slightly smaller *S. molnari* holotype (Figs. 2E,F; Longrich & Saitta 2024; Paul 2025a), with them phylogentically plotting as sister taxa (AppendFig. 1). Being only the rostrum and anterior dentary, LACM 28471 is a quite poor holotype bordering on nomen dubium status (which Zanno & Napoli 2025 think it is). Placing other specimens in *S. molnari* or *Stygivenator* risks rendering the genus and species wastebasket taxa. Even so, there is a sufficient diagnostic differentiation between LACM 28471and NCSM 40000 to assess if they appear to be the same taxon or not. They share the same unusually sharp pointed anterior maxillae and antorbital fossae, and upcurved, chinless dentary tips not present in other TT-zone baso-eutyrannosaurs (Fig. 2G-I). These features readily exclude both of them from the squarish nosed *Nanotyrannus* anatomical mold, barring them from inclusion in that genus (contra Zanno & Napoli 2025). It follows that, regardless of its qualifications as a type, LACM 28471 should not be subsumed into *Nanotyrannus* without presenting strong comparative evidence of its taxonomic compatibility gradistic and otherwise, and the specimen consequently needs to be scored separately in phylogenetic analyses. *Stygivenator*’s enlarged, hooked teeth (Fig. 2F,H) render its placement in *Nanotyrannus* all the more gradistically untenable (ditto). The distinctive shared features do indicate that NCSM 40000 and LACM 28471 are closely related members of the same distinct subclade, and Paul (2025a) tentatively placed them in the same genus. There are, however, marked differences between the two that further indicate they are not intragenera sibling species. In 28471 just five tooth sockets are anterior to the anterior end of the antorbital fossa, for 40000 it is eight. The snaggly maxillary teeth of *S. molnari* are much larger, more recurved, and fairly stout. NCSM 40000 sports two small incisiform teeth at the anterior end of the maxilla, LACM 28471 one. The promaxillary fenestra is laterally visible in one but not the other. Unusual for baso-eutyrannosaurs, LACM 28471 lacks the dentary groove that is so prominent in NCSM 40000 and other TT-zone baso-eutyrannosaurs. These differences are too extensive and major to be contained in a dinosaur species, all the more so since sexual dimorphism has not been shown to exist on such a scale in a dinosaur taxon, including predatory (as noted in Paul *et al*. 2022; Paul 2025a). Nor are such extensive divergences present within tyrannosaur genera, matching or exceeding for example that between *Albertosaurus* and *Gorgosaurus* or *Tyrannosaurus* and *Tarbosaurus* (Fig. 2 in Paul 2025a), or within *Daspletosaurus* (Fig. 7C-E) or *Tyrannosaurus* (Fig. 7F-J), as well as *Allosaurus* (Fig. 7A,B). In any case, assigning the superb NCSM 40000 fossil to the at least near nomen dubium that is the fragmentary LACM 28471 is itself problematic, all the more so because if the latter specimen were more complete it likely would show yet more differences with NCSM 40000. And the possibility that there is a substantial time difference between the fossils is itself systematically significant. Nor is it plausible to assign the extreme form that is NCSM 40000 with its low, stretched snouted, small orbit, exceptionally high tooth count and other items to any other known TT-zone taxon on grade grounds alone. As a result – contrary to Longrich and Saitta (2024), Paul (2025a) and Zanno & Napoli (2025) -- NCSM 40000 requires its own genus and species, *Larsonvenator elegans*. Whether it and *Stygivenator molnari* were contemporaries, or separated by time, is not yet certain; the latter is more probable than not. Being diagnostic, *Stygivenator molnari* is retained for the time being at least. These taxa are sufficiently allied with one another, and distinctive from other TT-zone baso-eutyrannosaurs – the exceptionally acutely sharp anterior maxilla being a distinctive apomorphy relative to other late Maastrichtian North American baso-eutyrannosaurs (that were perhaps paralleled in Asian tyrannosaurs (Fig. 5I) -- that they can form a subfamily tagged after the much more complete fossil of the two, Larsonvenatorini. That awaits assignment as the family level circumstances are sorted out as discussed below.

Lacking any postcrania, and having suffered significant cranial damage, CMNH 7451 is a deficient holotype bordering on a nomen dubium (most so regarding issues regarding potential *Tyrannosaurus* species and their names, Paul *et al*. 2022; Paul 2025a) - it is somewhat better in this regard than LACM 28471, but not by a lot. The lack of postcrania is an especially critical phylogenetic and taxonomic deficiency of the mini tyrannosaur. Lacking such, it cannot be ruled out that the specimen is a *Dryptosaurus* which has a straight shafted femur that is about the same length as the robust tibia (Figs. 3B, 6A; Carpenter *et al*. 1997; Brusatte *et al*. 2015), or a close relation. That *Nanotyrannus* falls out closest to the Appalachia baso-eutyrannosaurs among the TT-zone specimens (AppendFig. 1) elevates this possibility. In contrast NCSM 40000, BRMP 2002.4.1 and possibly BMRP 2006.4.4 have to be dismissed as dryptosaurs because they have elongated tibiae and metatarsals, and/or more bowed femora (Figs. 3B-D, 6C,D). Assigning these and other specimens to ambiguous *Nanotyrannus* in view of this large data gap is correspondingly problematic and best avoided unless harder evidence indicates otherwise, and the evidence on hand does not indicate otherwise. Likewise, if the maxilla is so distorted that its true shape cannot be determined as Zanno & Napoli (2025) hint, then its taxonomic utility is further undermined, and placing numerus other specimens in the taxon is all the more problematic, risking making *Nanotyrannus* a wastebasket taxon if other specimens are placed in it. So while using the extreme graciles to characterize and diagnose the postcrania of *Nanotyrannus* may hit the mark, it is at significant risk of being seriously errant. Although CMNH 7541 and BRMP 2002.4.1 are less extremely divergent from one another than either is from the larsonvenatorins, the differences are still extensive; of the two the *N. lancensis* skull is deeper and shorter snouted, the orbit is much larger, the jugal is much larger and the quadratojugal and lateral temporal fenestra taller and perhaps shorter fore-and-aft, the teeth are smaller, and there apparently are two more teeth anterior to the anterior end of the antorbital fossa, the mandible is more robust, and the dentary is squarer tipped than BRMP 2002.4.1 (Figs 1A,B, 2G,H). The status of the intra-quadratojugal-squamosal fenestra is not clear in *N. lancensis*, although it is very likely to have had the condition. The prominent ventral jugal cornual boss, a potential visual taxon identification feature similar to that seen in many tyrannosaurids, is distinctive from the much less developed processes of BMRP 2002.4.1, and HRS 08, as well as NCSM 40000. The extensive phylogenetic work has BRMP 2002.4.1 distinct from more basal *Nanotyrannus*, and nearer larsonvenatorins. Zanno & Napoli (2025) are correct that CMNH 7451 and BRMP 2002.4.1 are not the same taxon, which they distinguished at the sibling species level of *N. lancensis* and *N. lethaeus*. But BRMP 2002.4.1 is gradistically and it follows functionally set well between CMNH 7451 and NCSM 40000. And the differences between the first two – again -- exceed those present within other tyrannosaur genera, being gradistically comparable to those that distinguish *Albertosaurus* from *Gorgosaurus,* and exceeding those observed in *Daspletosaurus* or *Tyrannosaurus* (Fig. 7C-J). Or *Allosaurus* (Fig. 7A,B). BRMP 2002.4.1 is therefore the holotype of *Gilmoretyrannus lethaeus*. HRS 08 (Fig. 2I) is more similar to the latter in its modest-sized jugal and quadratojugal than to *N. lacensis*, and is a sister specimen with BRMP 2002.4.1, and is tentatively referred to at least that genus. That the configuration of the HRS 08 antorbital fossa is distinctive may indicate it was a distinct species, but naming it such is not yet warranted. KUVP 155809 is from very low in the Hell Creek (Burnham, *pers. comm.*), whether it is similarly stratigraphically placed with *Nanotyrannus* or *Larsonvenator* is not apparent at this time. Lacking postcrania in one case, nor can it be determined if these form a subfamily. Whether they are such or not, rounder snouted, square jawed *Nanotyrannus* and *Gilmoretyrannus* are easily distinguished and diagnosed at the genus and even higher level from the sharp nosed, chinless larsonvenatorins, BMRP 2002.4.1 and its cohort being differentiated from NCMS 40000 and LACM 28471 by the large sets of details in the differential diagnoses. The difference in that case is at least as great as it is between *Tyrannosaurus* and *Daspletosaurus* (Fig. 7F-J versus C-E). With NCSM 40000 removed from *Nanotyrannus* the diagnoses of the latter genus and its species are significantly altered and more specific from those in Zanno & Napoli (2025). Due to the great morphological variability found in TT-zone baso-eutyrannosaurs, their placement in one genus does not enjoy parsimony or null status.

Zanno & Napoli (2025) focus on cladistics and particular characters to sort out taxonomic assignments, such as NCSM 40000 (and its well-developed anterior palatine pneumatic chamber) and the *S. molnari* holotype being within *N. lancensis* that both are markedly divergent in basic form from. That wpitfir22ithout seeming to consider the overall anatomical configurations of the fossils remains to assess the total gradistic differentiations that is also critical to generic assignments. This study further finds that characters that Zanno & Napoli (2025) consider markers of genera, instead characterize a more general sub/family level, judging from the more limited degrees of variations observed in other Late Cretaceous tyrannosaur genera. Features of this type may include the openings present in baso-eutyrannosaur quadratojugals.

Despite the much smaller fossil sample size, substantially more baso-eutyrannosaur taxa are observed in the TT-zone, including genera, than among the giant tyrannosaurids. The latter being limited to *Tyrannosaurus* in spite of it including significant species grade diversity (Figs. 7F-J,9; Paul *et al*. 2022; Paul 2025a; prosionally Longrich & Saitta 2024; Zanno & Napoli 2025). This does not arise from our propensity to automatically split specimens based on minor individual or subgeneric variations – after all Paul (2025a) places dozens of specimens in *Tyrannosaurus*, and about a dozen of them each in the three named species. The conclusion stems from their simply being more gradistic and functional diversity contained in the limited baso-eutyrannosaur remains than the entirety of the giants’ combined fossil collective. Comparing Figure 1 to Figure 7F-J, the *Tyrannosaurus* crania are more consistent in overall profile and dimensions of the major elements, including the jugal and quadratojugal. Differences in that genus focus on the postorbital bosses which are exceptionally divergent species specific display features of the tyrant lizard (Paul 2025a). Even if all *Tyrannosaurus* specimens are squeezed into just *T. rex*, the much greater differentiation between its contemporary baso-eutyrannosaurs in the same formations indicates they were more taxonomically diverse. As the future sample of baso-eutyrannosaurs gradually expands it can be expected that many of the specimens will be assigned to the designated taxa. In any case, predator diversity being normally higher among smaller examples in a given habitat, it is not surprising this is proving true in the TT-zone.

Aside from their being intrinsically taxonomically necessary, the new names are designed to help redirect researchers from, as has been the habit, of over-focusing on *Nanotyrannus* as being the go to name for all or most of the lesser TT-zone tyrannosaur fossils that are not *Tyrannosaurus*. With the MSTH heavily favoured over the ESTNLH, there are now four diagnosable genera in that collection of remains.

A question is whether any of the TT-zone lesser taxa are synonymous with similar residents of Appalachia. It is very improbable that any late Maastrichtian baso-eutyrannosaur belonged to the same species as late Campanian *Appalachiosaurus montgomeriensis* (Figs. 3A, 6B) it having lived about 11 million years earlier (Carr et al. 2005). Being in the same genus is less improbable, although still a stretch. In any case, the tibia of the holotype is about the same length of the femur (Carr *et al*. 2005), unlike *Gilmoretyrannus* and *Larsonvenator* in which the tibia is markedly longer than the femur (Figs. 3C,D, 6C). This effectively excludes the two new genera from sharing the same genus as the older eastern avepod. Being a contemporary of the TT-zone fauna, and sporting a long manus, *Dryptosaurus aquilunguis* is a candidate for being conspecific or congeneric with western late Maastrichtian baso-eutyrannosaurs. However, the nearly straight femoral shaft, and/or low tibia/femur ratio of ANSP 9995 (Figs. 3B, 6A), indicate larsonvenatorins and *Gilmoretyrannus* (Figs. 3C,D, 6C,D) are not its close relations. Conversely, the exceptionally large *Dryptosaurus* thumb claw, and a more robust forelimb, add to its separation from *Gilmoretyrannus* and *Larsonvenator* (Fig. 4D-F, Appendfig. 1). Whether postcrania absent *Nanotyrannus lancensis* with its skull distinct from other regional tyrannosaurs, and nearly headless ANSP 9995 are the same taxon at least at the subfamily level, and possibly down to species, is up for taxonomic grabs. They share a lateral dentary groove. But the fragmentary nature of ANSP 9995 may qualify it as a nomen dubium for taxonomic purposes while CMNH 7541 is not much better. Further remains of these taxa are required to resolve these issues. As it is, the possibility of the synonymy at the family or below level is additional reason to be cautious about emphasizing *Nanotyrannus* as the central taxon of the non*Tyrannosaurus* TT-zone fossils. And the possibility that some western tyrannosaurs are within an eastern species or genus consequently cannot be ruled out.

The high diversity taxonomic and otherwise of the TT-zone baso-eutyrannosaurs reinforces that these are not the anatomically uniform specimens expected if they were juvenile *Tyrannosaurus*. To put it another way, with CMNH 7541, NCMS 40000, BRMP 2002.4.1, et al. too different from one another to be placed in the same species or even genus as sub/adults, they cannot be placed as juveniles within a united taxon.

### Family troubles

*Dryptosaurus* has its own family Dryptosauridae, the foundation of which is weak in view of the fragmentary specimen it is based upon (Fig. 3B) – a situation that is not likely to improve in the near future if ever because of the limited exposures of the New Egypt Formation. Zanno & Napoli (2025) placed the *Nanotyrannus* species it their own family, Nanotyrannidae. This is plausible if *Nanotyrannus* had elongated distal hindlimbs, but problematic because of the adequacy concerns with the skull only holotype, and because it therefore cannot be ruled out that it is a dryptosaurid at this time. If that ever happens Nanotyrannidae will be subsumed into Dryptosauridae. That possibility has the possibility of being tested by the finding at some point of more complete CMNH 7541 type postcrania in the extensive TT-zone deposits. The ability of phylogenetics to deal with the situation is limited because *Dryptosaurus* cannot be scored for critical cranial characters. And the postcranial attributes of *N. lancensis* cannot be reliably scored either. Our results (AppendFig. 1) suggest but do not establish that *Nanotyrannus* is not in the same family as other TT-zone mini tyrannosaurs. Until the fossil data base is improved, there is no good resolution for the north American baso-eutyrannosaur family problem (as noted by Paul 2025a). It is possible that both families are valid, in which case Larsonvenatorini is likely under Nanotyrannidae. And/or Larsonvenatorini may need to be elevated to sub or full family status. Because *Gilmoretyrannus* is at least gradistically quite distinct from all other genera its higher level status is indeterminate. In view of these currently recalcitrant paleotaxonomic vexations, this analysis does not set up and diagnose baso-eutyrannosaur families, and Larsonvenatorini is a taxonomic floater among baso-eutyrannosaurs at this time. This is similar to some other intrinsically ambiguous dinosaur sub/family conundrums, such those afflicting titanosaurs, as per Utetitaninae (Paul 2025b).

### Where did so many TT-zone baso-eutyrannosaurs come from? And why did they do so well?

In the ETRH, the origin of known TT-zone tyrannosaurs has simply been that of *Tyrannosaurus.* With the MTTH widely accepted, the circumstances were much more complicated in evolutionary-diversity-geographic terms. The TT-zone was inhabited by a host of mini baso-eutyrannosaur taxa of about a tonne or less when adult, dwelling alongside elephant sized tyrannosaurid species. The number of the latter is three in one genus (Paul *et al*. 2025; Paul 2025a, two are named in Longrich & Saitta 2024; number not specified Zanno & Napoli 2025), the lessers number four for a combined seven species, in five genera. In that case, the tyrannosaur diversity within the habitat was on the high side compared to other tyrannosaur dominated faunas. That when the diversity of TT-zone herbivorous ornithischians appears to have been lower than prior in the region (Fowler 2017; Paul 2010, 2016, 2024a,b). But that tendency cannot be used to reliably assess the diversity of the predatory fauna, which can only be achieved via comparative anatomy of the fossils. Because it was common for Mesozoic dinosaur bearing formations and faunas to include more than one giant avepod including tyrannosaurids, multiple species of TT-zone *Tyrannosaurus* is a norm rather than requiring special explanation. It is the multiplicity of the smaller taxa as per the MSTH that may appear atypical among tyrannosaurs, for which there is usually one identified taxon in a given biospace (Brusatte *et al*. 2009, 2016; Lu *et al*. 2014; Brusatte & Carr 2016; Paul 2010, 2016, 2024a,b). But such low counts may represent under sampling and examination of fossils from given strata in the field and collections (Zanno & Napoli 2025). This prospect is boosted by presence of two named mini tyrannosaurs in at least one formation (Lu *et al*. 2014; Mo & Xu 2015; Zheng *et al*. 2024). And more than one avepod in the one tonne area is known in some paleofaunas (Paul 2010, 2016, 2024a,b). That the TT-zone has been exceptionally heavily prospected since the late 1800s is a factor in its exceptionally high observed tyrannosaur diversity. And it is not as though all the TT-zone tyrannosaurs shared the same paleohabitat. The TT-zone consists of seven formations laid down over considerable time and territory. The large number of named taxa appears to reflect discreet sampling of a diverse sets of baso-eutyrannosaur populations over that time and space. Although the details remain obscure, while lower Hell Creek *Nanotyrannus lancensis* and *Larsonvenator elegans* may or may not have lived in the same time and place, neither probably dwelled alongside *Stygivenator molnari* and *Gilmoretyrannus lethaeus*, which in turn may or may have not been within the same given fauna at the same time. There are no cases to date of more than one tyrannosaur taxon being found in the same quarry; shifting territorial ranges at small and large scales differing habitat/niche preferences, and the like may have reduced real world competition among species that are found in the same gross formation level.

That said, a nontypical causal explanation for the systematic diversity anomaly is called for. That appears to be an exceptional regional geological-geographic event that was underway during the later portion of the Maastrichtian. The reunification after tens of millions of years of the North American continent resulted from the emergence of the Laralachia (Paul 2024b, 2025a) land bridge as the interior seaway withdrew (a hypothesis supported by Zanno & Napoli 2025). Basal eutyrannosaurs, at least some bearing an elongated manus, had been evolving in Appalachia. The new route west allowed them to invade Laramidia and intermingle with its short-armed tyrannosaurids. The degree to which late Maastrichtian baso-eutyrannosaur taxa were direct imports from Appalachia, or had evolved into new taxa since crossing the bridge, cannot be resolved with the data on hand. It is possible that not all the invaders were from eastern North America, some may have moved in from Asia via Beringia. Alioramin skulls are in particular similar to those of larsonvenatorins, sharing a distinctive sharp snout and upcurved, chinless anterior dentary (compare Figs. 1C to 5I). The sparsity of information on the dimensions of alioramin forelimbs stands in the way of settling this possibility to the degree that reliable phylogenetic results cannot be obtained, and whether the Asian examples were tyrannosaurids or basal to the family. For these modest-sized predators to make their way across the high latitude bridge would have been easier than for the giant tyrannosaurins. In those circumstances the unusual high diversity is yet better explained by immigration into the TT-zone faunas from multiple directions.

From wherever they came, the resulting intermixing in the TT-zone appears to have gone well for the new residents, with their fossils outnumbering those of juvenile *Tyrannosaurus* by half or more again. The baso-eutyrannosaurs sporting long jaws bearing many bladed teeth, more mobile, much longer arms and hands that approached those of some allosaurids in length (Paul 2024b), and longer legs should have been better adapted for filling the role of medium-sized predators than were relatively clunky juvenile tyrant lizards. The latter being compromise medium-sized predator niche *Tyrannosaurus* hampered by their severely reduced arms. fewer, blunter teeth, and shorter lower legs feet when it came to pursuing and dispatching the smaller game.

Not yet known is whether western tyrannosaurids successfully moved into Appalachia. It is also possible that baso-eutyrannosaurs whose Appalachian ancestors had earlier headed and settled in the west, then underwent speciation and perhaps beyond in their new home, and subsequently returned in part to Appalachia, *Dryptosaurus* being a possible example. Ergo, it cannot be simply assumed that late Maastrichtian eastern region tyrannosaur clan evolved entirely in place, the situation may have been more complicated. Returning to the Amero-Canadian west, having initially resulted of the novel geographic situation of the later Maastrichtian, the atypically high TT-zone tyrannosaur diversity might have proved temporary over deep time. What cannot be known is how baso-eutyrannosaurs versus tyrannosaurids would have fared proceeding beyond 66 MA had the K/Pg extinction not occurred.

### How did grownup *Tyrannosaurus* become gigantic?

Despite apparently being under intense competitive pressure by markedly more numerous baso-eutyrannosaurs, enough juvenile *Tyrannosaurus* managed to grow up to sustain the populations of the genus. Paul *et al*. (2022) proposed that the rapid appearance of multiple species of an elephant sized tyrannosaurid genus in late Maastrichtian western North America, was due to the appearance of the Laralachia land bridge suddenly enormously expanding the resource base for the great predators. The food base expansion then further allowing the coexistence of larger contemporary robust and gracile tyrannosaurid species than the rhino-sized albertosaur/daspletosaur examples present earlier on the narrow Laramidia peninsula. Paul (2025a) noted the apparent previous existence of oversized tyrannosaurids on Laramidia as early as the Campanian ((Stein & Triebold 2013, Dalman *et al*. 2024. The early existence of *“T.” mcraeensis* has since been challenged (Voris et al., 2025), leaving the issue up in the evolutionary air at this time.

### Restoring *Tyrannosaurus* developmental ontogeny

To repeat the caution by Paul (2025a) then reemphasized by Zanno & Napoli (2025), the common practice of using lesser TT-zone baso-eutyrannosaurs as models, the *Gilmoretyrannus lethaeus* holotype most commonly, for restoring the teen years of *Tyrannosaurus* is as errant as using coyotes to do the same with wolves, or bobcats for pumas, and the like (Fig. 3H,I; contra results of Paul 2008; Hutchinson *et al*. 2011; Carr 2020; Woodward *et al*. 2020). Only juvenile specimens that can be confidently assigned to the tyrant lizard should be used to examine and restore its maturation factors such as growth curves and rates and interruption, shifts in locomotion and agility, predatory habits and niche partition, mental acuity, etc. as they achieved adult status (Figs. 2A-D, 3E, 6E-G, 9). Unfortunately, the fragmentary nature of such fossils to date (Figs. 2B-D, 3E, 9G,H, Fig. 29 in Longrich & Saitta 2024) – Teen Rex may fill that gap -- as well as most being nonaccessioned, inhibits such efforts compared to *Gorgosaurus* and *Tarbosaurus* ontogeny. The latter is therefore the most suitable robust tyrannosaurin for this field of analysis at this time.

### The lithe baso-eutyrannosaur taxa were not functional equivalents to one another

Not only are the small eutyrannosaurs of the TT-zone not useable for examining *Tyrannosaurus* ontogeny other than for disparate comparative purposes, it is necessary to not presume they are functionally uniform among themselves. The opposite is true, the divergences in the *Nanotyrannus* and *Larsonvenator* skulls (Fig. 1A,B) are extensive to the degree of being adaptatively extreme. Far from studying them as a functional unit, the baso-eutyrannosaurs need to be analyzed as distinct genera and probably higher in the normal manner, with *Nanotyrannu*s apparently robust headed from purposes of high predation, and *Larsonvenator* as lightly structured for markedly different hunting and feeding modes. The same principle applies to examining growth patterns and other factors -- see where the data leads rather than assume where it will.

## Conclusion

Based on straightforward comparative anatomy and conventional ontogeny, and a large scale phylogenetic matrix, the 1946 Gilmore Hypothesis established that the TT-zone was inhabited by multiple tyrannosaur taxa. This conservative conclusion likely would have remained the established paradigm, especially had it been reinforced by Rozhdestvensky in 1965. In 1999 Carr contended otherwise, based on a critical ontogenetic tooth count error (which may have stemmed in part from an excessive tooth count for juvenile *G. libratus* AMNH 5664 in Bakker *et al*. 1988). Given that mistake, the radical, nonparsimonious speculations necessary to sustain the ETRH, and the presentation of fossils that directly contradict the opinion years ago, it is perplexing why it was such a popular theory. Instead, the fossil and the paleogeographic data show that the TT-zone experienced an incursion of one tonne or less basal eutyrannosaurs of likely eastern heritage that were successfully competing with the resident juvenile *Tyrannosaurus*. In the future tyrannosaurin ontogeny needs to follow the solid principles established in this decade by Longrich and Saitta (2014), Paul (2025a), Zanno & Napoli (2025) and herein. Rather than just one titanic taxon lasting over 1 MA, identifiable TT-zone species and genera from a few hundred kilograms to 8 tonnes over that span currently number seven and five respectively, with none likely to have been extant over that entire period. With the onset of the compelling MTTH papers over the last two years, study of the TT-zone tyrannosaurus has been released and opened to complex alternative possibilities that are inherently difficult to resolve. With the ongoing sudden displacement of the ETRH with the MTTH as the paradigm, the question is not if there were multiple species and genera in the TT-zone – as well as other tyrannosaur habitats -- but how many and at what systematic level. And what it means in paleozoological regards. The already late Gilmore would never know how prescient his 1946 hypothesis was, and we are honored to have named one of the pertinent genera in recognition of his achievement in dinosaurology.

## Appendix 1: Phylogenetic Cladograms and Character Matrix

**APPENDIX FIGURE 1.**
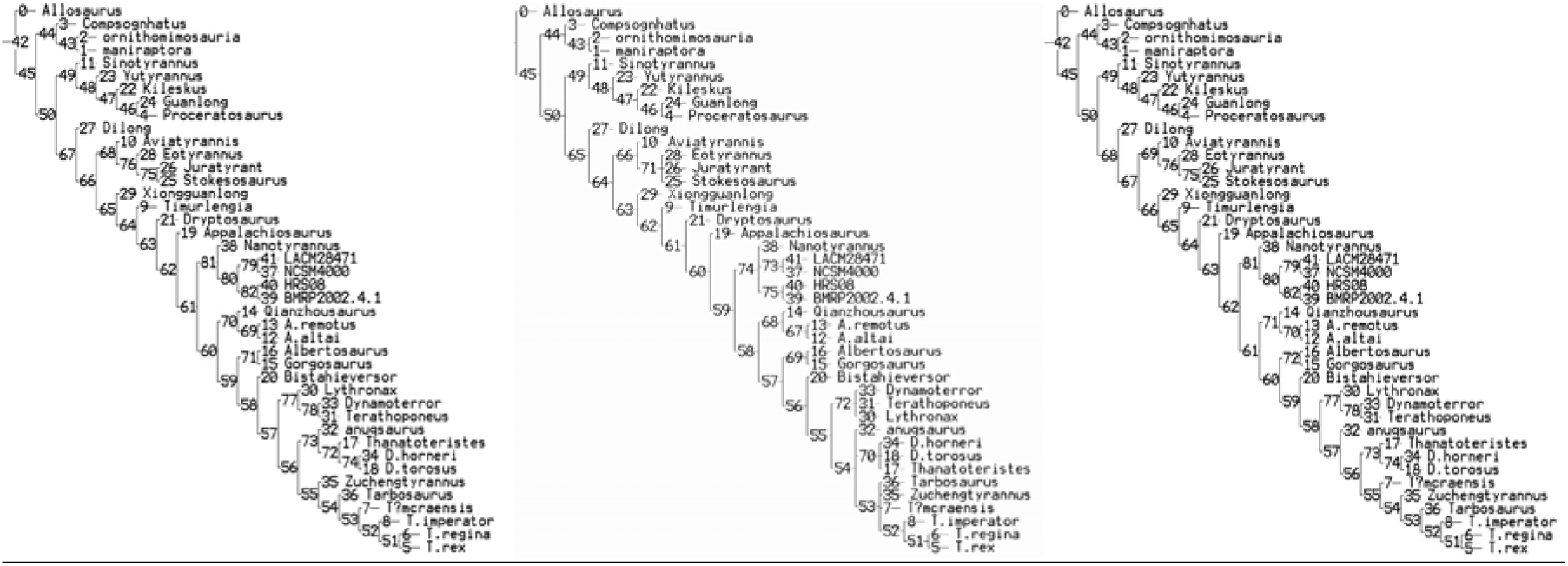
From left to right: New Technology search; traditional search strict consensus tree; traditional search most parsimonious tree. [Figure image will be upgraded for publication]

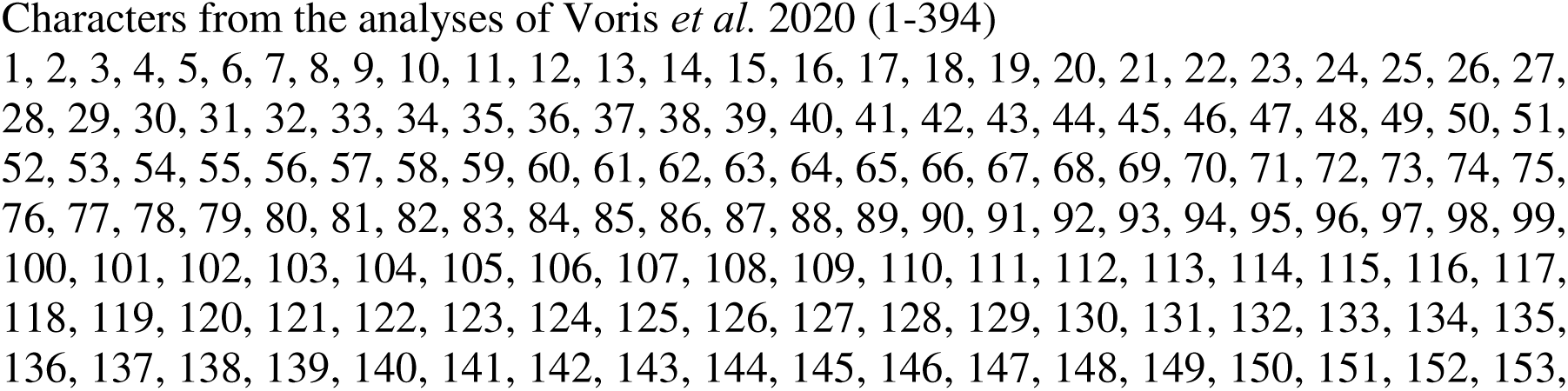
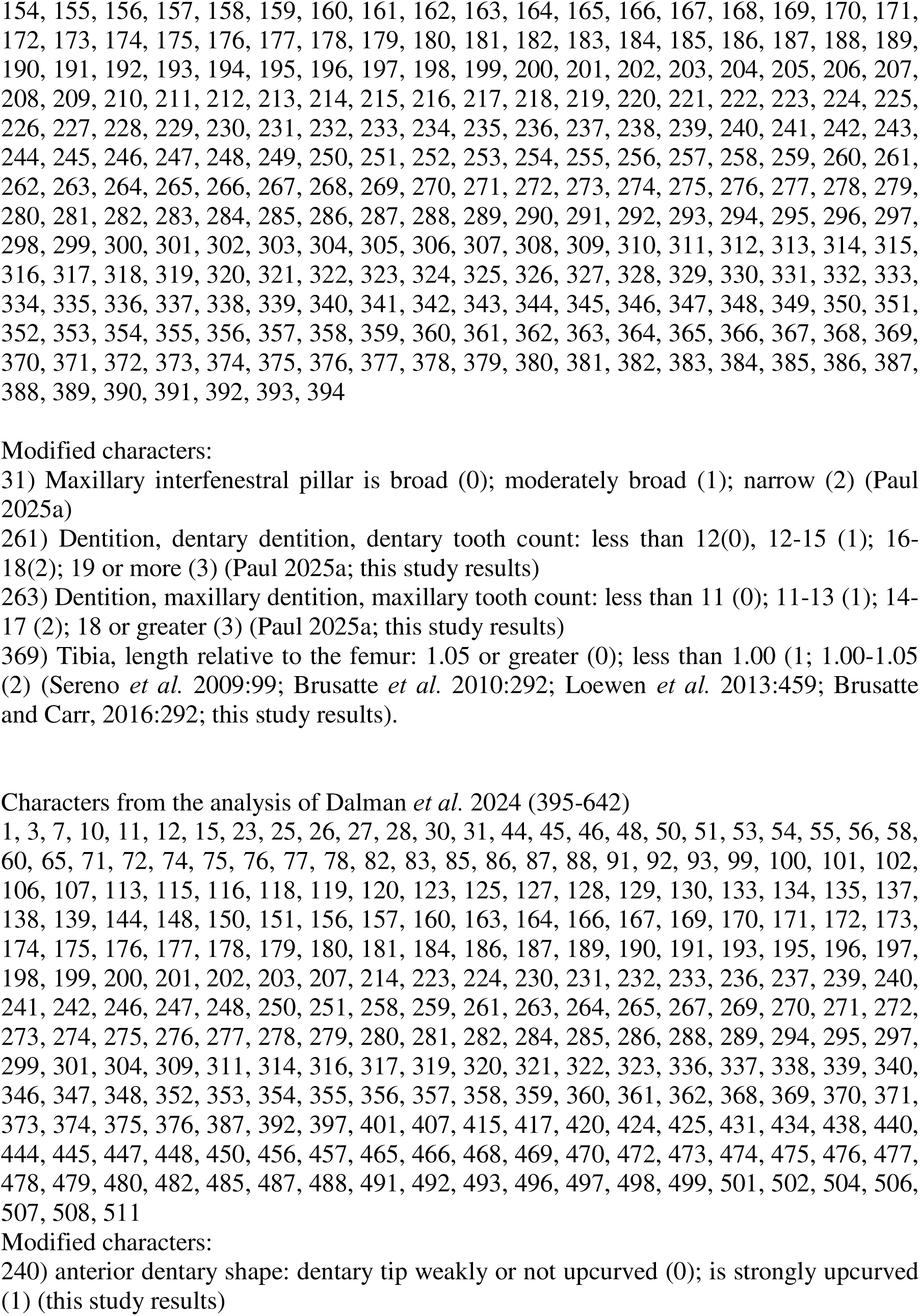
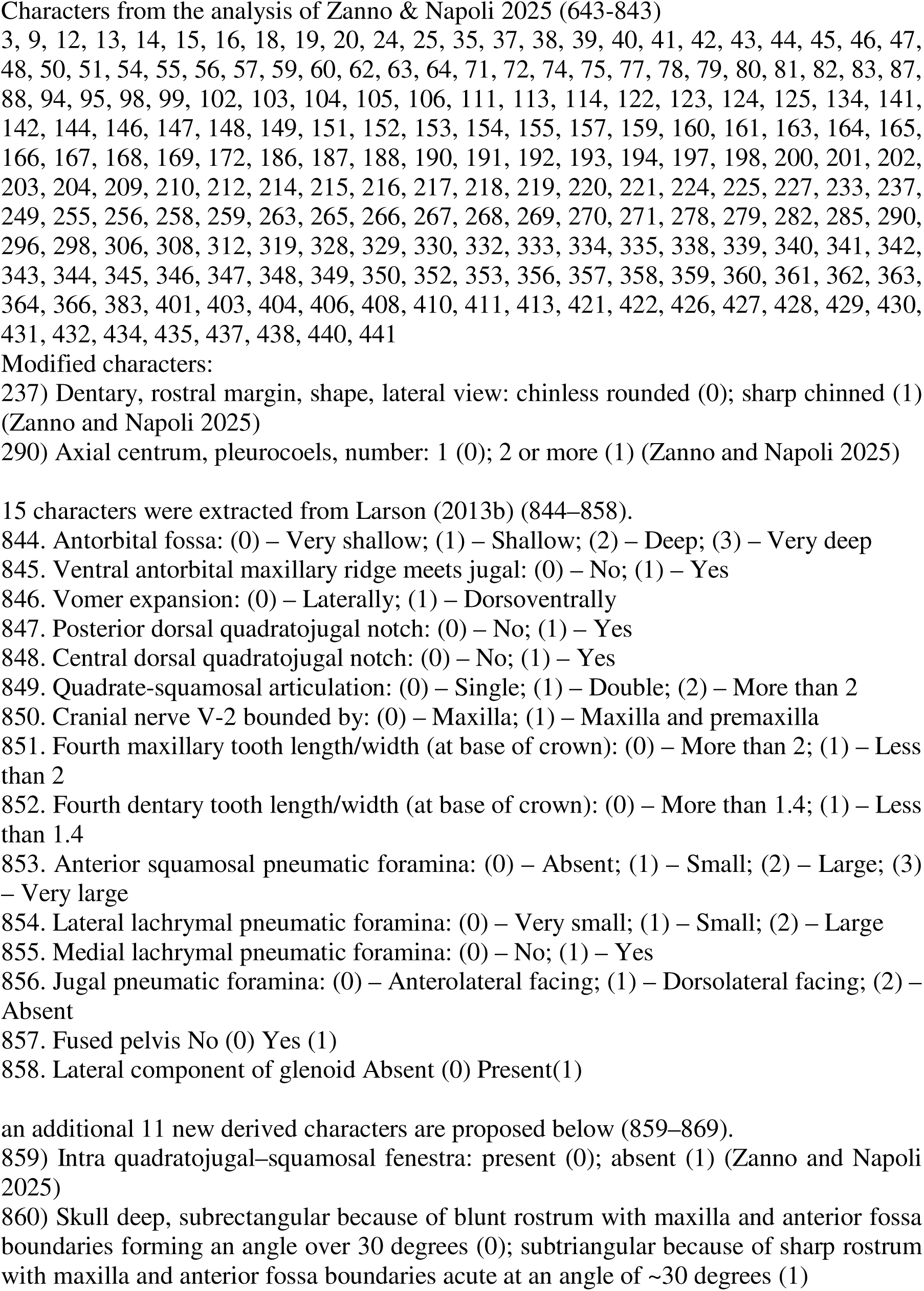
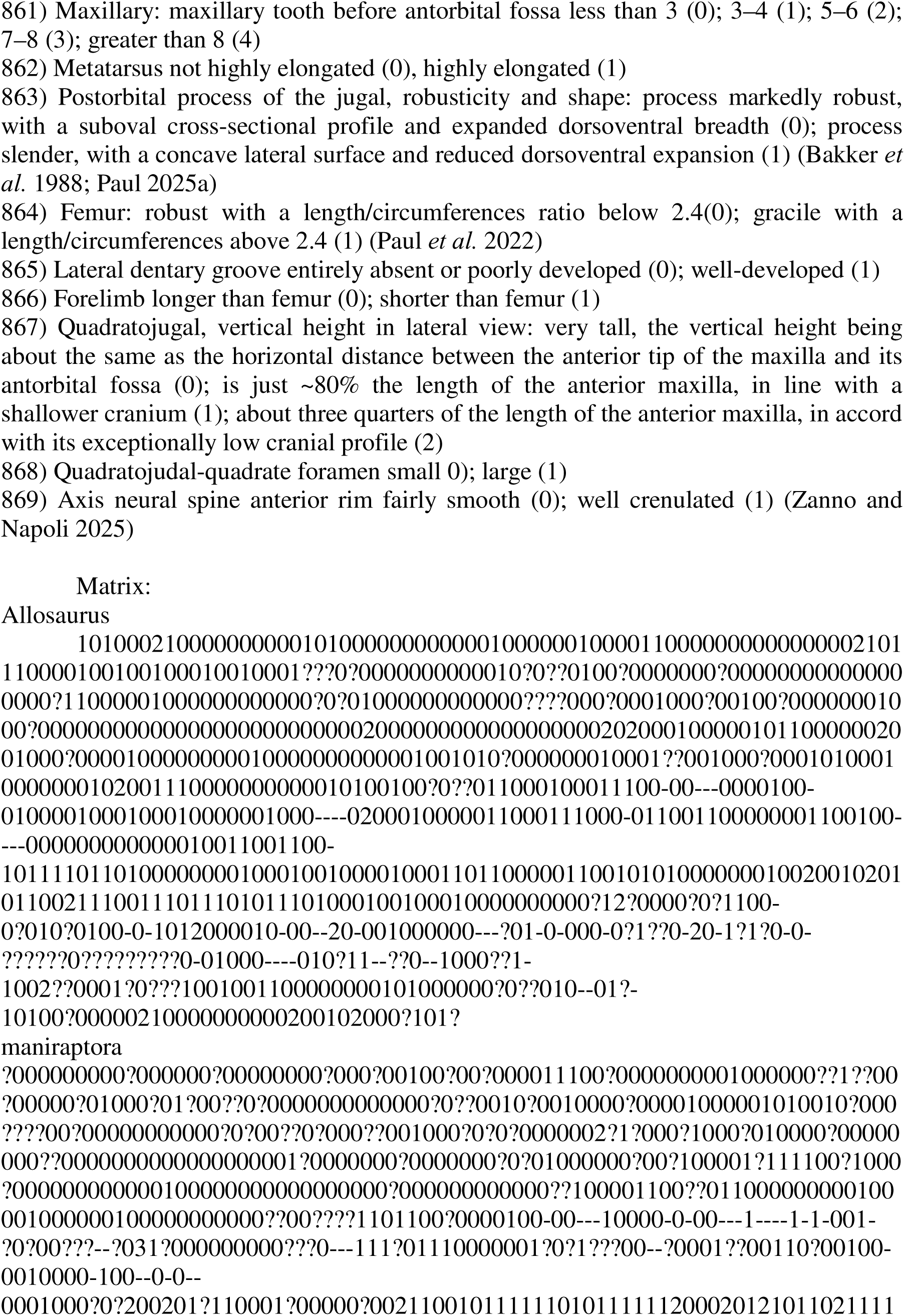
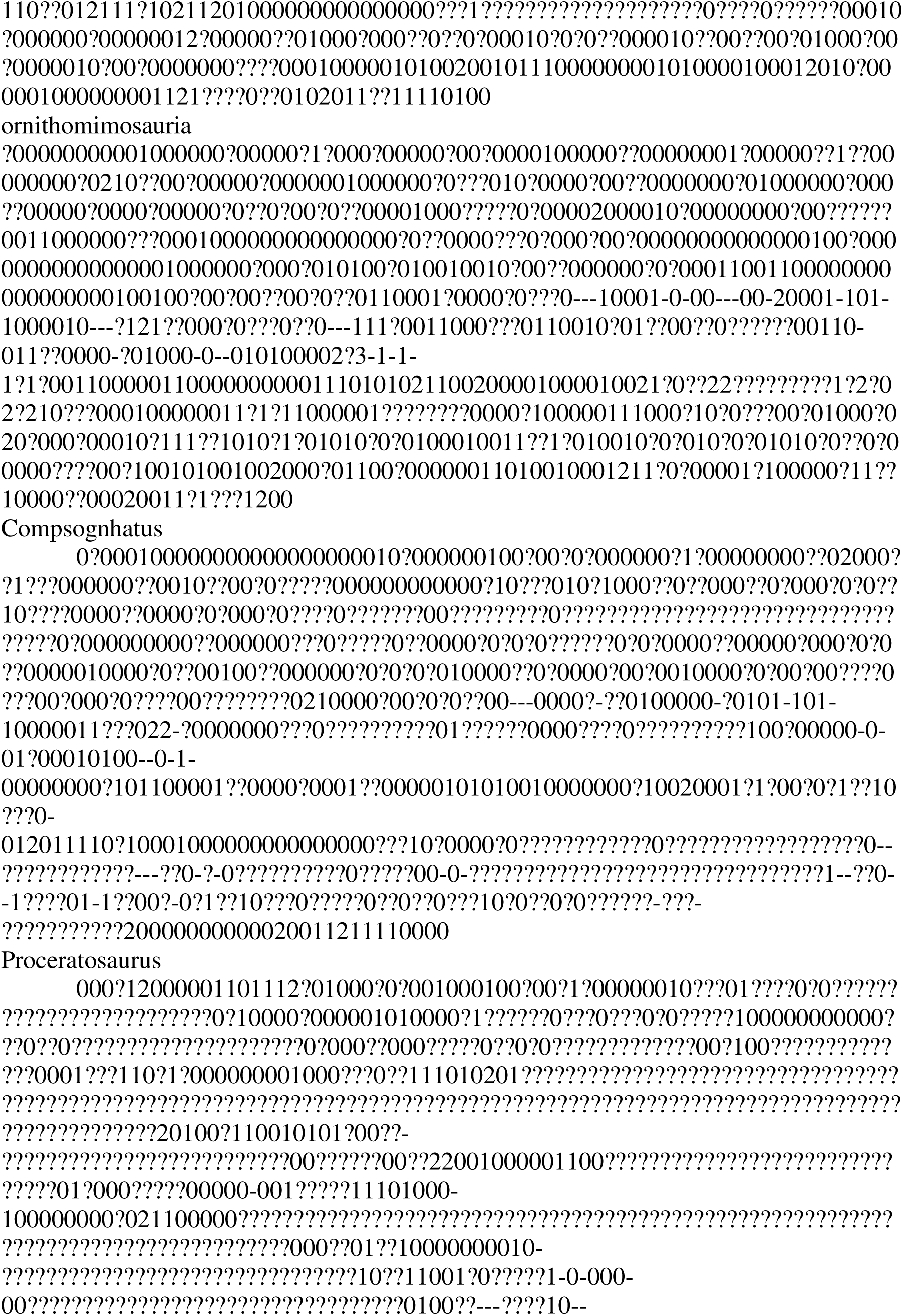
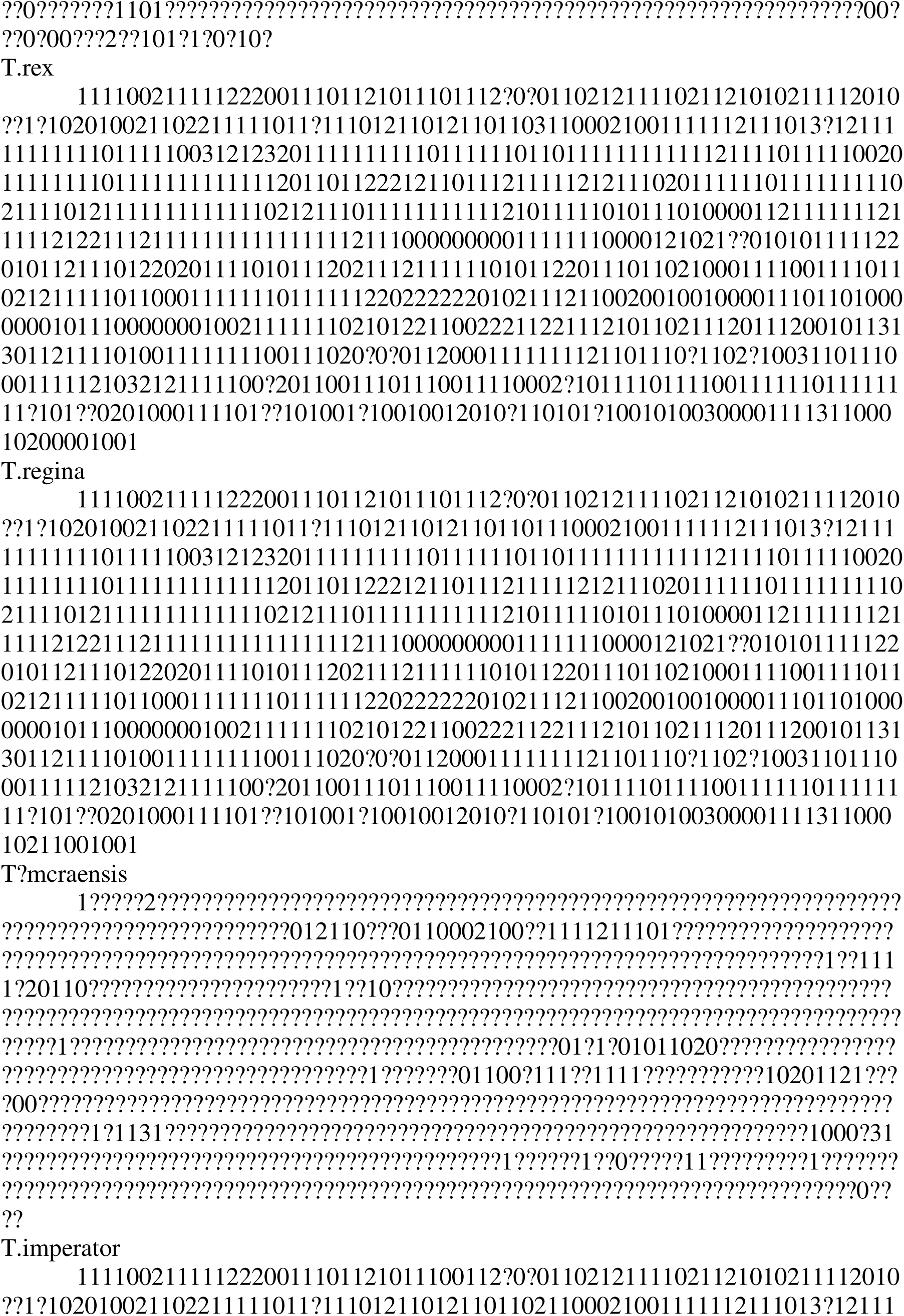
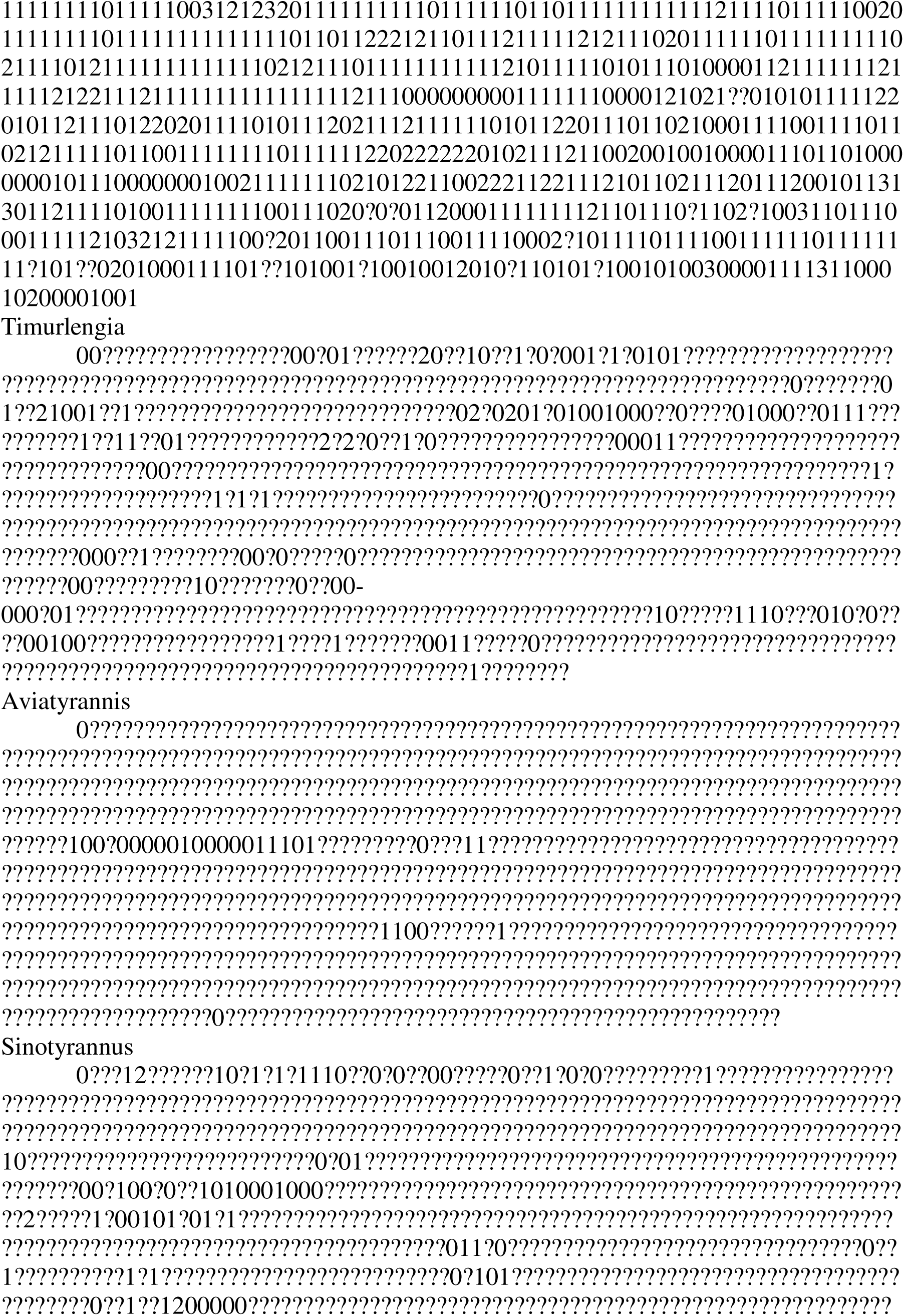
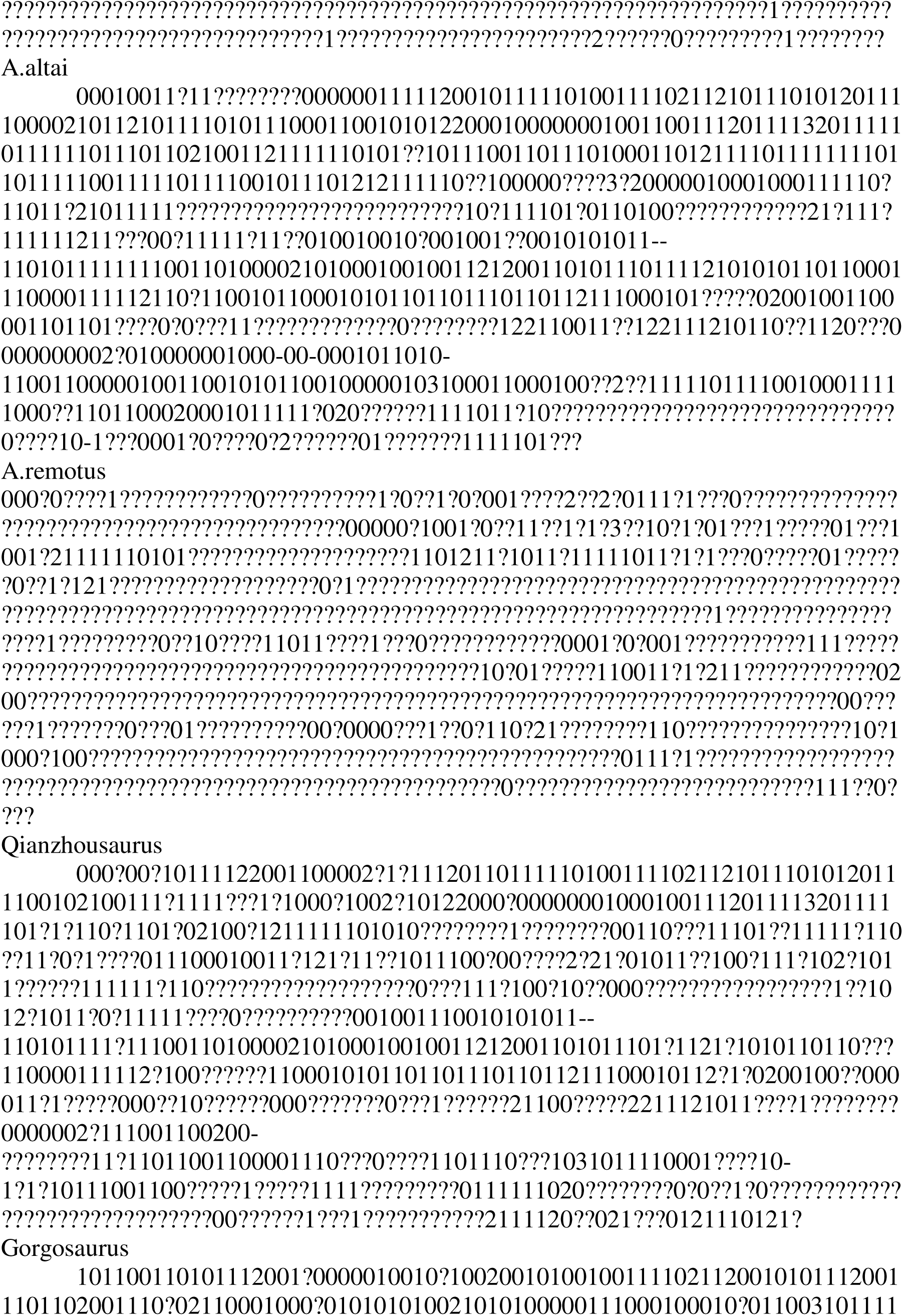
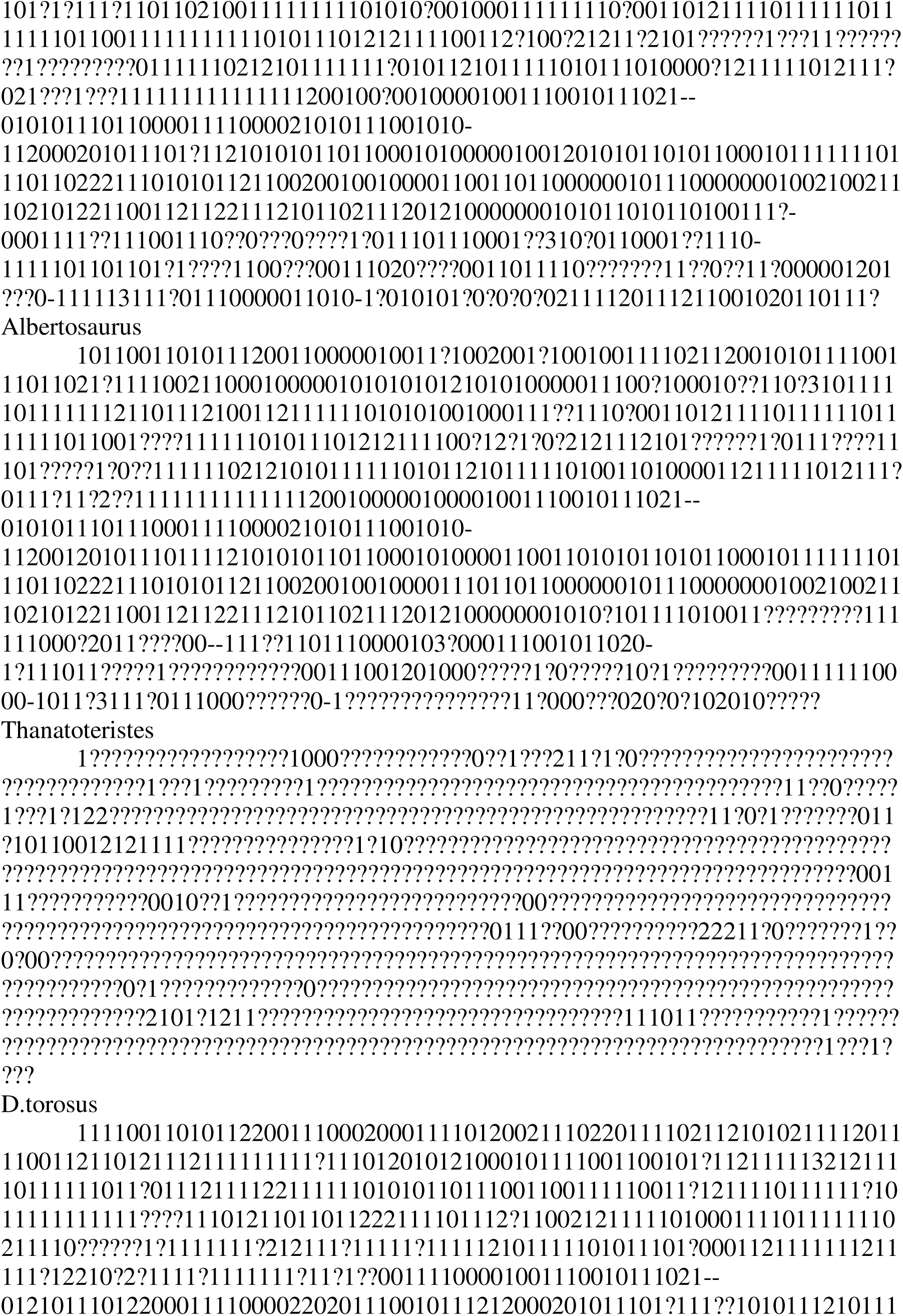
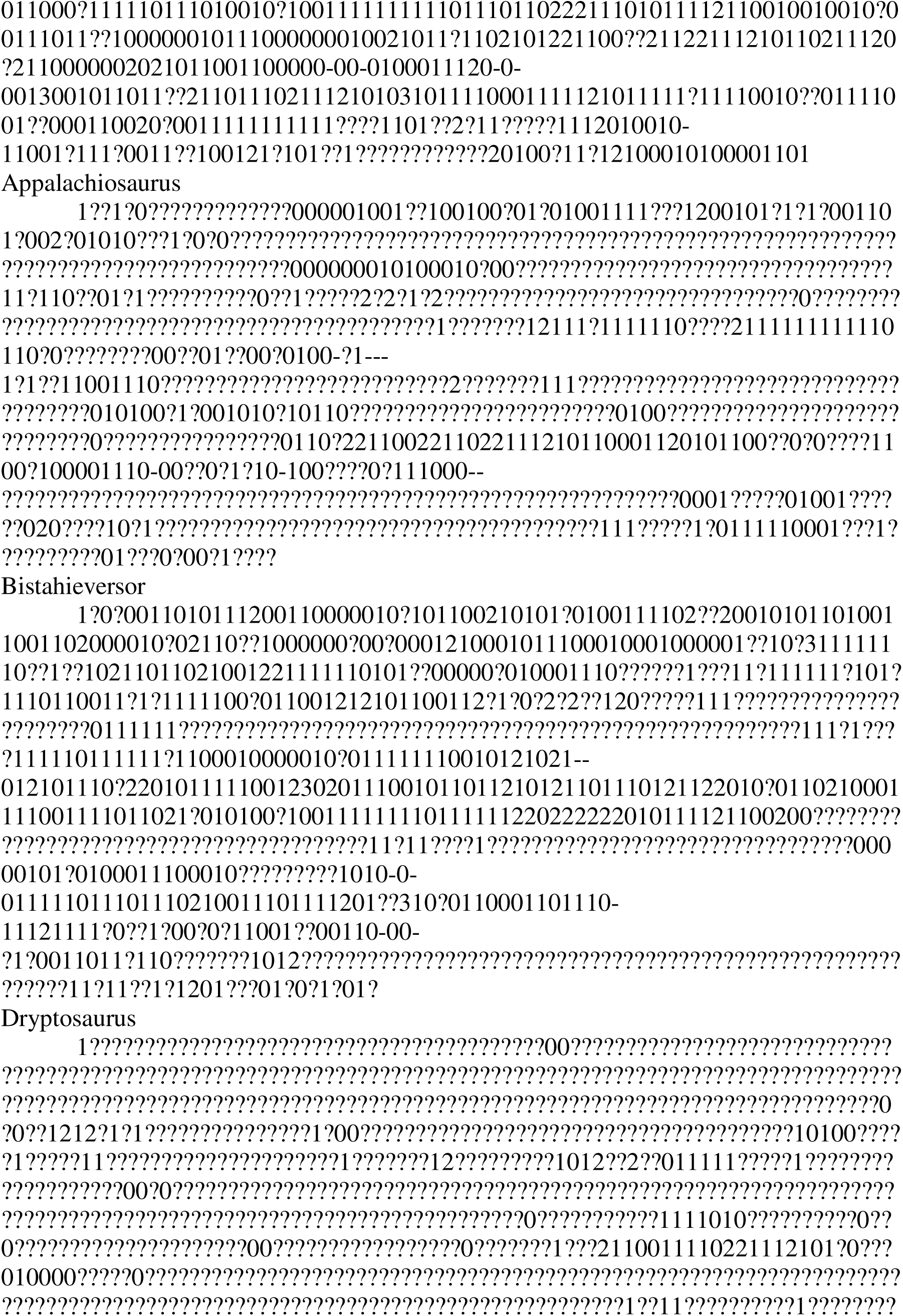
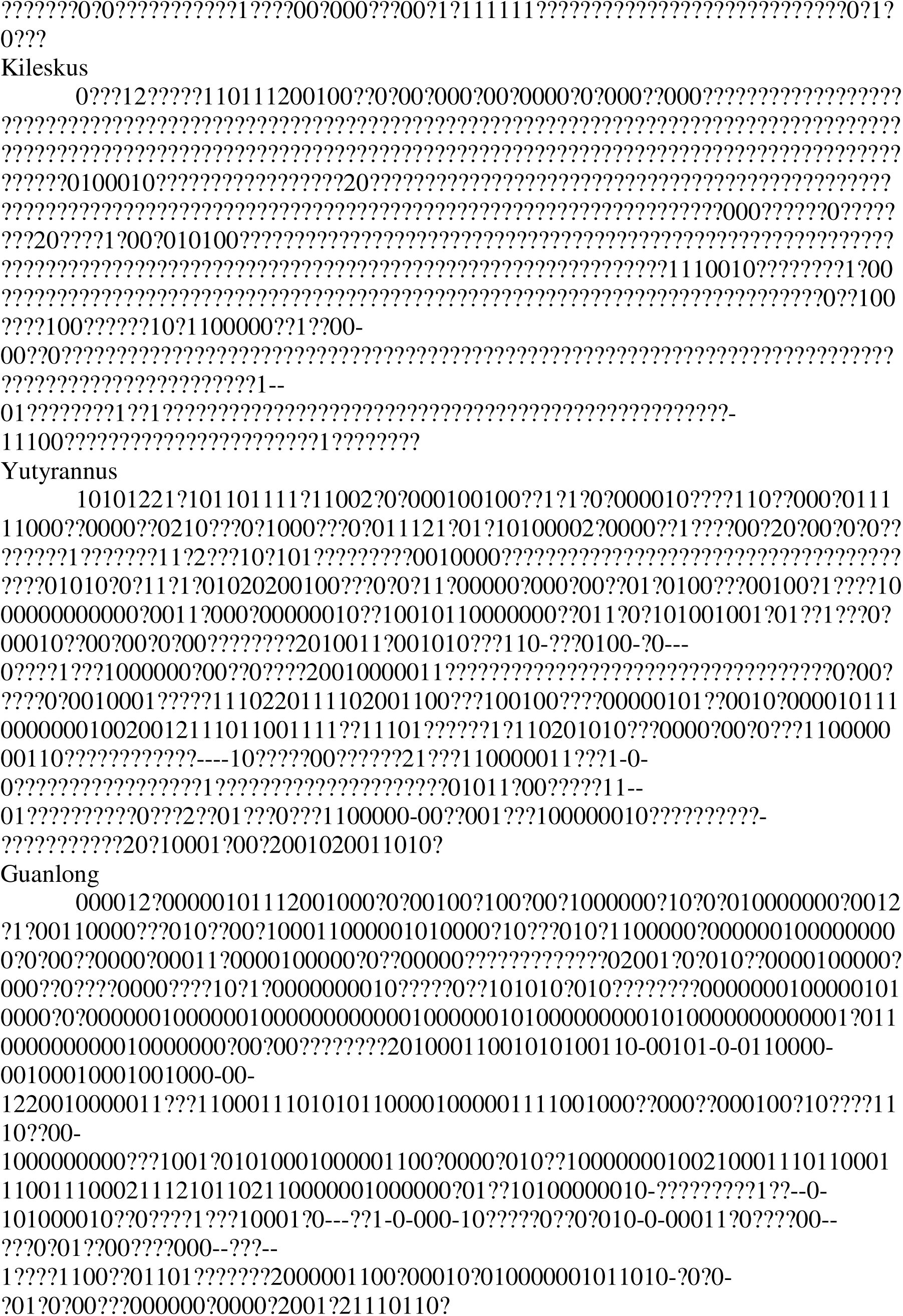
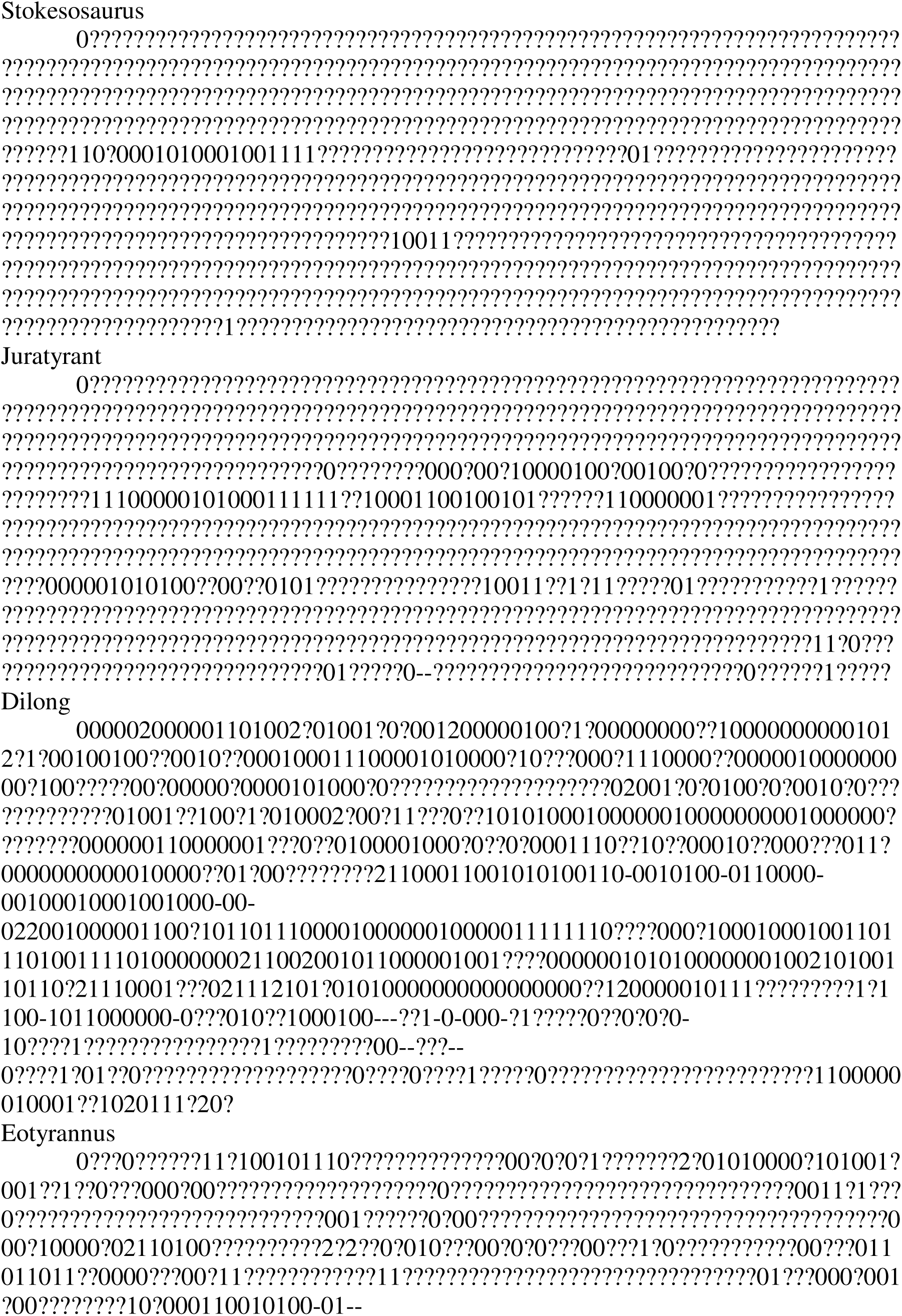
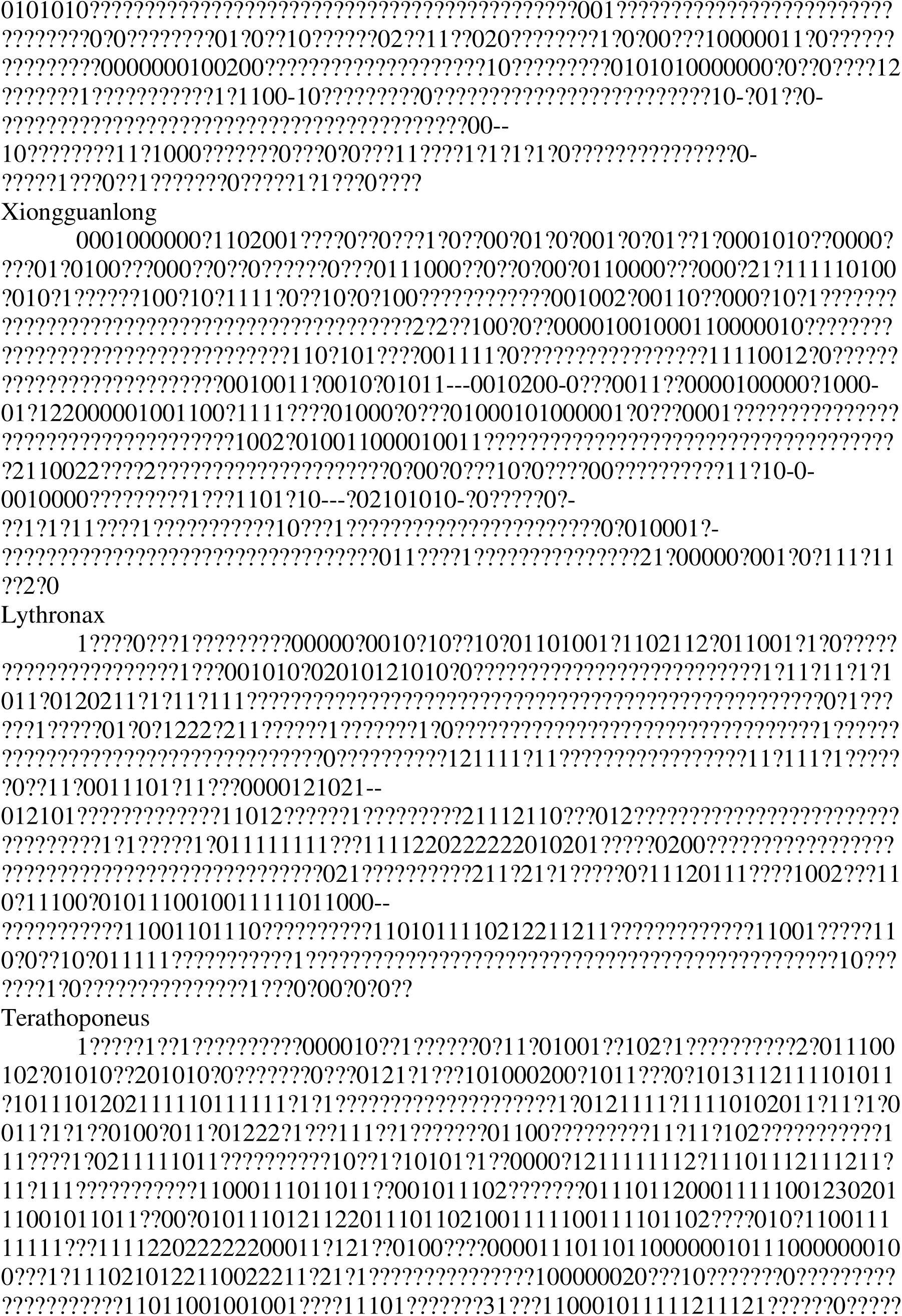
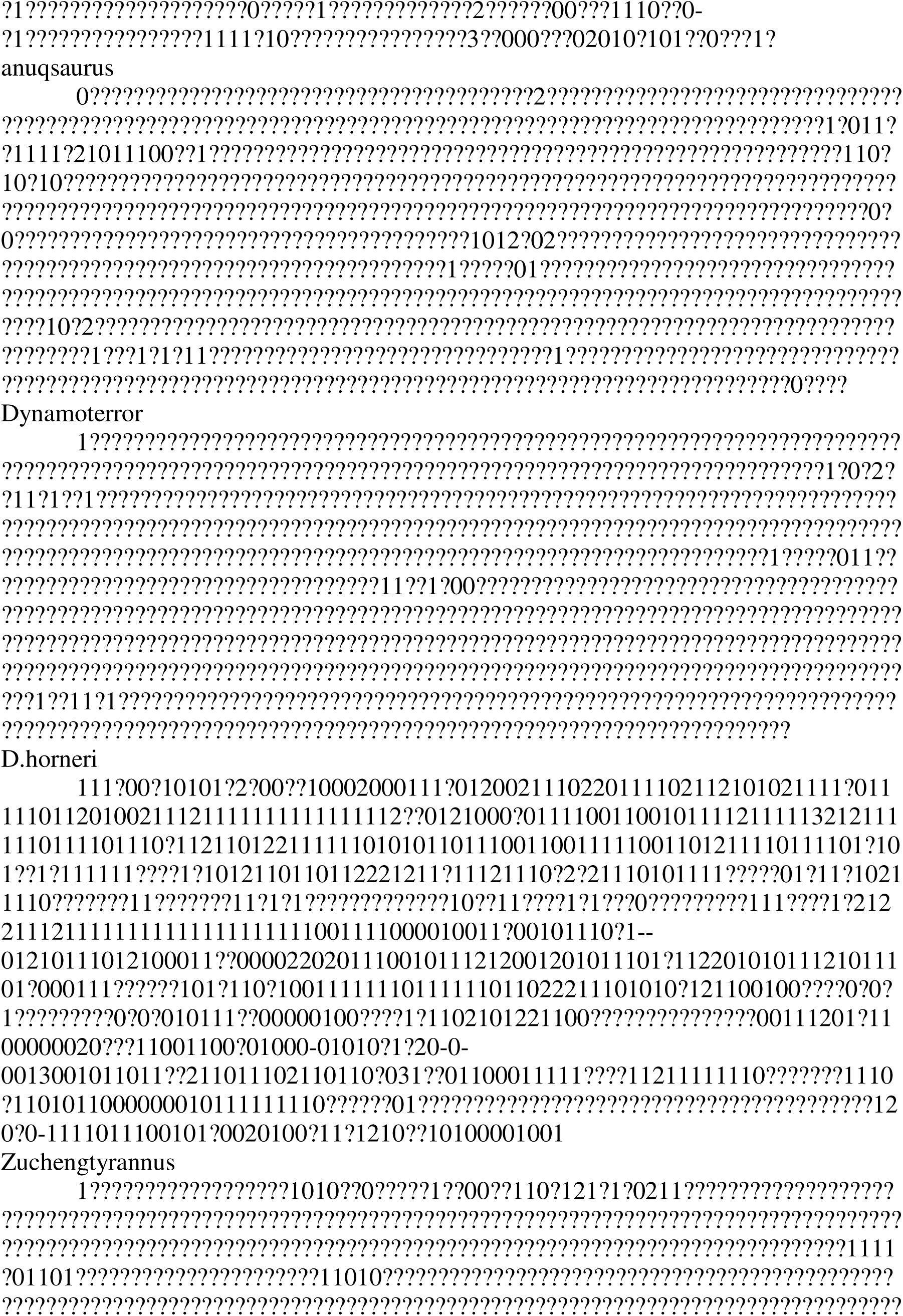
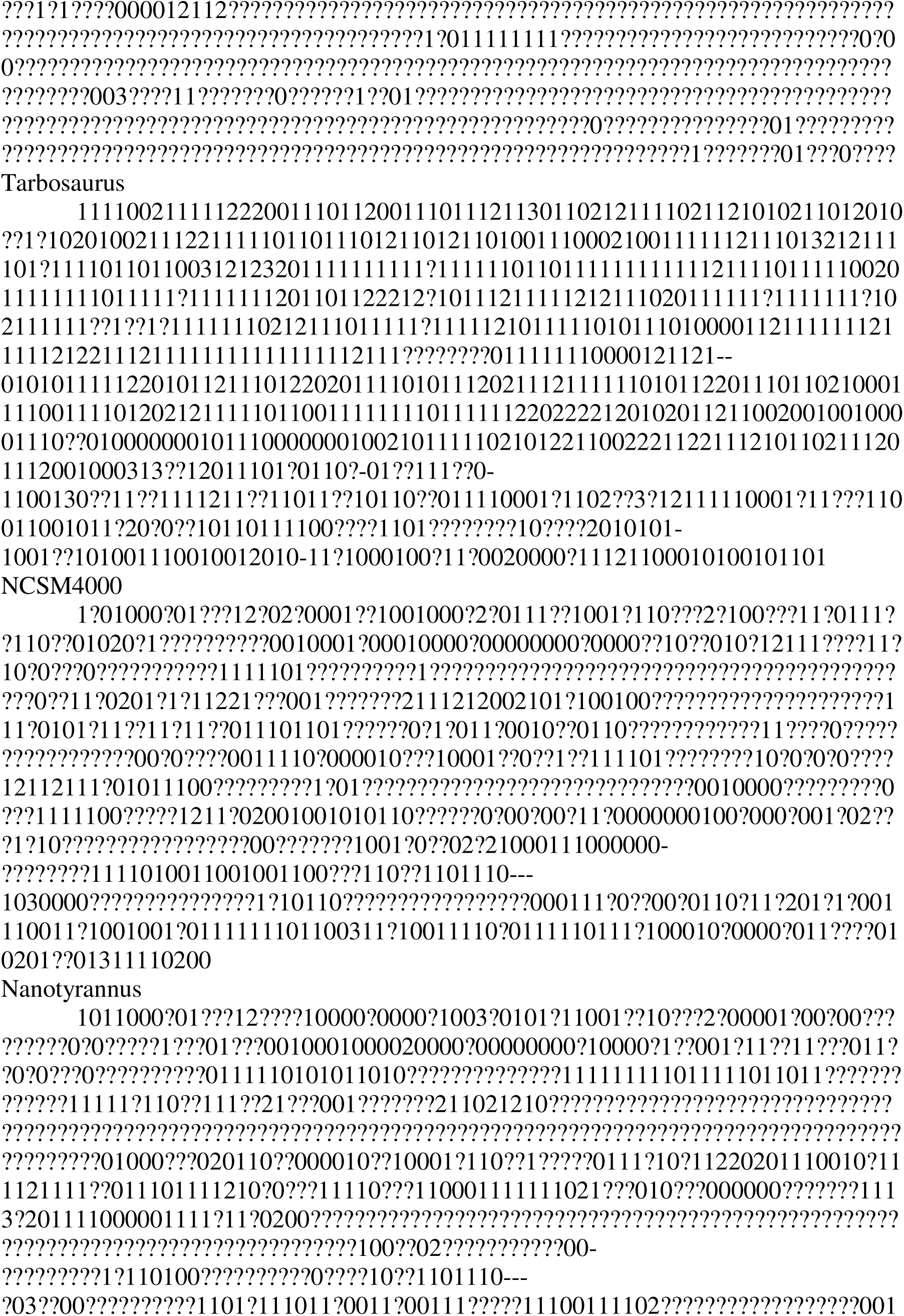
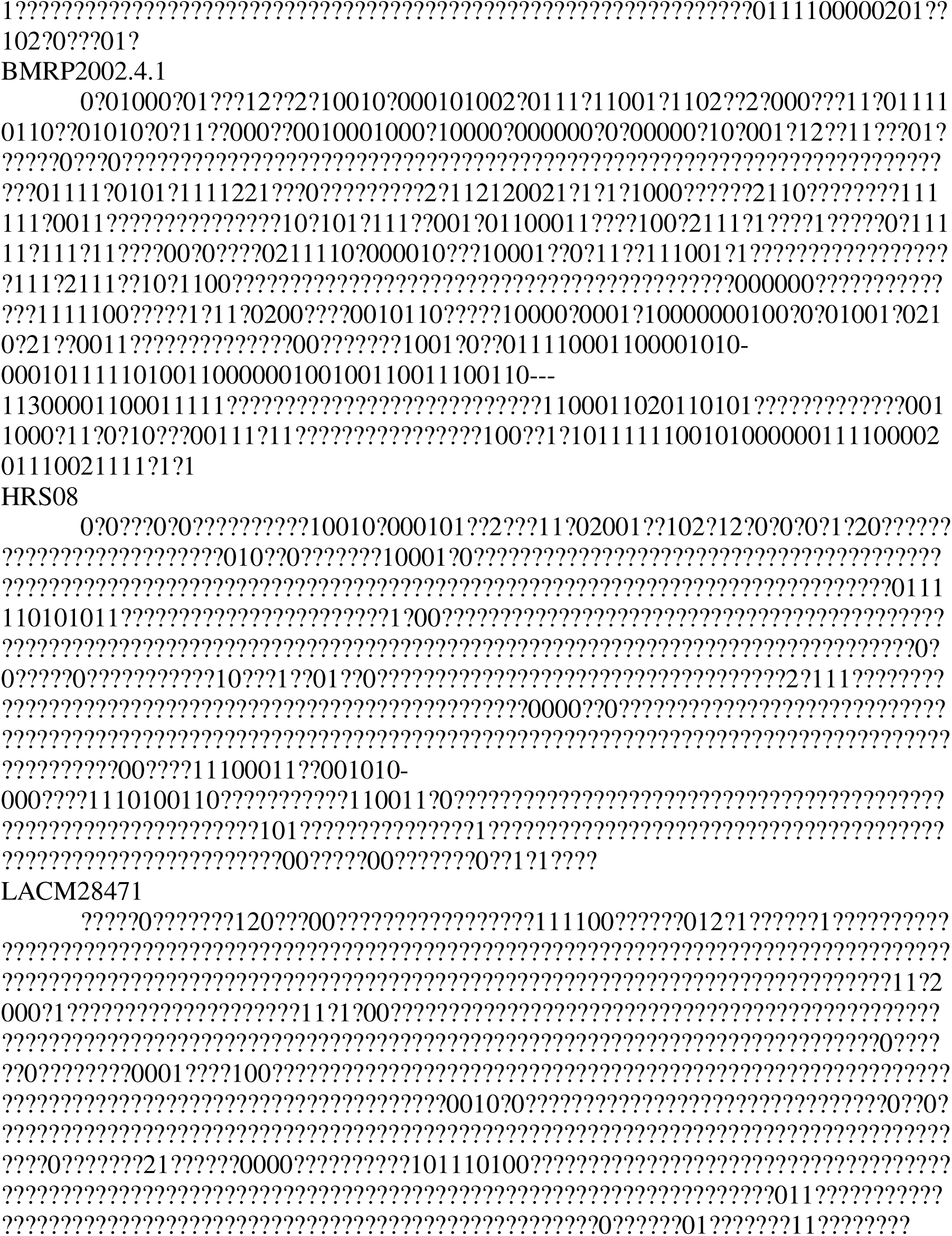

## Notes

### Competing Interest Statement

The authors have declared no competing interest.

### Summary of Updates

did some minor correction on grammar and general info

